# PRMT1 Inhibition Selectively Targets BNC1-Dependent Proliferation, but not Migration in Squamous Cell Carcinoma

**DOI:** 10.1101/2023.03.27.533164

**Authors:** Rafik Boudra, Bethany L. Patenall, Sandra King, Diana Wang, Sarah A. Best, Joo Yeon Ko, Shuyun Xu, Maria G. Padilla, Chrysalyne D. Schmults, Steven R. Barthel, Christine G. Lian, Matthew R. Ramsey

**Author notes:** Correspondence: Matthew R. Ramsey Brigham and Women’s Hospital 4 Blackfan Circle, Room 668 Boston, MA 02115.

## Abstract

Squamous Cell Carcinoma (SCC) develops in stratified epithelial tissues and demonstrates frequent alterations in transcriptional regulators. We sought to discover SCC-specific transcriptional programs and identified the transcription factor Basonuclin 1 (BNC1) as highly expressed in SCC compared to other tumor types. RNA-seq and ChIP-seq analysis identified pro-proliferative genes activated by BNC1 in SCC cells and keratinocytes. Inhibition of BNC1 in SCC cells suppressed proliferation and increased migration via FRA1. In contrast, BNC1 reduction in keratinocytes caused differentiation, which was abrogated by IRF6 knockdown, leading to increased migration. Protein interactome analysis identified PRMT1 as a co-activator of BNC1-dependent proliferative genes. Inhibition of PRMT1 resulted in a dose-dependent reduction in SCC cell proliferation without increasing migration. Importantly, therapeutic inhibition of PRMT1 in SCC xenografts significantly reduced tumor size, resembling functional effects of BNC1 knockdown. Together, we identify BNC1-PRMT1 as an SCC-lineage specific transcriptional axis that promotes cancer growth, which can be therapeutically targeted to inhibit SCC tumorigenesis.

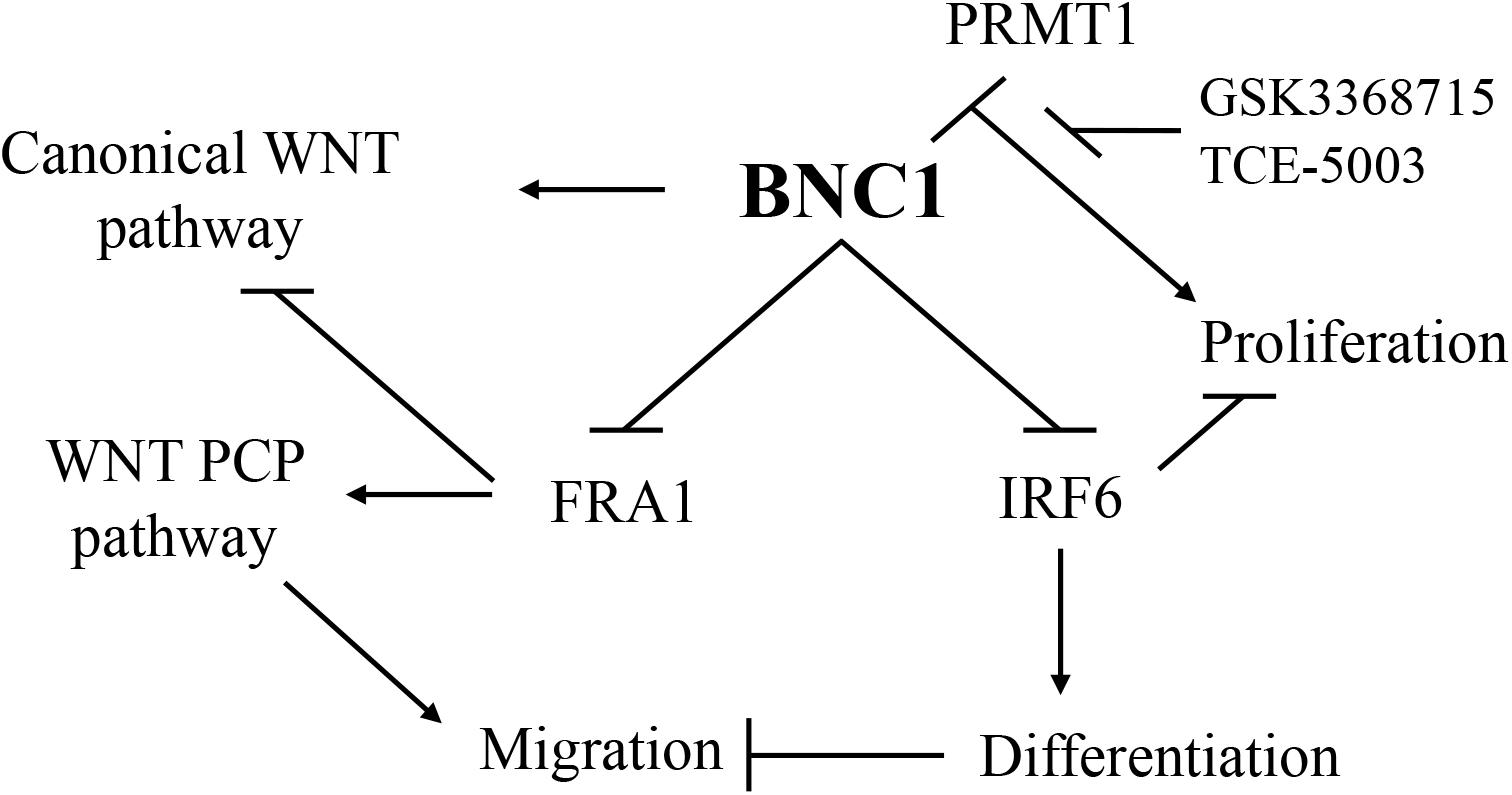

## Introduction

Squamous Cell Carcinoma (SCC) is a common form of cancer which develops most frequently in the skin, but can also arise in other stratified epithelial tissues such as the oral cavity and lungs, leading to greater than 50,000 deaths per year in all sites combined^1-3^. Once SCC becomes invasive, current therapies which include radiation, cisplatin, 5-fluorouracil, and paclitaxel, are only modestly effective and have many toxic side effects^4,5^. Recent advances in immunotherapy such as checkpoint receptor inhibition have improved clinical response rates^6,7^. Nonetheless, many patients with advanced disease do not respond, and overall survival gains have been modest, thus highlighting the need to both better understand SCC progression and identify new therapeutic targets for SCC patients.

Cancer cells are characterized by abnormal gene expression profiles when compared with their normal counterparts, which creates unique transcriptional dependencies^8^. This can result in broadly expressed transcription factors, such as c-MYC, creating oncogene dependence only in the tumor, allowing for the potential of therapeutic targeting without causing systemic side effects^9^. Similar to other malignancies, SCC tumors frequently have mutations in the transcription factor *TP53*^10^ and amplifications of the *MYC* oncogene^11^. There is also a high frequency of transcription factor alterations in SCC tumors, such as amplification of *TP63* and *SOX2*, deletions of *NOTCH1* and *ZNF750*, and mutations of *ZNF750*, *IRF6*, and *JUNB*^12^. Consistent with this mutation data, integrated TCGA data analysis reveals that distinct tumor types share gene expression patterns overlapping with their originating tissues^13^. Many of these frequently altered transcription factors control the normal differentiation processes such as P63, which is required for the commitment to epithelial lineage and for establishing proper epithelial stratification^14,15^. The activity of P63 is opposed by IRF6 and NOTCH1, which promote keratinocyte differentiation^16-19^. In lung epithelium, SOX2 coordinates with P63 and is required for proper trachea development and maintenance of basal cells^20,21^. Interestingly, P63 and SOX2 physically interact and are each required for tumor maintenance^22-24^, highlighting the interplay between transcriptional networks in SCC. In this study, we sought to dissect the novel functional interactions between transcription factors that arise in SCC which drive different stages of tumorigenesis. We have identified a transcription factor network centered on Basonuclin 1 (BNC1), which functions to toggle cells between a proliferative state and FRA1-dependent migration or IRF6-dependent differentiation. These functional states can be selectively targeted through the inhibition of PRMT1, which promotes proliferation, but does not affect migration, offering new therapeutic opportunities in SCC.

## Results

### Basonuclin 1 is Highly Expressed in Keratinocyte-Derived Cancers

To identify lineage-specific transcription factors important for SCC development and progression, we took advantage of the International Genomics Consortium Expression Project for Oncology^25^ data set, which contains expression data for 95 primary and metastatic SCC tumors from different organ sites (Lung, Cervix, Vulva, Tongue, Skin, Bladder, Breast, and Esophagus), and 1874 tumors of non-SCC histology (Adenocarcinoma, Clear Cell Carcinoma, Sarcoma, and others). Endometrioid tumors (N=158) and mixed adeno-squamous tumors (N=13) found in this cohort were excluded from analysis due to heterogeneous tumor histology (**Supplemental Table S1**). We sought to identify transcription factors whose expression is upregulated specifically in SCC versus other tumor types and compared gene expression profiles of SCC tumors to all other tumor types using the NCBI GEO2R tool. Differentially expressed genes were rank-ordered according to their Benjamini & Hochberg adjusted p-values (**Supplemental Table S2**). Many of the top 100 genes identified that distinguish SCC from other tumors were specific to stratified epithelial cells, such as keratins (*KRT6A, KRT6B, KRT5, KRT16, KRT13*), and regulators of epidermal differentiation (*SPRR1A, SPRR3, IVL*), validating this approach. Two well studied transcription factors, *TP63* (p= 5.60E-144) and *SOX2* (p= 7.83E-60) were both significantly and specifically overexpressed in SCC tumors in comparison to all other malignancies analyzed (**Figure 1A**) and represent established key lineage-dependent transcription factors essential for SCC maintenance^23,24^. Interestingly, the second most significantly enriched transcription factor (p=5.85E-83) in SCC cells compared to other tumor types was BNC1 (**Figure 1A**), which is expressed in the basal cells of stratified epithelium and is associated with proliferative capacity in keratinocytes^26,27^.

**Figure 1:**
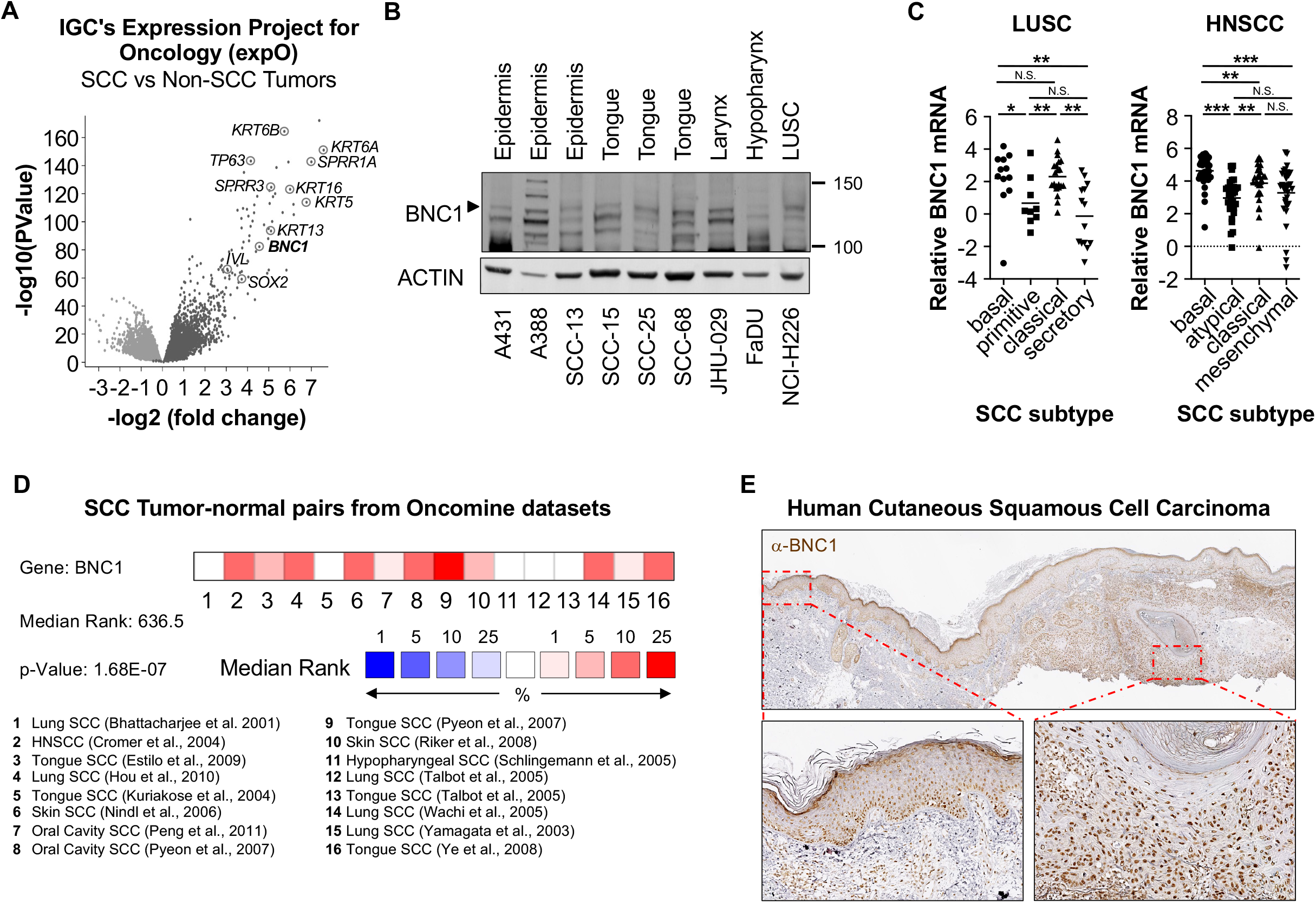
Identification of Basonuclin 1 as a lineage-specific transcription factor in SCC. **A.** Volcano plot of differentially expressed genes in Human primary SCC tumors (N=95) comparted to non-SCC tumors (N=1874) from IGC’s Expression Project for Oncology. **B.** Western blot analysis of BNC1 expression in SCC cell lines from indicated anatomic sites. Arrow indicates BNC1. ACTIN serves as a loading control. **C.** Relative mRNA expression of BNC1 in sqNSCLC tumors^31^ and HNSCC tumors^32^ of indicated subtype. **D.** Outlier analysis of BNC1 mRNA expression in 16 data sets from Oncomine comparing Human SCC to corresponding normal tissue. **E.** Immunohistochemical staining of BNC1 in Human cutaneous SCC (right) and adjacent normal skin (left).

We have found that commercially available BNC1 antibodies lack specificity, so we proceeded to generate a new antibody (BW175) that recognizes an epitope in the N-terminus of the protein shared by both human (A.A. 267-283) and murine (A.A. 266-282) BNC1 (**Supplemental Figure S1A**). Notably, antibody BW175 antibody reacted with endogenous BNC1 in human keratinocytes and SCC cells (**Supplemental Figure S1B**), while no protein was detected in human fibroblasts. As previously reported^28^, BNC1 localization was predominantly nuclear (**Supplemental Figure S1C**), and specificity was further validated by robust detection of over-expressed V5-epitope tagged BNC1 in 293T cells (**Supplemental Figure S1dD**) and markedly reduced detection following doxycycline-inducible shRNA-mediated knockdown of BNC1 using two independent hairpins in immortalized human keratinocytes (**Supplemental Figure S1E**). Immunohistochemical (IHC) staining of human skin with BW175 demonstrates specific expression of BNC1 in the epidermis and hair follicle (**Supplemental Figure S1F**), consistent with previous reports^29,30^. In addition, BW175 recognized BNC1 in human SCC cell lines (**Supplemental Figure. S1G**) and xenografts (**Supplemental Figure S1H**), as well as in murine SCC tumors (**Supplemental Figure S1I**). Examination of SCC cell lines from different anatomical sites demonstrated that BNC1 was expressed in lines derived from the epidermis, oral cavity, larynx, hypopharynx, and the lung (**Figure 1B**). To further characterize the patterns of *BNC1* expression in primary SCC tumors, we compared mRNA levels in previously defined molecular subtypes of Lung Squamous Cell Carcinoma (LUSC: basal, primitive, classical, secretory)^31^ and Head and Neck Squamous Cell Carcinoma (HNSCC: basal, atypical, classical, mesenchymal)^32^. In LUSC, the basal and classical groups had significantly higher BNC1 levels than the primitive and secretory groups, while the classical subtype was not significantly different from the basal group (**Figure 1C**). A similar pattern was seen in HNSCC (**Figure 1C**), where the basal group had the highest expression, while the atypical and mesenchymal groups had significantly lower *BNC1* mRNA levels. Interestingly, the basal subtype in both LUSC and HNSCC demonstrate similar global gene expression patterns as basal epithelial cells^31,32^, where BNC1 expression is normally high^33^ (**Supplemental Figure S1F**). In addition, comparison of *BNC1* levels in 16 different datasets from the Oncomine database^34^ comparing matched human SCC tumor-normal pairs found that *BNC1* is significantly overexpressed in SCC tumors (**Figure 1D**). Finally, we performed immunohistochemical analysis of human primary cutaneous SCC tumors, and found that in adjacent normal skin, BNC1 levels were high in the basal epithelial layer, and lower in more differentiated super-basal cells (**Figure 1E, left**). Interestingly, BNC1 levels in tumor cells were generally elevated in comparison to normal skin, with a decrease in expression in more differentiated regions around keratin pearls (**Figure 1E, right**). In total, these data identify Basonuclin 1 as a highly expressed lineage-specific transcription factor in keratinocyte-derived SCC tumors.

### BNC1 directly regulates proliferation in both keratinocytes and SCC cells

In prior studies, BNC1 was found to binds to and regulates ribosomal RNA genes^35,36^, but dissection of the complete BNC1-regulated transcriptome in epithelial and SCC cells has not been performed. We knocked down BNC1 expression in human keratinocytes (N/TERT-1, OKF6/TERT) and SCC cells (SCC-13, SCC-68) using shRNAs and then systematically characterized BNC1 downstream targets by RNA-seq analyses. Differential expression of transcripts between control scrambled shRNA cells (n=3 per line) and cells with shRNA directed against BNC1 (n=3 per line) was assessed following DESeq2 normalization. In keratinocytes, 5307 transcripts were significantly different with greater than 1.5-fold change between control and BNC1 knock-down cells (**Figure 2A, Supplemental Table S3**), while analysis of SCC cells identified 2674 significantly different transcripts (**Figure 2B, Supplemental Table S4**). To determine which of these transcripts were directly regulated by BNC1, we performed ChIP-seq in keratinocytes (**Supplemental Table S5**) and SCC cells (**Supplemental Table S6**) using the BW175 antibody, which specifically immunoprecipitated BNC1 and enriches for known BNC1 binding sites^35,37^ as demonstrated by ChIP-QRT-PCR (**Supplemental Figures S2A-C**), thus validating this antibody’s utility. BNC1 binding was enriched in promoter regions of target genes, and similar distribution of BNC1 binding around gene bodies (**Figure 2C**) and across the genome (**Figure 2D**) was seen in both keratinocytes and SCC cells. Motif enrichment analysis using HOMER^38^ identified a consensus binding sequence for BNC1 associated with ZBTB33/Kaiso, a methylation-dependent regulator of Non-canonical WNT signaling^39^ in both keratinocytes and SCC cells (**Figure 2E, Supplemental Table S7**). BNC1 binding was also enriched at motifs for P63, AP2-α, and AP-2γ which are known to regulate epidermal development^40^. Interestingly, BNC1 binding sites were enriched for AP-1 motifs (**Figure 2E**), and AP-1 transcription factors known to regulate re-epithelialization after wounding^41-44^.

**Figure 2:**
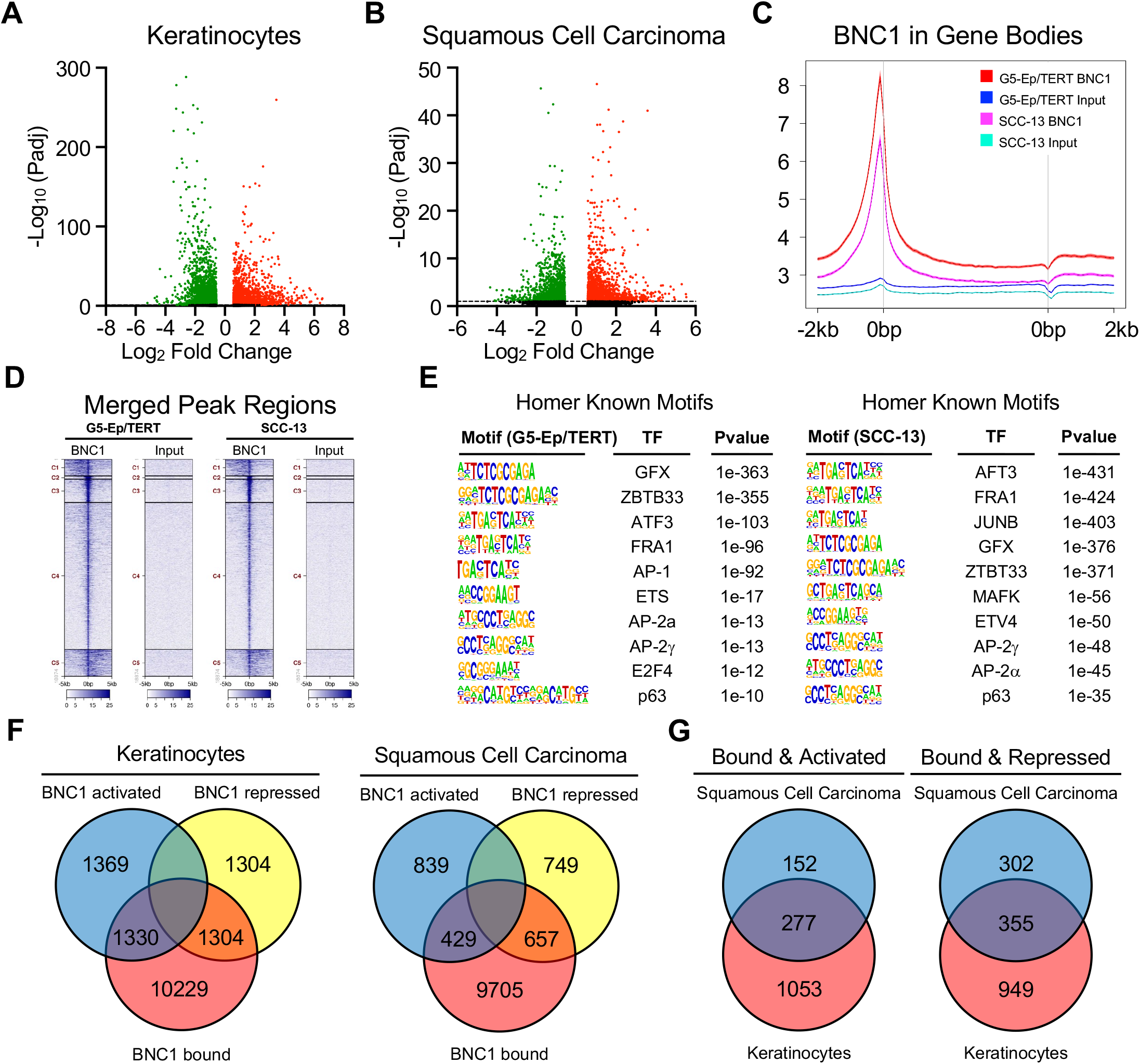
Analysis of BNC1 target genes in human keratinocytes and SCC cells. **A.** Volcano plot of differentially expressed genes in human keratinocytes (N/TERT-1, OKF6/TERT) following induction of shRNA targeting BNC1 assessed by RNA-seq. N= 3 replicates per shRNA in each cell line. **B.** Volcano plot of differentially expressed genes in human SCC cells (SCC-13, SCC-68) following induction of shRNA targeting BNC1 assessed by RNA-seq. N= 3 replicates per shRNA in each cell line. **C.** Average distribution of BNC1 near gene bodies in keratinocytes (G5-Ep/TERT) and SCC cells (SCC-13) assessed by ChIP-seq. **D.** Clustered distribution of merged peak regions bound by BNC1 in keratinocytes and SCC cells from ChIP-seq analysis. **E.** HOMER known motif enrichment analysis of BNC1 bound regions in keratinocytes (left) and SCC cells (right). **F.** Venn diagram indicating overlap between genes bound by BNC1 and genes significantly regulated by BNC1 in keratinocytes (left) and SCC cells (right) **G.** Venn diagram indicating overlap between genes bound and regulated by BNC1 in both keratinocytes and SCC cells

In order to identify cellular processes directly regulated by BNC1, we performed gene enrichment analysis of BNC1-bound and regulated genes (**Figures 2F, G**) using the ENRICHR portal^45^. For genes shared by both keratinocytes and SCC cells (**Figure 2G**), there was significant enrichment for terms associated with proliferation, such as “DNA metabolic process“, “Cell cycle”, “Mitotic Prometaphase”, and “DNA strand elongation”(**Supplemental Table S8**). Previous data in keratinocytes has suggested BNC1 is associated with proliferative capacity^26,27,29,46^, which could be potentially regulated by BNC1’s role in promoting ribosomal biogenesis^35,36^. We found that shRNA-mediated knock-down of BNC1 resulted in a large increase in the cell cycle inhibitor *CDKN1A* and significant reductions in key S-phase genes (*MCM2, GINS4, RFC4, RFC5, LIG1, PRIM1*) and M-phase genes (*CCNB1, CDC20, SMC2, SMC4, SPC25, DSN1*) in both keratinocytes (**Figure 3A**) and SCC cells (**Figure 3B**). To test if BNC1 was required for proliferation, we performed shRNA-mediated knock-down of BNC1 followed by colony forming assays. In SCC cells (SCC-13, JHU-029, SCC-68), primary keratinocytes (G5-Ep), and immortalized keratinocytes (N/TERT-1, OKF6/TERT) there was a significant reduction in colony formation following BNC1 knock-down with 2 independent shRNAs (**Figure 3C, Supplemental Figure S3A**). This was accompanied by a significant reduction in BrdU incorporation, as assessed by flow cytometry (**Figure 3D, Supplemental Figure S3B**), indicating decreased DNA synthesis. To examine if BNC1 is required for tumor growth *in vivo*, we generated xenografts using SCC-13 and JHU-029 cells. Mice were treated with Doxycycline after all mice had visible tumors to induce shRNA expression and to avoid potential earlier effects on tumor engraftment. Reduction of BNC1 (**Supplemental Figure S3C**) caused a significant reduction in tumor volume (**Figure 3E**), which was associated with significantly reduced Ki67 staining assessed by IHC (**Figure 3F**). These data demonstrate that BNC1 controls proliferation through direct transcriptional regulation of genes required for both DNA synthesis and mitosis.

**Figure 3:**
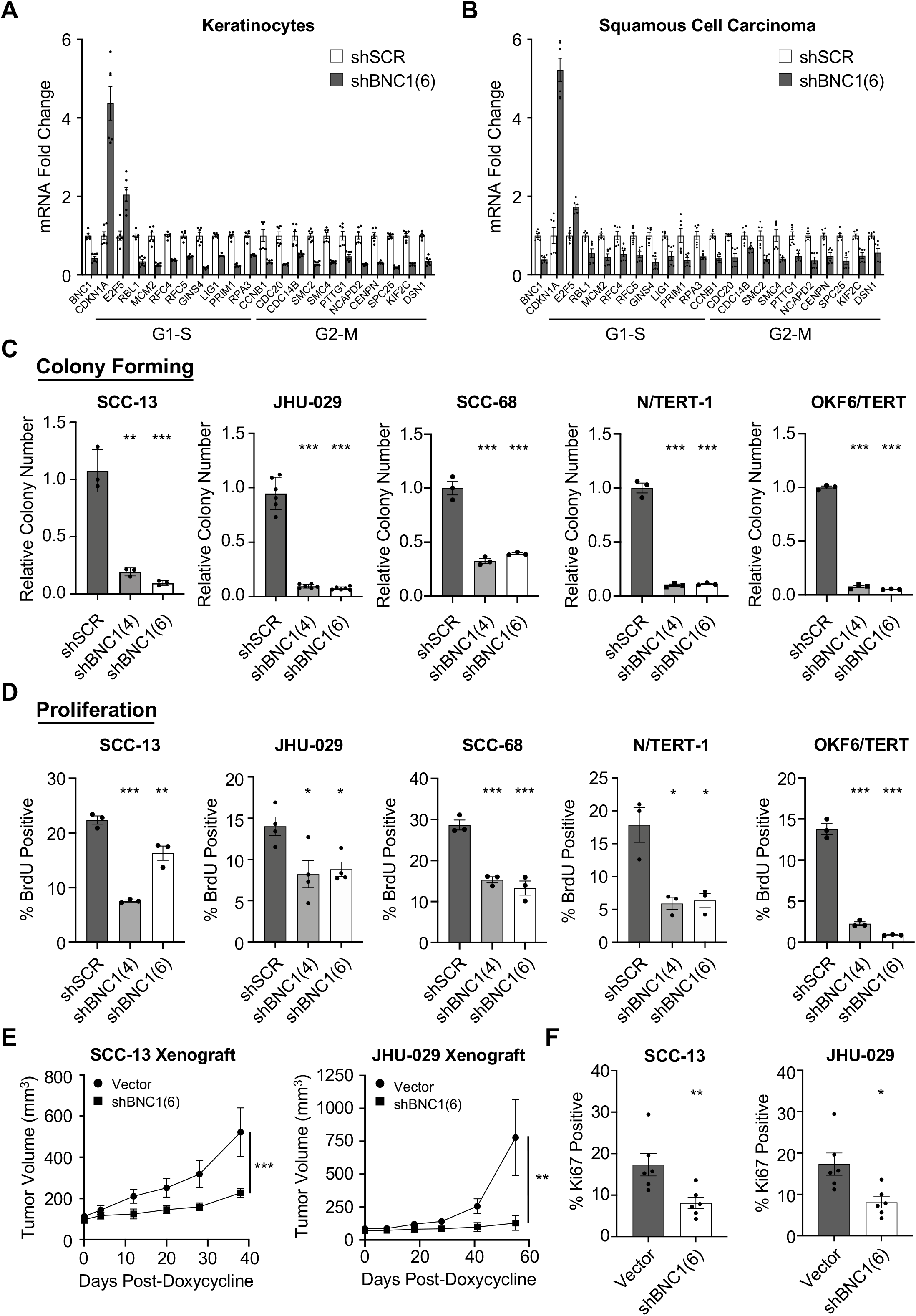
BNC1 promotes proliferation in keratinocytes and SCC cells. **A.** Significantly changed genes associated with proliferation in keratinocytes following BNC1 knock-down as assessed by RNA-seq. N= 3 replicates per shRNA in each cell line. **B.** Significantly changed genes associated with proliferation in SCC cells following BNC1 knock-down as assessed by RNA-seq. N= 3 replicates per shRNA in each cell line. **C.** Colony forming assays of indicated cell lines 2 weeks after induction of indicated shRNA construct. Quantification (right) of colony number by OD595 of extracted crystal violet stain (n=3-6 wells). **p<0.01, ***p<0.001. Error bars +/- S.E.M. **D.** FACS analysis of BrdU positive cells in cell lines with indicated shRNA construct after a 2-hour pulse of BrdU. Quantification (right) of BrdU positive cells (n=3-4). *p<0.05, **p<0.01, ***p<0.001. Error bars +/- S.E.M. **E.** Growth of SCC-13 (left, n=10 per group) and JHU-029 (right, n= 6 per group) Xenograft tumors with indicated shRNA construct. **p<0.01, ***p<0.001. as assessed by Multiple measures ANOVA. Error bars +/- S.E.M. **F.** Quantification of Ki67 IHC staining in SCC-13 (left) and JHU-029 (right) Xenograft tumors. *p<0.05, **p<0.01, Error bars +/- S.E.M.

### BNC1 cooperates with FRA1 to regulate SCC cell migration

In addition to the clear effects on proliferation, we noted significant enrichment for terms such as “regulation of epithelial cell migration,” “signaling by Rho GTPases”, and “cell-substrate junction assembly” in SCC cells suggesting a potential role for BNC1 in regulating migration (**Supplemental Table S8**). We performed shRNA-based knockdown of BNC1 followed by *in vitro* scratch assay analysis and found a significant increase in migration in SCC cells after BNC1 reduction (**Figure 4A**). We sought to understand the mechanisms regulating migration in SCC cells and noted enrichment for AP-1 motifs in our ChIP-seq analysis, particularly FRA1 and JUNB (**Figure 2E**). Interestingly, reduction of either JUNB or FRA1 has been shown to reduce migration of SCC cells^47,48^ and overexpressed FRA1 can promote migration of both SCC cells^48^ and immortalized keratinocytes^49^. We performed ChIP-seq of FRA1 and JUNB in SCC cells and found high overlap with BNC1 binding, although FRA1 and JUNB were less enriched at promoter regions than BNC1 (**Figure 4B**, **Supplemental Figure S4A, and Supplemental Table S9**). There was an highly significant correlation (Spearman = 0.947) between FRA1 and JUNB binding peaks across the genome (**Supplemental Figure S4B**), and virtually all binding peaks identified for FRA1 were also bound by JUNB (**Figure 4C**) consistent with FRA1-JUNB heterodimerization. However, only 1/3 of the total JUNB sites were also bound by FRA1 (**Figure 4C**), suggesting that JUNB has other AP1 dimerization partners (**Supplemental Figure S4C, Supplemental Table S10**). Approximately half of all BNC1 peaks co-localized with both JUNB and FRA1 (**Figure 4C, Supplemental Figure S4D**), suggesting that a major part of BNC1’s function is mediated through JUNB-FRA1 dimers. We hypothesized that the pro-migratory functions of BNC1 require FRA1. To examine the functional requirement for FRA1 in BNC1-dependent migration, we performed scratch assays following BNC1, FRA1, or FRA1 and BNC1 knock-down. We found that reduction of FRA1 alone abrogated migration and was required for the increased migration seen following BNC1 knock-down (**Figure 4D**).

**Figure 4:**
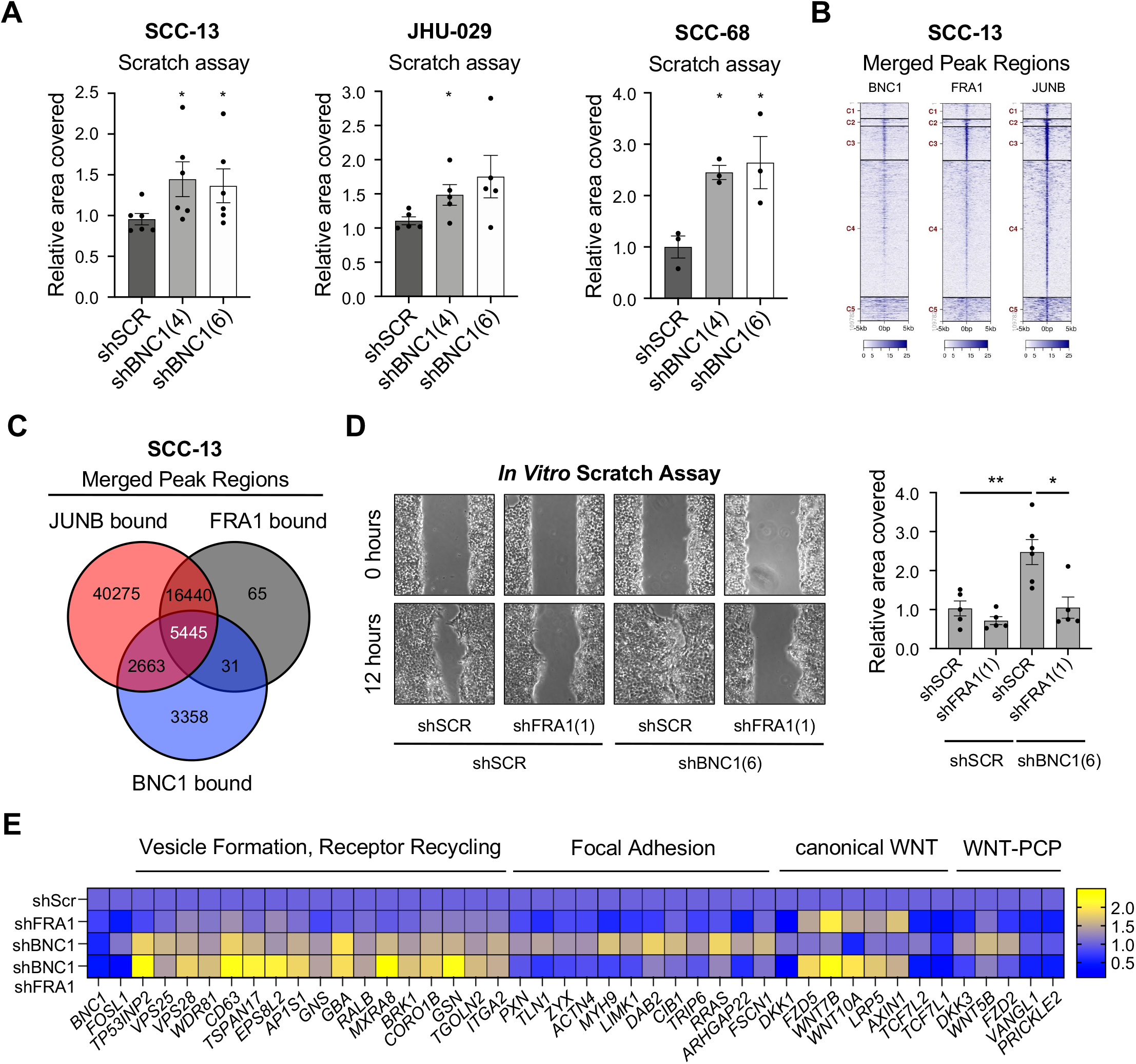
BNC1 and FRA1 co-regulate migration in SCC cells. **A.** Quantification of *In vitro* scratch assays in SCC cells at 6-8 hours scratch. Cells were treated with doxycycline 96 hours prior to scratch initiation to induce expression of indicated shRNA. Quantification of area covered by cells relative to shScr control. (n=3-6) *p<0.05, **p<0.01. Error bars indicate +/- S.E.M. **B.** Clustered distribution of merged peak regions bound by BNC1, FRA1, and JUNB in SCC-13 cells from ChIP-seq analysis. **C.** Venn diagram of shared merged peak regions bound by BNC1, FRA1, and JUNB in SCC-13 cells from ChIP-seq analysis. **D.** *In vitro* scratch assays of SCC-13 cells 6 hours following scratch. Cells were treated with doxycycline 96 hours prior to scratch initiation to induce expression of indicated shRNA. Quantification (right) of area covered by cells relative to shScr control. (n=5) *p<0.05, **p<0.01. Error bars indicate +/- S.E.M. **E.** Heat Map of gene expression in SCC-13 cells following shRNA-mediated knock-down of indicated gene assessed by RNA-seq. N= 3 replicates per shRNA.

To dissect the transcriptional programs controlling migration, we performed RNA-seq on SCC-13 cells following knockdown of BNC1, FRA1, or both BNC1 and FRA1. Independently of FRA1, reduction of BNC1 caused an increase in genes associated with vesicle formation (*VPS25*, *VPS28*, *WDR81*, *TGOLN2*, *CD63*, *AP1S1*) and trafficking (*BRK1*, *CORO1B*, *GSN*), consistent with dynamic restructuring of cell-cell and cell-ECM interactions^50,51^. Migrating cells require the re-organization of focal adhesions, and BNC1 knock-down caused an increase in focal adhesion proteins (*PXN*, *TLN1*), filament bundle proteins (*ZYX*, *ACTN4*, *MYH9*, *VIM*), and signaling adaptor proteins (*TRIP6*, *CIB1*, *DAB2*, *RRAS*, *ARHGAP22*), dependent on expression of FRA1 (**Figure 4E**). We also observed an increase in members of the canonical WNT/β-catenin signaling pathway (*FXD5*, *WNT7B*, *WNT10A*, *LRP5*) with reduction of FRA1, while loss of BNC1 increased expression of members of the migration-associated^52^ WNT-Planar Cell Polarity Pathway (*WNT5B*, *FZD2*, *VANGL1*, *PRICKLE2*), which required the presence of FRA1. In total, these results demonstrate that BNC1 suppresses a FRA1-dependent pro-migratory program in SCC.

### A BNC1-IRF6 regulatory loop regulates keratinocyte differentiation

Previous studies have reported a role for murine Bnc1 in epithelial wound repair, a phenotype attributed to changes in proliferation due to altered transcription of ribosomal DNA^53^. Given our data in SCC cells we first sought to determine if BNC1 levels changed in keratinocytes when cells migrate. We performed *in vitro* scratch assays followed by western blotting and found that in both SCC cells and keratinocytes, BNC1 protein levels were decreased in migrating cells (**Figure 5A**). In addition, knock-down of BNC1 resulted in phosphorylation of p130 (Y165) and FAK (Y397) (**Figure 5B**) which promotes turnover of focal adhesions near the leading edge of migrating cells^54,55^. We then sought to determine if BNC1 levels changed during epidermal wound healing. In humans, the re-epithelialization process involves migration across the wound bed, while in mice contraction of the skin is a major mechanism of closure^56^. We employed an epidermal splinting system in which a silicone ring is glued and sutured to the skin around the wound (**Figure 5C**), which stops the contraction and forces keratinocytes to migrate to close a wound^57^. Immunohistochemical staining seven days after wounding showed that while there was high Bnc1 expression away from the wound (**Figure 5D**, left), leading edge keratinocytes had marked down-regulation of Bnc1 protein (**Figure 5D,** right). Immunofluorescent staining with pan-cytokeratin antibodies confirmed that these cells lacking Bnc1 were keratinocytes (**Supplemental Figure S5A**). Surprisingly, following BNC1 knockdown, cultured keratinocytes did not migrate, but instead appeared to differentiate (**Figures 5E, 5F**). Immunofluorescent staining following BNC1 knockdown demonstrated that keratinocytes increased levels of Involucrin, a marker of terminal differentiation (**Supplemental Figure S5B**). Western blotting confirmed increased Involucrin and SPRR1B 48 hours after BNC1 knock-down in keratinocytes (**Figure 5G**).

**Figure 5:**
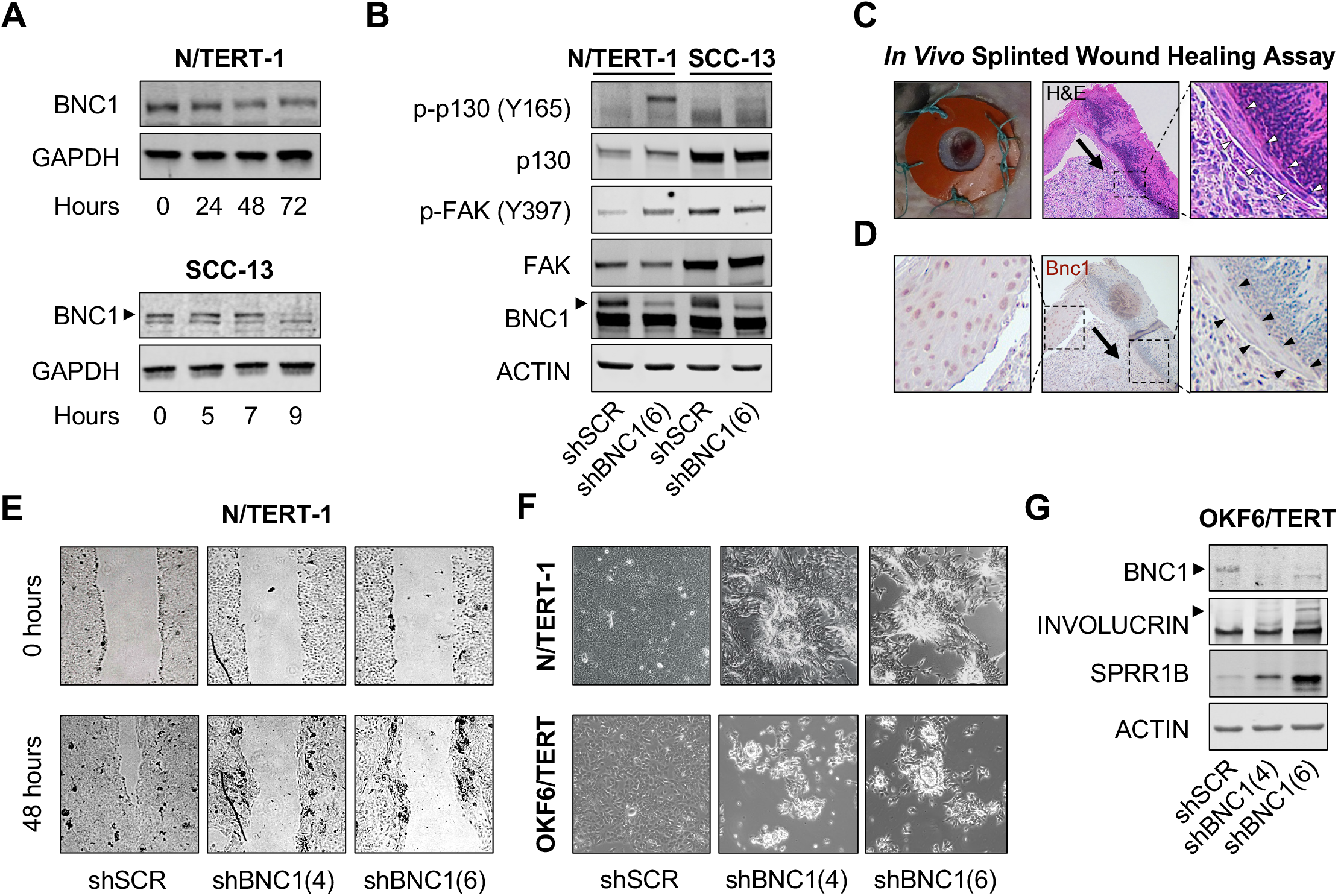
Loss of BNC1 in keratinocytes induces differentiation. **A.** Western blot analysis of BNC1 levels in keratinocytes (top) or SCC cells (bottom) at indicated time after initiation of an *in vitro* scratch assay. Arrow indicates BNC1. GAPDH serves as a loading control. **B.** Western blot analysis of adhesion signaling in keratinocytes and SCC cells following BNC1 knock-down. ACTIN serves as a loading control. **C.** Splinted *in vivo* wound healing assay to assess migration of keratinocytes. Hematoxylin and Eosin (H&E) stain and 7 days after wounding. Arrow indicates direction of migration. Arrowheads indicate leading edge keratinocytes. **D.** Bnc1 immunohistochemistry of splinted wound 7 days after wounding. Arrow indicates direction of migration. Arrowheads indicate leading edge keratinocytes. **E.** *In vitro* scratch assays in keratinocytes at initiation and 48 hours following scratch. Cells were treated with doxycycline 96 hours prior to scratch initiation to induce expression of indicated shRNA. **F.** Photomicrographs of keratinocytes 96 hours after treatment with doxycycline to induce indicated shRNA construct. **G.** Western blot analysis of differentiation markers in keratinocytes 96 hours after treatment with doxycycline to induce indicated shRNA construct. ACTIN serves as a loading control.

To identify potential regulators of this differentiation phenotype, we performed a pairwise comparison of directly repressed BNC1 target genes in keratinocytes with publicly available transcriptomic datasets using the ARCHS4 platform^58^. Multiple transcription factors involved in the regulation of epidermal differentiation, such as IRF6^17,59^, GRHL2^60^, and GRHL3^61^ were significantly enriched in this analysis (**Figure 6A, Supplemental Table S12)**. Interestingly, *GRHL3* is transcriptionally regulated by the IRF6 signaling axis^62^ and, IRF6 is required for epidermal differentiation^17^. Western blot analysis demonstrated that IRF6 levels were increased following BNC1 knock-down in keratinocytes, but not in SCC cells (**Figure 6B, Supplemental Figure S6A**). We performed ChIP-seq analysis of IRF6 binding in keratinocytes and found that BNC1 and IRF6 are bound in close proximity at many sites across the genome (**Figure 6C**) and identified 7561 genes with significantly enriched IRF6 binding (**Supplemental Table S13**). IRF6 binding was enriched in promoter regions (**Supplemental Figures S6B, C**), and there were far fewer binding peaks seen in SCC cells than in keratinocytes, consistent with the reported tumor suppressor function for IRF6 in SCC^63^. We hypothesized that reductions in BNC1 protein may cause the activation of an IRF6-dependent differentiation program in keratinocytes. While keratinocytes showed a pronounced differentiation phenotype after BNC1 knock-down, this was abrogated by additional knock-down of IRF6 (**Figure 6D**). Western blot analysis demonstrated that the differentiation markers SPRR1B and Filaggrin were increased in keratinocytes following BNC1 knock-down while knock-down of both BNC1 and IRF6 blocked induction of SPRR1B and Filaggrin (**Figure 6E**). To determine if BNC1 could block IRF6-dependent transcription, we performed a luciferase assay using a reporter control (pGL3-CMV-Luc) or IRF6 reporter (pGL3-IFNβ-Luc) in HEK293T cells transfected with expression vectors coding for BNC1, IRF6 or both. As expected, IRF6 expression activates the pGL3-IFNβ-Luc reporter, but co-expression with BNC1 completely abrogated this effect (**Figure S6D**), showing that BNC1 represses IRF6 transcriptional activity. We hypothesized that BNC1 could repress IRF6-dependent transcription of differentiation genes by direct binding in the promoter of target genes, blocking IRF6-dependent activation. We used ChIP-QRT-PCR to examine changes in IRF6 occupancy following BNC1 knock-down at sites bound by both BNC1 and IRF6 in the *SPRR1B* promoter (**Figure S6E**) and found that IRF6 remained bound at similar levels in both control and BNC1 knockdown cells (**Figure 6F**). Consistent with increased *SPRR1B* mRNA levels (**Figure S6F**), knockdown of BNC1 resulted in a significant increase in H3K9ac at the promoter (**Figure 6F**), indicating transcriptional activation. To examine the physiological relevance of this effect, we examined the leading-edge keratinocytes in our splinted wound healing model (**Figure 5C**) 7 days after wounding. Immunohistochemical staining for BNC1 and IRF6 demonstrated that both BNC1 and IRF6 are expressed in cells away from the leading edge, but migrating keratinocytes have a significant reduction in both proteins (**Figure 6G**). We then sought to determine if the IRF6-dependent differentiation effect was responsible for reduced migration in keratinocytes. While loss of BNC1 resulted in minimal migration of cells in scratch assays after 24 hours, combined loss of IRF6 and BNC1 rescued this migration defect (**Figure 6H**). This data supports a model where BNC1 prevents differentiation by co-occupying IRF6 sites at transcriptional control regions, suppressing IRF6-dependent transcriptional activity. Reduction of both BNC1 and IRF6 are therefore required for proper migration during physiological damage such as during epidermal wound healing.

**Figure 6:**
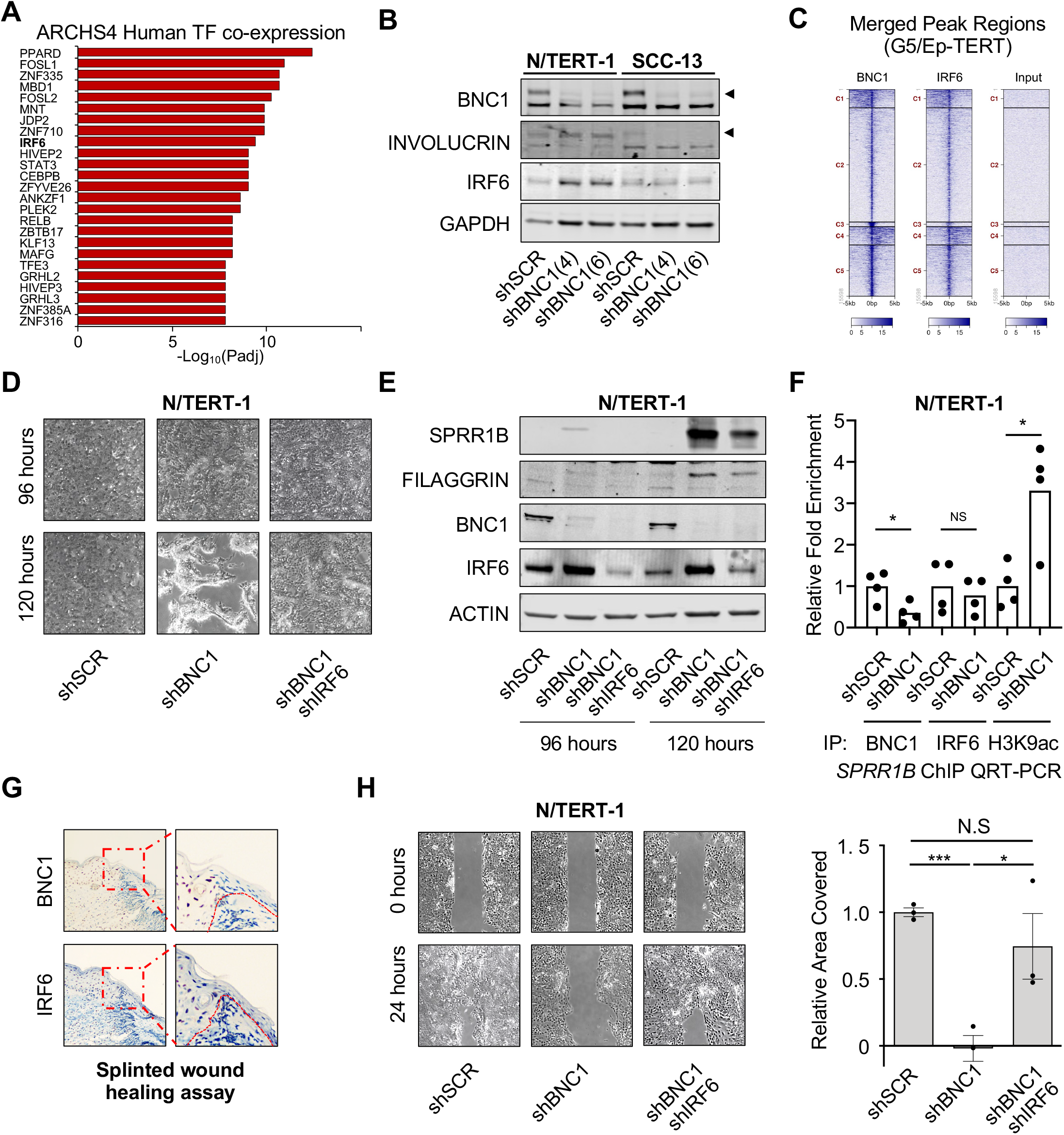
BNC1 regulates an IRF6-dependent transcriptional program. **A.** Top 25 most significant transcription factors pairwise comparison of direct BNC1 target genes regulated in primary keratinocytes with publicly available transcriptomic datasets using the ARCHS4 platform **B.** Western blot analysis of indicated proteins in keratinocytes (N/TERT-1) and SCC cells (SCC-13). Cells were treated with doxycycline 96 hours prior to analysis to induce expression of indicated shRNA. GAPDH serves as a loading control. **C.** Clustered distribution of merged peak regions bound by BNC1 and IRF6 in G5-Ep/TERT cells from ChIP-seq analysis. **D.** Genetic rescue of differentiation in keratinocytes 96 and 120 hours after induction of indicated shRNAs. **E.** Western blot analysis of differentiation markers in keratinocytes 96 and 120 hours after induction of indicated shRNAs. Actin serves as a loading control. **F.** ChIP-QRTPCR in N/TERT cells expressing the indicated shRNA of BNC1, IRF6 and H3K9ac enriched regions in the promoter of *SPRR1B*. *p<0.05, NS: not significant, Error bars indicate +/- SEM. **G.** Immunohistochemistry for BNC1 (left) and IRF6 (right) in leading edge keratinocytes 7 days after wounding in splint model. **H.** *In vitro* scratch assays in keratinocytes 24 hours following scratch. Cells were treated with doxycycline 96 hours prior to scratch initiation to induce expression of indicated shRNA. Quantification (right) of area covered by cells relative to shScr control. (n=3) *p<0.05, **p<0.01. Error bars indicate +/- S.E.M.

### PRMT1 is a therapeutically targetable BNC1 transcriptional co-activator

Direct therapeutic targeting of transcription factors has been challenging, but inhibition of co-factors, such as Histone Deacetylases offers a way to modulate the activity of transcriptional complexes^64^. Given the strong pro-tumorigenic function of BNC1-dependent activation of proliferation genes, we sought to identify co-factors essential for transcriptional activation. We performed RIME^65^ on N/TERT-1 keratinocytes using α-BNC1 antibodies to isolate BNC1-interacting nuclear proteins. Among the interacting proteins, we identified the transcription factors JUNB, IRF6, TP63, and TFAP2A (**Figure 7A**) consistent with the motif analysis of BNC1 bound regions following ChIP-seq (**Figure 2E**). Interestingly, we found SMARCA5 which mediates nucleosome spacing, influencing transcription factor binding^66^ and the transcriptional activator complex proteins PRMT1, CHTOP and ERH^67^ also bound to BNC1. PRMT1 is highly expressed in basal keratinocytes and a comparison between PRMT1-regulated genes^68^ and direct BNC1 target genes in keratinocytes showed significant overlap and enrichment for cell cycle genes (**Table S14**). We hypothesized that inhibition of the BNC1-PRMT1 transcriptional activator complex would lead to a block in proliferation, but not increase migration. SCC cells demonstrated a dose-dependent inhibition of growth following treatment with the PRMT1 inhibitor TCE-5003 and were more sensitive than immortalized keratinocyte lines (**Figure 7B**). Treatment of SCC cell lines with TCE-5003 resulted in a significant reduction in BrdU incorporation as assessed by flow cytometry (**Figure 7C**), but unlike BNC1 knockdown, we did not observe an increase in SCC cell migration (**Figure 7D**). To examine the therapeutic potential of PRMT1 inhibition in SCC, we generated Xenografts using JHU-029 cells, and mice were treated with the Type I PRMT inhibitor GSK3368715, which has previously been shown to inhibit growth in a variety of tumor types^69^, at 10mg/kg by intraperitoneal injection every other day. Importantly, we started treatment with GSK3368715 seven days after tumor cell injections to eliminate any effects on tumor engraftment and more accurately mirror clinical practice. After only 4 days of treatment, tumors in GSK3368715 treated mice were significantly smaller than vehicle treated mice (**Figure 7E**). In total, these experiments support a model by which BNC1-dependent activation of pro-proliferative target genes requires PRMT1, while BNC1-repressed pro-migratory genes are unaffected.

**Figure 7:**
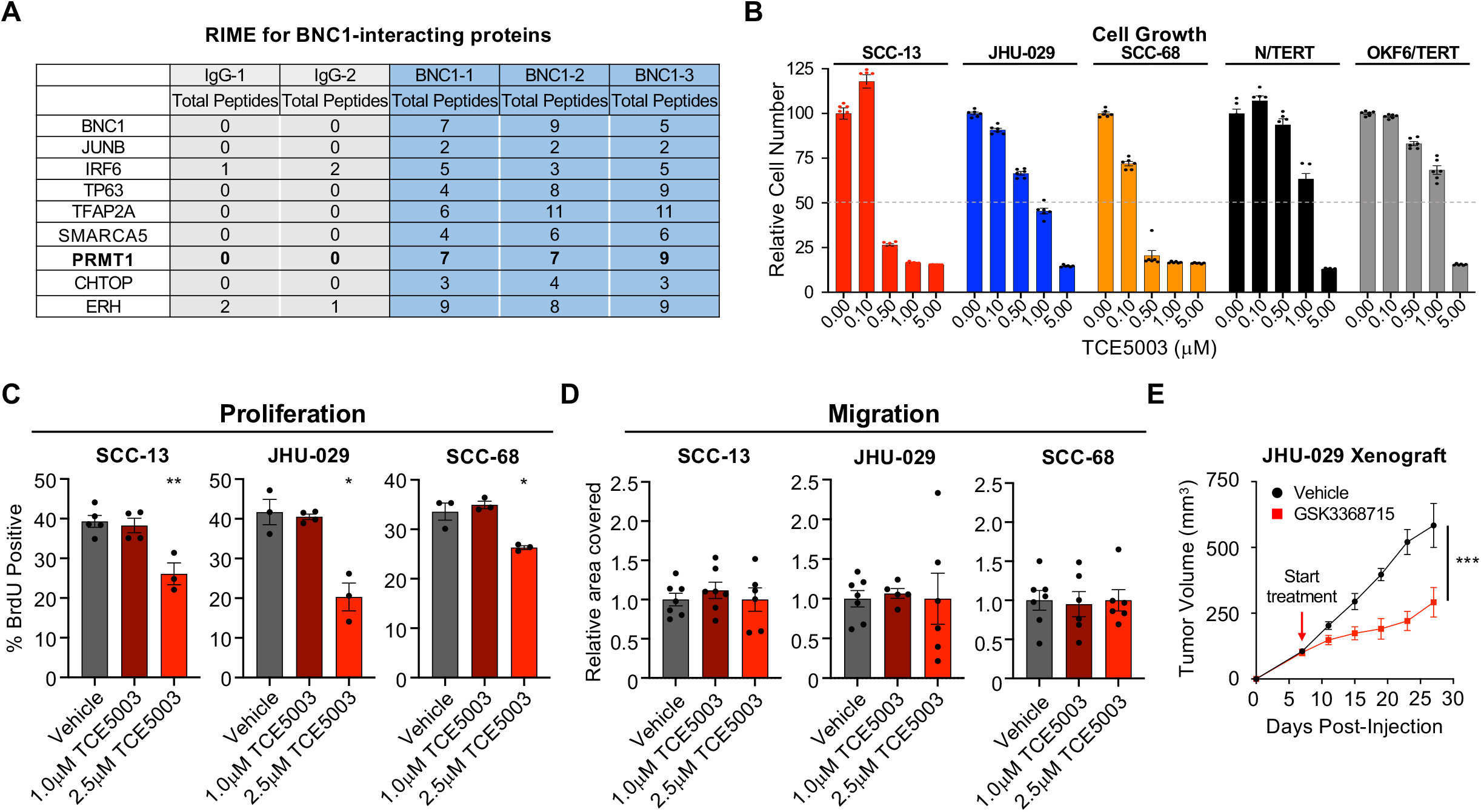
PRMT1 interacts with BNC1 to promote SCC proliferation. **A.** BNC1-interacting proteins identified by RIME. **B.** Cell growth of SCC cells and keratinocytes after 3 days of treatment with indicated dose of TCE5003. Error bars +/- S.E.M. **C.** FACS analysis of BrdU positive cells in SCC cells treated with vehicle or indicated dose of TCE5003 for 6 hours. Cells were pulsed with BrdU for last 2 hours of treatment. (n= 3-5), *p<0.05, **p<0.01, Error bars +/- S.E.M. **D.** *In vitro* scratch assays of indicated SCC cell lines 6 hours following scratch incubated with vehicle or indicated dose TCE3005. Quantification of area covered by cells relative to vehicle control. (n= 4-7) Error bars indicate +/- S.E.M. **E.** Growth of JHU-029 (n= 8-9 per group) Xenograft tumors treated every 2 days with vehicle (n= 8) or 10mg/kg GSK3368715 (n= 9). ***p<0.001. as assessed by Multiple measures ANOVA. Error bars +/- S.E.M.

## Discussion

Despite its’ discovery more than 30 years ago^33^, the contribution of Basonuclin 1 to keratinocyte biology and SCC pathology has remained largely unexplored. High expression of BNC1 has previously been associated with proliferative capacity in keratinocytes^26,27,46^, and our studies demonstrate that BNC1 is required for sustained proliferation of keratinocytes and SCC cultures and in SCC tumors. In the epidermis, BNC1 expression is largely restricted to the proliferative basal cells and is highest in the “basal” subtype of HNSCC and LUSC^31,32^, consistent with the idea that BNC1 contributes to maintaining cells in an undifferentiated state. The discovery of BNC1 binding to the rRNA promoter supported the hypothesis that BNC1 controlled proliferation through Pol I-dependent regulation of rRNA^35,36^, as rRNA levels are correlated with proliferation^70^. Our study now links the control of ribosomal biogenesis to direct transcriptional regulation of cell cycle genes by BNC1. Intriguingly, the transcription factor Myc, which is amplified in SCC^11^ appears to have overlapping functions, as it also can promote ribosomal biogenesis and proliferation. Surprisingly, we see little evidence that BNC1’s functions are mediated through interactions with Myc, as there was no significant enrichment of Myc binding motifs in our BNC1 binding site analysis and neither Gene Ontogeny nor pathway analysis identified the Myc pathway or target genes enrichment. A potential explanation for this is that BNC1 exerts tissue-specific control of the proliferation-differentiation-migration switch, while Myc could serve as a global amplifier of these transcriptional programs as has been described in other contexts^71,72^.

Dividing cells can permanently exit the cell cycle in response to unrepairable DNA damage, cellular stress, or activation of differentiation signals. In contrast, changes in external stimuli, such as withdrawal of growth factors can result in a reversible quiescent state which actively suppresses terminal differentiation^73^. During tissue repair, cells can also stop dividing to migrate, and in both these cases, cells retain the ability to start dividing again when environmental signals change. Despite a great deal of study, the molecular mechanisms governing the transcriptional switch from proliferation to migration in normal cells remains largely undefined^43^. We have found that while keratinocytes and SCC cells share a common core of pro-proliferative genes activated by BNC1, there is divergence in other transcriptional programs, leading to context-dependent physiological responses to BNC1 reduction. Importantly, these responses are dependent on functional interactions with IRF6 and AP-1 transcription factors, allowing the cells to choose between migration and differentiation. Previous studies have found that IRF6 promotes keratinocyte differentiation^17,59^ and our data shows that BNC1 co-localizes with IRF6 on promoters to suppress this differentiation program, and this is reinforced by the increased levels of IRF6 following BNC1 knockdown. Thus, during normal differentiation, BNC1 expression is reduced while IRF6 increases and activates keratinocyte differentiation genes. Intriguingly, while IRF6 and BNC1 have inverse protein expression in the stratified epithelial tissues, both are downregulated during wound healing. Future studies will be needed to elucidate how BNC1 and IRF6 levels are coordinately and individually controlled during wound healing and differentiation.

A key aspect of epithelial wound healing is the spatial differences between leading edge migrating cells and the more distal proliferative zone. Indeed, we observe reduced expression of BNC1 and IRF6 in the leading edge of migrating keratinocytes, while BNC1 expression is maintained away from the wound edge, consistent with its’ pro-proliferative functions. We find that knocking down both IRF6 and BNC1 in keratinocytes allows for increased migration, like what is seen in SCC cells, a context where IRF6 is frequently inactivated^63^. Intriguingly, AP-1 transcription factors have previously been shown to have a role during normal wound healing in keratinocytes and at early timepoints, the chromatin around AP-1 binding sites is opened, allowing for AP-1 regulated transcription^41,74^. In addition, JunB is expressed in leading edge keratinocytes during wound healing^75^ and *JunB*^-/-^ mice have reduced keratinocyte migration^42^. We have found that AP-1 heterodimers containing JUNB and FRA1 are found at approximately half of all BNC1 binding sites and that FRA1 is required for the pro-migratory functions of BNC1. Consistent with this data, FRA1 overexpression can accelerate migration of immortalized HaCAT cells^49^, and reduction of either JUNB or FRA1 has been shown to reduce migration and metastasis of SCC cells^47,48^. Mechanistically, we have found that loss of BNC1 increases expression of many proteins that are part of the focal adhesion complex, such as *Paxillin*, *Talin*, and *Zyxin*, and this is dependent on FRA1. Downstream, we see phosphorylation of p130 (Y165) and FAK (Y397) in response to reduction of BNC1, consistent with turnover of focal adhesions. We have also observed a FRA1-dependent switch from canonical WNT/β-catenin signaling to the WNT-Planar Cell Polarity pathway following loss of BNC1. Interestingly, FZD2, which is a key receptor for WNT5B during migration of cancer cells^76,77^ is localized in the leading edge of cells and regulates focal adhesion turnover^78^, functionally linking these pathways. In addition, FRA1 maintains expression of *TCF7L1* and *TCF7L2*, which are expressed at the leading edge of wounds, are pro-migratory^79^, and function as repressors of canonical WNT/β-catenin signaling^80^. This supports a model in which BNC1 maintains canonical WNT/β-catenin signaling in basal epithelial cells by suppressing FRA1, and this repression is relieved during migration, allowing for the activation of pro-migratory WNT-Planar Cell Polarity pathway members.

The direct targeting of transcription factors for cancer therapy has traditionally been very challenging, but success has been found in inhibition of enzymatically active co-factors^81^. However, it is important to note that these co-factors can be in complexes with a variety of different transcription factors, and thus the effects of inhibition will be highly context dependent. In SCC, we have found that loss of BNC1 reduces tumor proliferation, but the increased migration could potentially lead to increased metastasis and poor patient outcomes. It is therefore notable that the pro-proliferative program consists of BNC1-activated genes, while the migration program is largely made up of repressed target genes. To this end, we identified PRMT1 as a BNC1 co-factor, which was previously shown to promote proliferation in keratinocytes^68^ and functions in the activation of many BNC1-regulated cell cycle genes. PRMT1 inhibitors have been found to have anti-tumor activity in a variety of non-SCC malignancies^69^ and high expression of PRMT1 in HNSCC tumors correlates with poor survival^82,83^. Analysis of *in vitro* sensitivity of cells to PRMT1 inhibition showed that SCC cells were more sensitive than keratinocytes, and this translated into significant reduction in the growth of established *in vivo* SCC tumors treated with PRMT1 inhibitors. Importantly, while PRMT1 inhibition reduces proliferation, it does not increase migration, eliminating this potential downside of targeting the BNC1 transcriptional complex. In summary, we have identified a novel transcriptional network comprised of BNC1, IRF6, and JUNB/FRA1 which toggles cells from a proliferative state to a differentiated state or a migratory state, and we can specifically inhibit the pro-proliferative activities through inhibition of PRMT1.

## Acknowledgements

We would like to thank Norman Sharpless and James Rheinwald for comments on this manuscript and Daniel J. Murphy (Beatson Institute for Cancer Research) for helpful comments on this project. We would like to thank Amy Nwaobasi, Catherine Douds, and Ashley Njiru for technical support, and Matthew Wilkerson and Neil Hayes (UNC-Chapel Hill) for assistance with tumor subtype analysis. We thank Antonio Amelio (UNC-Chapel Hill) for advice and reagents and Shannan Ho Sui of the Harvard Chan Bioinformatics Core, Harvard T.H. Chan School of Public Health (Boston, MA) for assistance with the analysis of ChIP-seq data. Pathology samples were processed by the Dana-Farber/Harvard Cancer Center Specialized Histopathology Core. This work was supported by NCI R00CA157730, American Cancer Society Research Scholar Award, the Brigham and Women’s Hospital Fund to Sustain Research Excellence, and the Brigham and Women’s Hospital Department of Dermatology Fund for New Investigators (MRR). RB was supported by a SID-SunPharma fellowship. S.R.B. was supported by NIH/NCI grants R01CA247957 and R01CA258637

## Author Contributions

Conceptualization, M.R.R.; Methodology, M.R.R., S.R.B. and R.B.; Formal Analysis, M.R.R and R.B; Investigation, R.B, B.L.P., S.K., D.W., S.A.B, J.Y.K, S.X., M.P., and M.R.R.; Resources, C.D.S., C.G.L.; Data Curation, M.R.R.; Writing-Original Draft, M.R.R. and R.B.; Writing-Review & Editing, M.R.R., C.G.L., and R.B.; Visualization, M.R.R. and R.B.; Supervision, C.G.L and M.R.R; Funding Acquisition, M.R.R.

## Materials and Methods

### Cell Culture

SCC-13, SCC-15, SCC-25 ^84^, SCC-68 ^85^, G5-Ep, G5ep/TERT ^86^, OKF6/TERT-1, N/TERT-1 ^87^, and C810 (Cascade Biologics, C0015C) were cultivated in Keratinocyte SFM (ThermoFischer Scientific, 17005042) supplemented with 0.4M CaCl_2_, 0.25 ng/ml Epidermal Growth Factor (Life Technologies, 10450-013), 1/2 vial Bovine Pituitary Extract (Life Technologies, 13028014) and 100U/ml Penicillin/Streptomycin (ThermoFischer Scientific, 15190122). FaDU ^88^, JHU-029 ^89^, and NCI-H226 ^90^ were cultivated in RPMI 1640 media (ThermoFischer Scientific, 11875119) supplemented with 10% Bovine Calf Serum (ThermoFischer Scientific, SH3007203) and 100U/ml Penicillin/Streptomycin. P1F/TERT ^91^ were cultivated in 1:1 M199 (ThermoFischer Scientific, 11150059): M106 (ThermoFischer Scientific, M106500) supplemented with 15% Bovine Calf Serum, 0.4mg/ml Hydrocortisone (Millipore-Sigma, H0888)+ 10 ng/ml Epidermal Growth Factor and 100U/ml Penicillin/Streptomycin ^92^. A431, A388 ^93^, HaCAT ^94^, NIH3T3 ^95^, and 293T ^96^ were cultivated in DMEM media (ThermoFischer Scientific, 11965126) supplemented with 10% Bovine Calf Serum and 100U/ml Penicillin/Streptomycin. All cells were maintained at 37°C with 5% CO_2_. Cell line identity was authenticated using STR profiling (ATCC, 135-XV).

### RNA-seq Analysis

Cells were grown in normal culture media supplemented with 10 ng/ml doxycycline for 72 hours and total RNA was extracted using the RNA STAT-60 method (Amsbio, CS-110) according to manufacturer’s instructions. Samples were sent to Genewiz for processing and analysis. RNA was treated with DNAse, followed by Ribosomal RNA depletion, cDNA synthesis, adaptor ligation, PCR enrichment, and paired end sequencing on Illumina HiSeq. Sequence reads were trimmed to remove possible adapter sequences and nucleotides with poor quality using Trimmomatic v.0.36, then mapped to the Homo Sapiens GRCh38 reference genome available on ENSEMBL using the STAR aligner v.2.5.2b. Unique gene hit counts were calculated by using featureCounts from the Subread package v.1.5.2. Only unique reads that fell within exon regions were counted. After extraction of gene hit counts, the gene hit counts table was used for downstream differential expression analysis. Using DESeq2, a comparison of gene expression between groups of samples was performed. The Wald test was used to generate p-values and log_2_ fold changes.

### Stable Cell Line Generation

Stable cell lines were derived by lentiviral transduction as previously described ^97^. Briefly, lentiviral constructs pCMV-VSV-G and pCMV-dR8.2 dvpr ^98^ were transfected in HEK293T cells by the calcium-phosphate method (CalPhos Mammalian Transfection Kit, Takara, 631312) for virus production. Viral supernatants were collected 36 hours later and passed through a 0.45 µm filter to eliminate any contaminating HEK293T cells. Target cells were spin-infected at 900 x g for 1 hour at 25°C, and the viral supernatant was then replaced with fresh growth media. Selection was started the next day using puromycin (0.5 µg/ml), Blasticidin (2 µg/ml) or G418 (0.5 mg/ml). pCMV-VSV-G was a gift from Bob Weinberg (Addgene plasmid # 8454). pCMV-dR8.2 dvpr was a gift from Bob Weinberg (Addgene plasmid # 8455).

### Cloning

shRNA sequences were obtained from the Genetic Perturbation Portal of the Broad Institute (https://portals.broadinstitute.org/gpp/public/) and cloned into the doxycycline-inducible tetON-pLKO backbone ^99^, as previously described ^97^. Briefly, shRNA oligos were mixed, boiled at 95°C and let to cool at room temperature to allow annealing. TetON-plko constructs were digested using AgeI and EcoRI restriction enzyme and the 8.8 kb fragment was purified from a 1% agarose gel using the Qiagen Gel Extraction kit (Qiagen, 28704). Annealed shRNA oligos and digested TetON-pLKO constructs were mixed together with T4 DNA Ligase (NEB, M0202) and incubated for 3 hours at room temperature. Expression plasmids were cloned using gateway recombinational cloning ^100^. Human cells total RNA was extracted using the RNA STAT-60 method (Amsbio, CS-110) and cDNA was obtained by reverse transcription (Superscript IV Vilo Master Mix, ThermoFisher Scientific, 11756050). BNC1 and IRF6 cDNA were specifically amplified by standard PCR and inserted into a pENTR/D-TOPO entry vector (ThermoFisher Scientific, K240020). Resulting clones were transferred into pLenti CMV GFP Blast ^101^ or pcDNA3.1/nV5-DEST (ThermoFisher Scientific, 12290010) by LR recombination (ThermoFischer Scientific, 11791019) for subsequent transduction in target cells. IFN-Beta_pGL3 was a gift from Nicolas Manel (Addgene plasmid # 102597). pLenti-CMV-GFP-Blast (659-1) was a gift from Eric Campeau & Paul Kaufman (Addgene plasmid # 17445). pRL-SV40P was a gift from Ron Prywes (Addgene plasmid # 27163). Tet-pLKO-puro was a gift from Dmitri Wiederschain (Addgene plasmid # 21915).

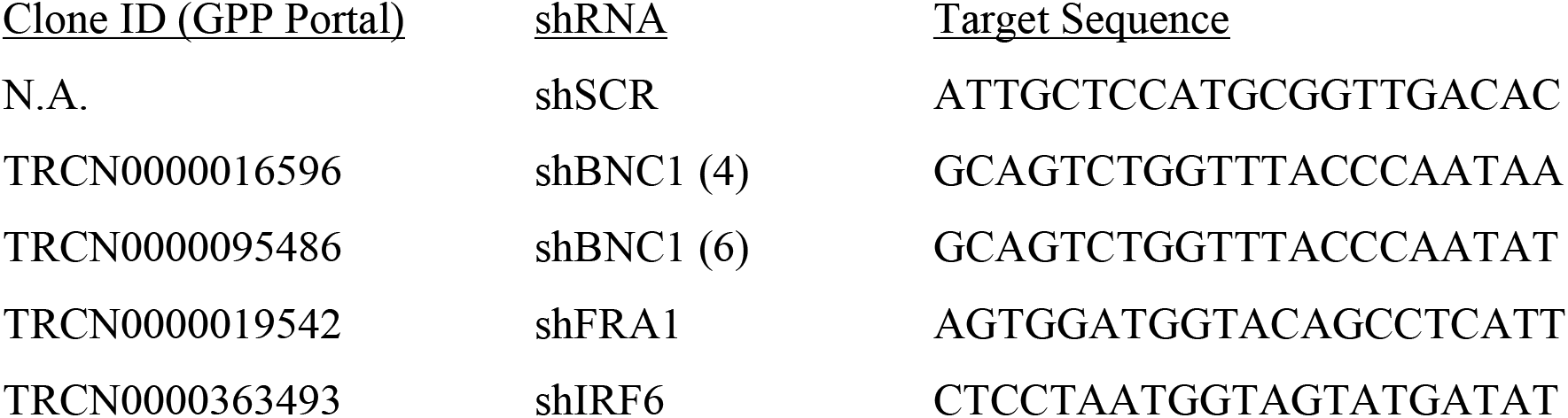

### Luciferase Assay

Luciferase assays were performed using Promega Dual-Glo Luciferase Assay system (Promega, E2920), following manufacturer protocol. Briefly, cells were transfected with pRL-SV40 (Promega, E2231) and pGL3-Control (Promega, E1741) or pGL3-IFNB (Addgene, 102597) together with expression vectors coding for BNC1 (pLenti-BNC1) and/or IRF6 (pCMV6-XL4-IRF6, Origene, SC116274), or their empty counterparts (pLenti-Empty; pCMV6-XL4-IRF6, Origene, PCMV6XL4). After 48 hours, cells were lysed in Dual Glo Reagent and firefly luminescence was measured using Promega Glomax Microplate Luminometer. DualGlo Stop & Glo Reagent was then added to the cell lysate and Renilla luminescence was measured. Luciferase activity was expressed as the ratio of Firefly over Renilla luminescence.

### ChIP-QRT-PCR

Cells were grown to 70% confluency were washed twice using ice cold PBS (ThermoFisher Scientific, 14190144) and fixed with 1% formaldehyde (Millipore-Sigma, F8775) in PBS for 15 minutes at room temperature. Fixation was stopped with the addition 0.125 M Glycine and cells were harvested and washed twice with ice cold PBS. Nuclear lysates were prepared by incubating cell pellets with Nuclear Lysis Buffer (50 mM Tris pH 8, 1% SDS, 10 mM EDTA) and sonicated using a Diagenode Bioruptor Plus (power setting: high, 10×30” with 30” cooldowns). Sheared chromatin (≈ 500-1000 bp) was incubated with the indicated antibodies in IP Dilution Buffer (1% Trito-X100, 2 mM EDTA pH 8, 150 mM NaCl, 20 mM Tris pH 8) overnight at 4°C. Antibody-chromatin complexes were captured with Dynabeads Magnetic Protein G and chromatin was decrosslinked with incubation in Elution Buffer (100 mM NaHCO3, 1% SDS) overnight at 65°C. Chromatin was purified using Zymo Research ChIP DNA Clean & Concentrator and recovered DNA was subsequently used for qPCR analysis.

Primers:

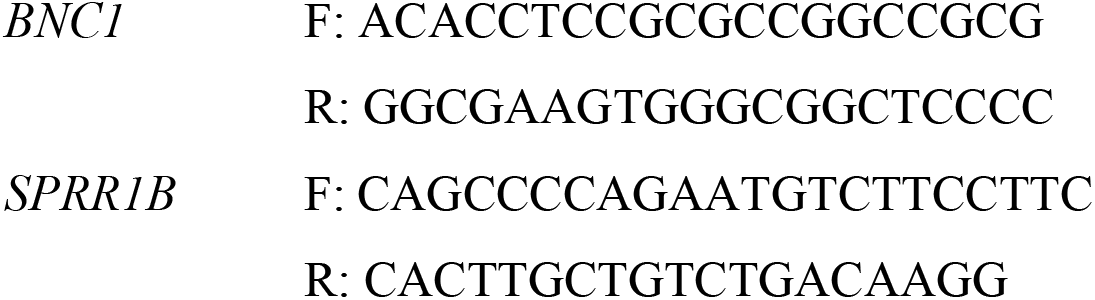

### Antibodies

*Immunoprecipitation:* BNC1 (2μg, BW175)

*Chromatin Immunoprecipitation:* BNC1 (5μl, BW175), FRA1 (25μl, Cell Signaling Technology, 3753), JUNB (25μl, Santa Cruz, sc-605), IRF6 (20μl, Novus, NBP1-51911).

*Western Blotting:* BNC1(BW175, 1:1000), IRF6 (Cell Signaling, 6948, 1:500), SPRR1B (Abcam, ab176322, 1:1000), FILAGGRIN (Abcam, Ab81468, 1:500), INVOLUCRIN (ThermoFisher Scientific, MA5-11803 1:200), p-p130 (Y165) (Cell Signaling, 4015, 1:1000), p130 (Cell Signaling, 13383, 1:1000), p-FAK (Y397) (Cell Signaling, 8556, 1:1000), ACTIN (Thermofisher Scientific, MA5-15739, 1:10000), GAPDH (Cell Signaling, 2118, 1:5000)

*Immunohistochemistry*: BNC1 (BW175, 1:800), IRF6 (Cell Signaling, 6948, 1:100), Ki67 (NCL-Ki67p, Novacastra, 1:1000)

*Immunofluorescence:* BNC1 (BW175, 1:800) Pan-cytokeratin (SCBT, sc-81714, 1:100)

### Rapid Immunoprecipitation Mass spectrometry of Endogenous proteins (RIME)

N/TERT cells were cross-linked with 1% formaldehyde for 10 minutes at room temperature, then the reaction was quenched by adding 125 mM glycine and incubating for 5 minutes. Cells were washed with PBS, lifted off the plate, then centrifuged at 4°C for 10 min at 800 x g. The cell pellet was washed once with cold PBS containing 0.5% IGEPAL-CA630 (Sigma Aldrich, I302). Pellets were then washed once with PBS-IGEPAL containing 1 mM PMSF (Sigma Aldrich, P7626) then snap frozen on dry ice. RIME analysis was performed by Active Motif, CA., according to published protocols ^65^ using IgG and BNC1 antibodies.

### ChIP-seq Analysis

SCC-13 and G5-Ep/TERT cells grown to 70% confluency in 150 mm tissue culture dishes, washed twice using ice cold PBS (ThermoFisher Scientific, 14190144) and fixed with 1% formaldehyde (Millipore-Sigma, F8775) in PBS for 15 minutes at room temperature. Fixation was stopped with the addition of 0.125 M Glycine and cells were harvested and washed twice with ice cold PBS, snap-frozen and sent to Active Motif for subsequent steps. Chromatin was isolated by adding lysis buffer, followed by disruption with a Dounce homogenizer. Lysates were sonicated and the DNA sheared to an average length of 300-500 bp with Active Motif’s EpiShear probe sonicator (cat# 53051) and cooled sonication platform (cat# 53080). Genomic DNA (Input) was prepared by treating aliquots of chromatin with RNase, proteinase K and heat for de-crosslinking, followed by ethanol precipitation. Pellets were resuspended and the resulting DNA was quantified on a NanoDrop spectrophotometer. Extrapolation to the original chromatin volume allowed quantitation of the total chromatin yield. An aliquot of chromatin (30μg) was precleared with protein A agarose beads (Invitrogen) and genomic DNA regions of interest were isolated using indicated antibodies. Complexes were washed, eluted from the beads with SDS buffer, and subjected to RNase and proteinase K treatment. Crosslinks were reversed by incubation overnight at 65°C, and ChIP DNA was purified by phenol-chloroform extraction and ethanol precipitation. Illumina sequencing libraries were prepared from the ChIP and Input DNAs using the standard consecutive enzymatic steps of end-polishing, dA-addition, and adaptor ligation using Active Motif’s custom liquid handling robotics pipeline. After the final 15 cycle PCR amplification step, the resulting DNA libraries were quantified and sequenced on Illumina NexSeq 500. Sequences (75 bp, single end) were aligned to the human genome (hg38) using the BWA algorithm (v0.7.12) _102_ with default settings. Duplicate reads were removed, and only uniquely mapped reads (mapping quality >= 25) were used for further analysis. Alignments were extended in silico at their 3’-ends to a length of 200 bp, which is the average genomic fragment length in the size-selected library and assigned to 32-nt bins along the genome. The resulting histograms (genomic “signal maps”) were stored in BigWig files. BNC1, FRA1, JUNB, and IRF6 enriched regions were identified using the MACS2 algorithm (v2.1.0) ^103^ with a cutoff of False Discovery Rate = 1.00E-07. Signal maps and peak locations were used as input data to Active Motifs proprietary analysis program, which creates Excel tables containing detailed information on sample comparison, peak metrics, peak locations, and gene annotations. Peak-associated sequences were analyzed for significantly enriched transcription factor and chromatin binding protein motifs using the **findMotifsGenome.pl** tool in the HOMER software suite ^38^.

### Immunohistochemistry and Immunofluorescence

Assistance in processing of Xenograft samples was provided by the Dana-Farber/Harvard Cancer Center Specialized Histopathology Core Facility. Formalin-fixed paraffin-embedded human or mouse tissues were cut into 5μM sections, then deparaffinized, rehydrated, and blocked with 3% hydrogen peroxide. Antigen retrieval was performed by heating slides in a pressure cooker for 45 minutes with Target Antigen Retrieval Solution (Dako, S1699). Sections were blocked with 10% goat serum for 30 minutes and primary antibodies were incubated at 4°C overnight. Sections were then incubated with the following horse radish peroxidase (HRP)-conjugated secondary antibodies for 1 hour at room temperature: goat anti-mouse IgG (H+L) (Invitrogen, 626520, 1:2000) and goat anti-rabbit IgG (H+L) (Vector Laboratories, PI-1000, 1:200). Color development was performed using a DAB chromogen solution (Dako, K3468) with an incubation time of 5 minutes. Sections were counterstained with Harris Hematoxylin (Fisher Scientific, 245-697) and then treated with Defining Solution (Fisher Scientific, 310-350) and followed by Bluing Solution (American MasterTech Scientific, HXB00242E). Quantification of Ki67 positive cells was performed by assessing number of positive cells in 1-3 independent fields for each sample.

### Western Blotting

For total cell lysates, cells were harvested and homogenized in RIPA Lysis Buffer (150 mM NaCl, 1% Igepal, 0.5% Sodium Deoxycholate, 0.1% SDS, 25mM Tris pH 7.4) for 30 minutes at 4°C followed by centrifugation at ≥ 15,000 x g for 10 minutes at 4°C. Protein concentration of clarified cell lysates was assessed with Bradford reagent (BioRad, 5000001). 20 µg of proteins were mixed with Laemmli buffer (BioRad, 1610737) containing 5% 2-Mercaptoethanol (Sigma-Aldrich, M6250) and boiled at 95°C for 5 minutes. Denatured lysates were run on 4-20% gradient Tris-Glycine polyacrylamide gels (Biorad, 4561096) at 200V for 30 minutes, then transferred at 100V for 90 minutes to a low fluorescence PVDF membrane (Millipore-Sigma, IPFL00010) using a wet tank transfer system. After drying and rehydration, membranes were blocked with the Odyssey TBS blocking buffer (LI-COR, 927-50000) and incubated overnight on a shaker with the primary antibody diluted in the blocking buffer supplemented with 0.2% Tween 20. After 3 washes with TBS-0.1% Tween 20, membranes were incubated with the secondary anti-mouse (LI-COR, 926-680721:10,000) or anti-rabbit (LI-COR, 926-32211, 1:15000) antibodies in blocking buffer supplemented with 0.2% Tween20 for 1 hour at room temperature. Membranes were washed again three times and image were acquired using the Odyssey CLX imaging system (ThermoFisher).

### Colony Formation Assay

Cells were plated at a density of 500-1000 cells/cm^2^ in 6 well plates and allowed to grow for 2 weeks in normal culture media or growth media containing 10 ng/ml doxycycline (Sigma-Aldrich, D9891). Cultures were then fixed with 10% buffered formalin (Fisher Scientific, 245-684) for 20 minutes at room temperature, thoroughly washed with water, stained with 0.1% crystal violet (Sigma-Aldrich, C0775) for 20 minutes at room temperature and washed again with water. To get a quantification of cell growth, crystal violet staining was extracted using 10% acetic acid and optical density was measured with a spectrophotometer at 595 nm.

### Cell Growth Inhibition Assay

Drug inhibition assays were performed using TCE-5003 (Tocris, 5099) dissolved in DMSO. Dose-response curves were performed in 96-well plates, by seeding 2,000-10,000 cells in triplicate wells, the day before treatment. The following day, compounds were added at corresponding serial dilutions, from the highest dose. Three days after treatment, cell viability was measured using CellTiter 96 AQueous One Solution Cell Proliferation Assay (Promega, G3582).

### *In Vitro* Migration Assay

Cells were plated at full confluency in silicone 2-well culture inserts (Ibidi, 80209) and incubated in growth media containing 10 ng/ml doxycycline (Sigma-Aldrich, D9891) for 48h to induce shRNA expression. Culture inserts were then gently removed, and pictures of the cell-free gap were taken every 2 hours to monitor cell migration. For PRMT1 inhibitor treatment, cells were seeded and incubated for 18 hours, culture inserts were removed, and then cells were treated with vehicle or 1μM TCE5003 (Tocris, 5099). Quantification of cell migration was made using Image J software (https://imagej.nih.gov).

### Fluorescence-Activated Cell Sorting

Cells grown in normal culture media supplemented with 10 ng/ml doxycycline for 48 hours to induce shRNA or treated with 1 μM TCE5003 (Tocris, 5099) for 22 hours. Cells were then incubated for 2 hours with 3 µg/ml 5-bromo-2’-deoxyurine (Sigma-Aldrich, B5002). Cells were then detached using trypsin/EDTA (GIBCO, 25200056) and fixed with ice cold 90% ethanol in PBS. Nuclei were extracted by 20 minutes treatment of the cell pellet with 0.08% Pepsin solution and chromatin was denatured by 20 minutes incubation with 2M HCl. Recovered nuclei were stained with FITC-conujugated α-5-bromo-2’-deoxyurine (Invitrogen, MoBu-1) in staining buffer (2 mM HEPES pH 7.4, 30 mM NaCl, 0.8% FBS, 0.02% Sodium azide), and washed with IFA buffer (10 mM HEPES pH 7.4, 150 mM NaCl, 4% FBS, 0.1% Sodium Azide, 0.5% Igepal). Stained nuclei were analyzed using a BD FACSCanto flow cytometer and quantification was performed with FlowJo software (BD Life Science).

### Animals

Xenograft tumors were generated by subcutaneous injection into *Nude* mice of 1.00E+06 NIH3T3 + 1.00E+06 SCC-13 or 1.00E+06 NIH3T3 + 1.00E+06 JHU-029 in 50/50 (v/v) PBS:Matrigel (Corning, 354234). For shRNA experiments, 7 days after cell injections water was exchanged with 2% sucrose and 2 mg/mL doxycycline water which was freshly prepared every other day. For PRMT1 inhibitor experiments, 10mg/kg GSK3368715 (MedChemExpress USA, Monmouth Junction, NJ) was injected IP every two days starting 7 days after cell injections. Tumor volumes were calculated using the formula: tumor volume (mm^3^) = 4/3π x (length/2) x (width/2). For wound studies, mice were anesthetized using isoflurane, followed by shaving the back with electric clippers. Sterile silicone rings were glued and sutured onto the back of mice. Full-thickness excisional wounds (5 mm in size) were made inside the silicone ring of anesthetized mice using sterile biopsy punches. The day of the punch biopsy is considered as day 0.

### Statistical Analysis

For cell cycle analysis, colony forming assays, migration assays, ChIP, QRT-PCR, and Luciferase assays, Students unpaired t-test was used to determine statistical significance. For xenograft growth experiments, P-values were determined using multiple-measures analysis of variance. Ki67 IHC staining in Xenograft tumors was analyzed using the Mann-Whitney test. SCC-specific transcription factors were identified using data from the International Genomics Consortium Expression Project for Oncology ^25^ (GEO: GSE2109). We compared gene expression profiles of SCC tumors to all other tumor types using the NCBI GEO2R tool, and genes overexpressed in SCC tumors compared to non-SCC tumors were rank-ordered by Benjamini & Hochberg adjusted p-value. P-values for Gene Ontogeny, Reactome and ARCHS4 as analyzed using Enrichr ^45^ were determined using Fisher exact test. Only data sets with a false discovery rate of less than 5% were included. P-values less than 0.05 were considered significant for all experiments.

### Ethics Approvals

All animals were housed and treated in accordance with protocols approved by the Brigham and Women’s Hospital Institutional Animal Care and Use Committee (IACUC). The Mass General Brigham IRB approved use of tissues for this study. All human tissue studies used exclusively de-identified and discarded material collected in the course of routine clinical care, for which the Mass General Brigham IRB has determined that signed informed consent was not required.

### Availability of Data and Materials

The datasets generated and/or analyzed during the current study are available in the NCBI Gene Expression Omnibus (GSE212906, GSE209993, GSE212905).

