## Supplementary Figures S1-S6 for "PRMT1 Inhibition Selectively Targets BNC1-Dependent Proliferation, but not Migration in Squamous Cell Carcinoma"

### Supplemental information

**Table S1:** GSE2109 Histology key, Related to Figure 1

**Table S2:** ExpO differentially expressed genes, Related to Figure 1

**Table S3:** Significant differentially expressed genes in keratinocytes following BNC1 knockdown, Related to Figure 2

**Table S4:** Significant differentially expressed genes in SCC following BNC1 knockdown, Related to Figure 2

**Table S5:** Genomic Intervals bound by BNC1 in G5Ep-TERT, Related to Figure 2

**Table S6:** Genomic Intervals bound by BNC1 in SCC-13, Related to Figure 2

**Table S7:** HOMER motif analysis of BNC1 bound genomic regions, Related to Figure 2

**Table S8:** Gene Enrichment analysis of BNC1 bound & regulated genes, Related to Figure 2

**Table S9:** Genomic regions bound by FRA1, and JUNB in SCC-13 cells, related to Figure 4

**Table S10:** HOMER motif analysis of FRA1 and JUNB bound genes, Related to Figure S4

**Table S11:** Differentially expressed genes in SCC-13 following BNC1, FRA1, BNC1-FRA1 knockdown, Related to Figure 4

**Table S12:** ARCHS4 analysis of BNC1 repressed genes in keratinocytes, Related to Figure 6

**Table S13:** Genomic Intervals bound by IRF6 in G5Ep-TERT, Related to Figure 6

**Table S14:** BNC1 bound and regulated genes in keratinocytes regulated by PRMT1, Related to Figure 7

**Figure S1:** Validation of BW175 $\alpha$-BNC1 antibody, Related to Figure 1

**Figure S2:** Analysis of BNC1 DNA binding, Related to Figure 2

**Figure S3:** BNC1-dependent regulation of proliferation, Related to Figure 3

**Figure S4:** BNC1-dependent regulation of migration, Related to Figure 4

**Figure S5:** BNC1 in keratinocyte migration and differentiation, Related to Figure 5

**Figure S6:** IRF6-BNC1 interactions in keratinocytes Related to Figure 6

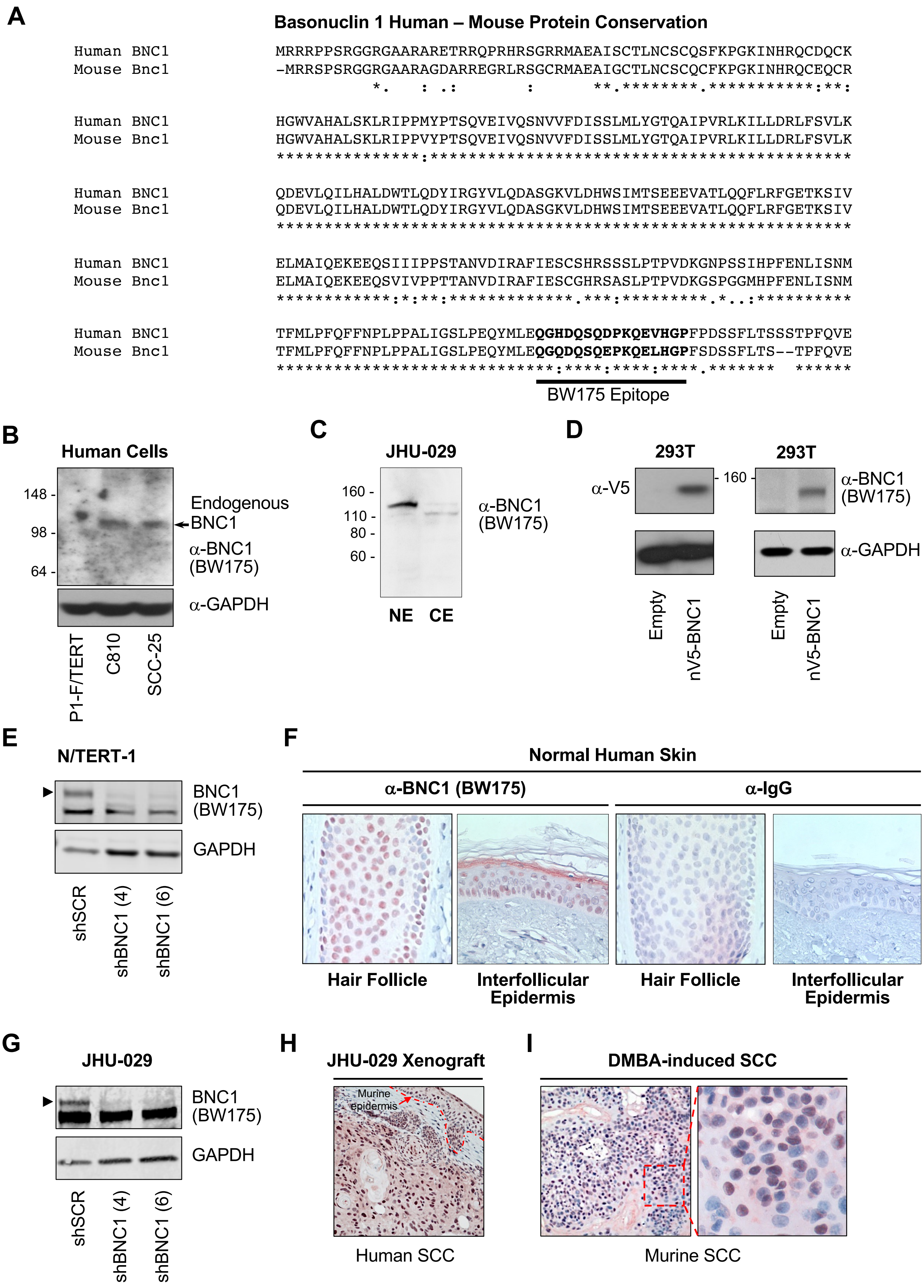


**Supplemental Figure 1**

**Figure S1: Validation of BW175 α-BNC1 antibody, Related to Figure 1**

**A.** Sequence of the N-terminus of Human and Murine Basonuclin 1 with the location of the human epitope for BW175 indicated, along with the corresponding murine sequence.

**B.** Western blot analysis of BNC1 (indicated by arrow) in human P1-F/TERT (fibroblast), C810 (keratinocyte), and SCC-25 (Squamous Cell Carcinoma) cells using BW175 α-BNC1 antibody. GAPDH serves as a loading control.

**C.** Western blot analysis of localization of BNC1 in nuclear (NE) and cytoplasmic (CE) extracts from JHU-029 HNSCC cells using the BW175 antibody.

**D.** Western blot analysis of over-expressed BNC1-V5 in 293T cells using V5 epitope tag antibody (left) and α-BNC1 antibody BW175 (right). GAPDH serves as a loading control.

**E.** Western blot analysis of BNC1 expression in the immortalized keratinocyte line OKF6/TERT after 48 hours of treatment with Doxycycline (10ng/mL) to induce shRNA-mediated knock-down of BNC1. ACTIN serves as a loading control.

**F.** Immunohistochemical analysis of human skin stained with α-BNC1 antibody BW175 (left) or IgG isotype control (right).

**G.** Western blot analysis of BNC1 expression in SCC-13 cells 48 hours lentiviral infection of indicated shRNA. GAPDH serves as a loading control.

**H.** Immunohistochemical analysis of BNC1 using BW175 antibody in human HNSCC Xenografts, and endogenous murine SCC tumors.

**I.** Immunohistochemical analysis of BNC1 using BW175 antibody in DMBA-induced murine SCC tumors.

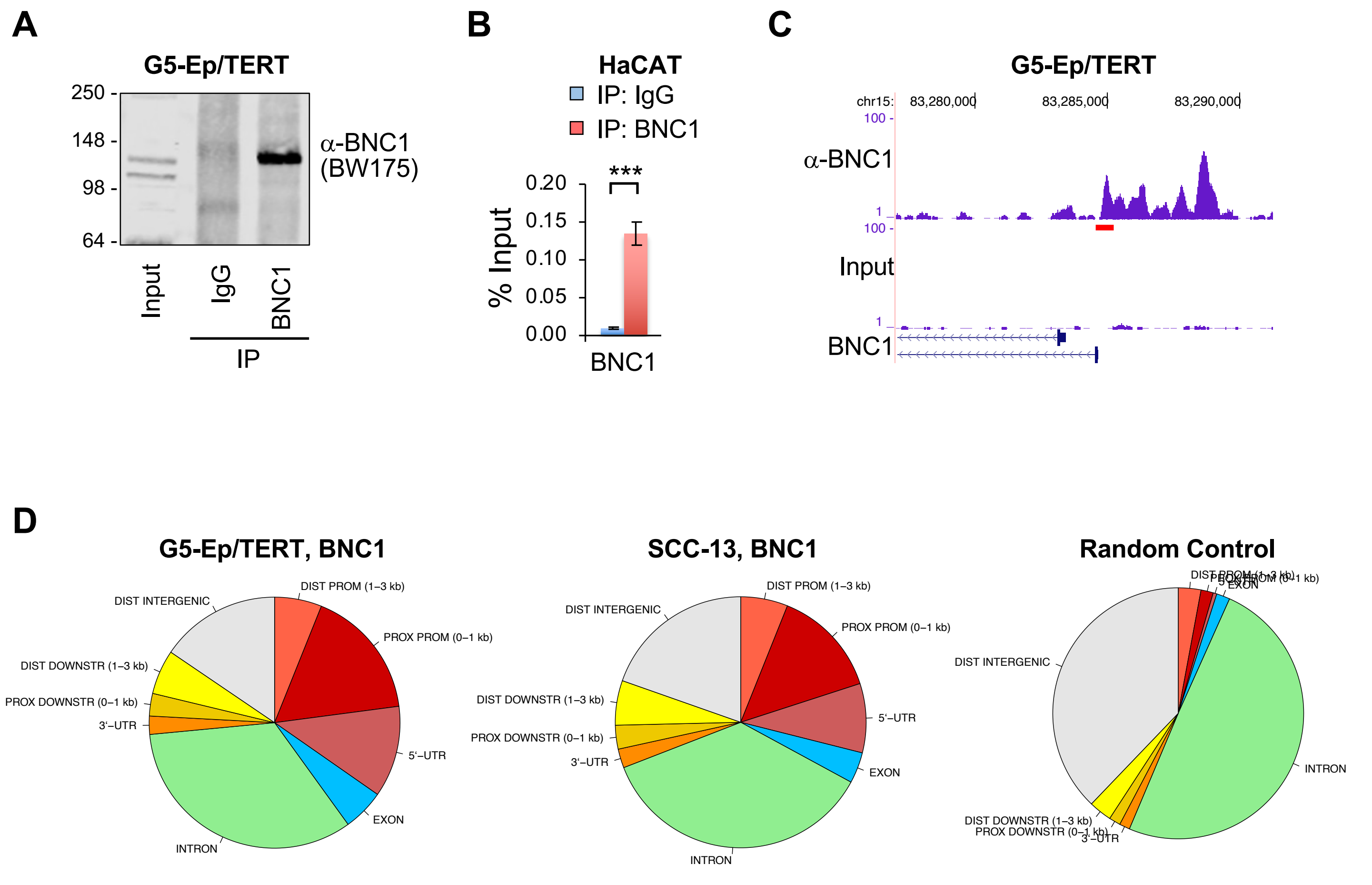


**Supplemental Figure 2**

**Figure S2: Analysis of BNC1 DNA binding, Related to Figure 2**

**A.** Immunoprecipitation of BNC1 protein in keratinocytes using BW175 antibody or IgG control antibody.

**B.** Quantitative Real-Time PCR of previously reported binding sites near the *BNC1* gene in HaCAT cells following ChIP using α-BNC1 or α-IgG antibodies. ***$p<0.001$

**C.** ChIP-seq binding peaks of BNC1 in G5-Ep/TERT cells at the *BNC1* gene. Red bar indicates location of primers used in (B).

**D.** Distribution of sequenced tags following ChIP-seq analysis of BNC1 located in different genomic features in indicated cell lines. Random control represents randomly located "peaks" run against the same genomic features database

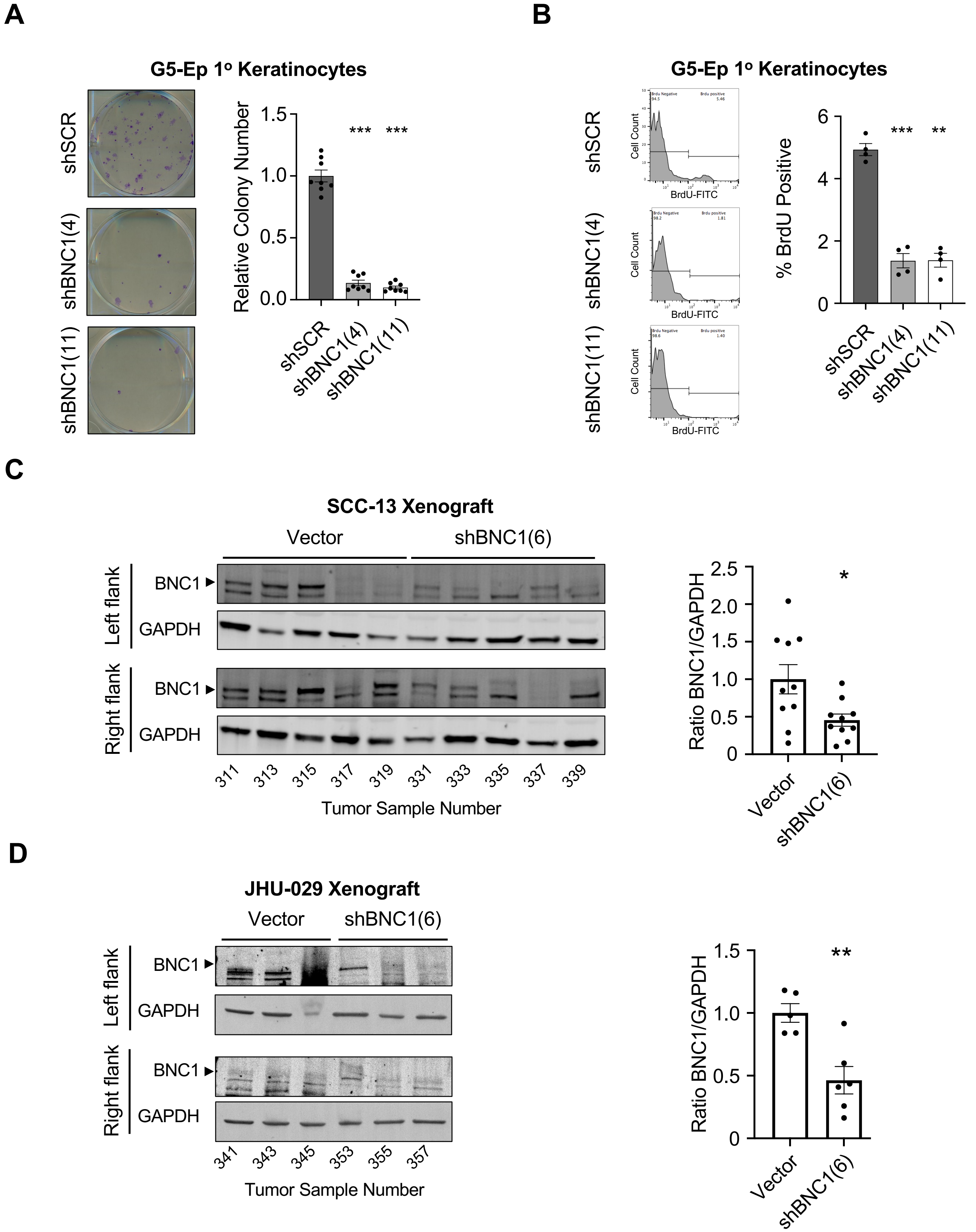


**Supplemental Figure 3**

**Figure S3: BNC1-dependent regulation of proliferation, Related to Figure 3**

**A.** Colony forming assays in primary keratinocytes 2 weeks after infection with indicated shRNA construct. Quantification (right) of colony number by OD595 of extracted crystal violet stain (n=8 wells). ***$p<0.001$. Error bars +/- Std. Dev.

**B.** FACS analysis of BrdU positive cells in primary keratinocytes with indicated shRNA construct after a 2-hour pulse of BrdU. Quantification (right) of BrdU positive cells (n=4). **$p<0.01$, ***$p<0.001$. Error bars +/- Std. Dev.

**C.** Western blot analysis of BNC1 levels in SCC-13 Xenograft tumors following treatment with doxycycline to induce BNC1 shRNA. GAPDH serves as a loading control.

**D.** Western blot analysis of BNC1 levels in JHU-029 Xenograft tumors following treatment with doxycycline to induce BNC1 shRNA. GAPDH serves as a loading control.

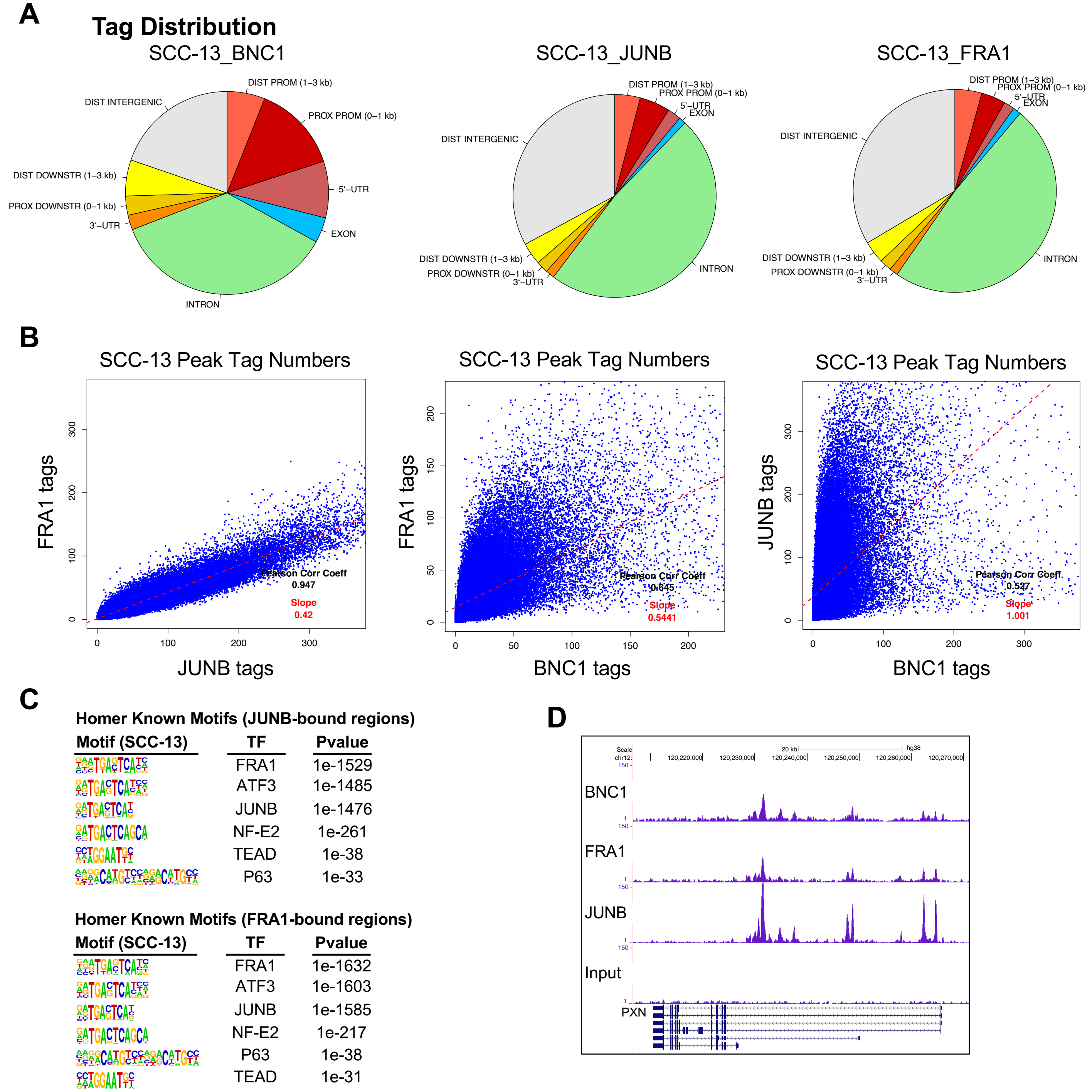


**Supplemental Figure 4**

**Figure S4: BNC1-dependent regulation of migration, Related to Figure 4**

**A.** Distribution of sequenced tags following ChIP-seq analysis of BNC1 and FRA1 located in different genomic features in SCC-13 cells.

**B.** Correlation between sequenced tags following ChIP-seq analysis of BNC1, FRA1, and JUNB in SCC-13 cells.

**C.** HOMER known motif enrichment analysis of JUNB (top) and FRA-1 (bottom bound regions in SCC cells.

**D.** DNA binding peaks of BNC1, FRA1, and JUNB in the *Paxillin* (*PXN*) gene in SCC-13 cells, assessed by ChIP-seq.

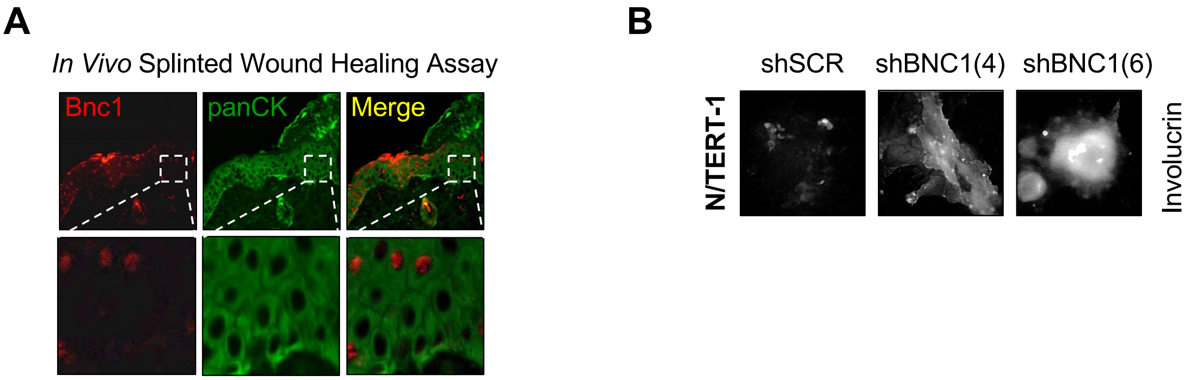


**Supplemental Figure 5**

**Figure S5: BNC1 in keratinocyte migration and differentiation, Related to Figure 5**

**A.** Bnc1 and pan-cytokeratin (Pan-CK) immunofluorescence of splinted wound 7 days after wounding.

**B.** Immunofluorescent staining of Involucrin in N/TERT-1 keratinocytes 96 hours after treatment with doxycycline to induce indicated shRNA construct.

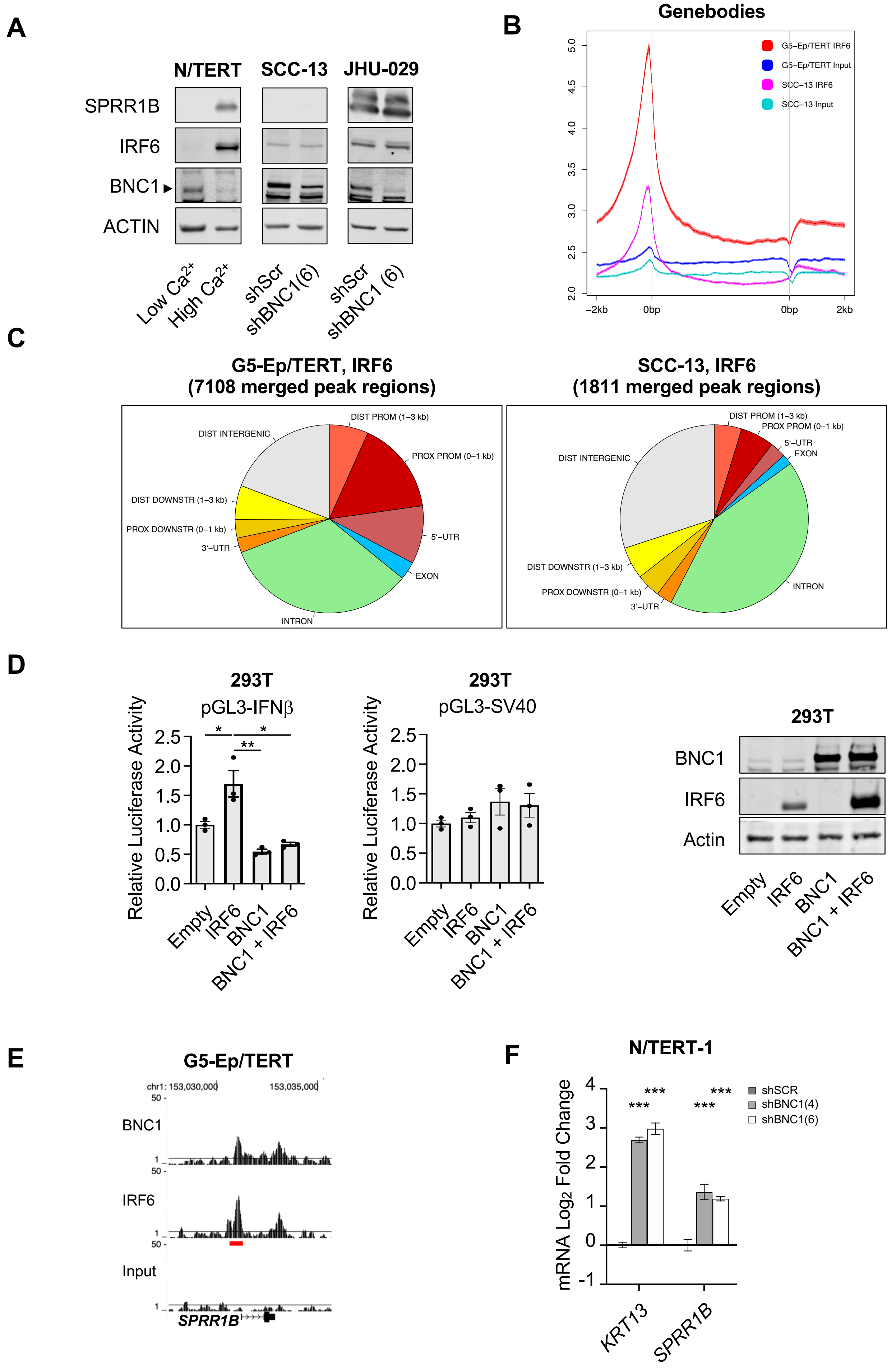


**Supplementary Figure 6**

**Figure S6: IRF6-BNC1 interactions in keratinocytes Related to Figure 6**

**A.** Western blot analysis of differentiation markers in keratinocytes and SCC cells 120 hours after induction of indicated shRNAs. N/TERT-1 cells grown in low $Ca^{2+}$ (0.04mM) and high $Ca^{2+}$ (2.0mM) serves as a positive control for induced differentiation. Actin serves as a loading control.

**B.** Average distribution of IRF6 near gene bodies in keratinocytes (G5-Ep/TERT) and SCC cells (SCC-13) assessed by ChIP-seq.

**C.** Distribution of sequenced tags following ChIP-seq analysis of IRF6 located in different genomic features in G5-Ep/TERT and SCC-13 cells.

**D.** Luciferase reporter assay of 293T cells transfected with the indicated constructs. *$p<0.05$, **$p<0.01$, Error bars indicate +/- S.E.M. Western blot analysis of indicated proteins in 293T cells used for luciferase assay. Actin serves as a loading control.

**E.** ChIP-seq binding peaks for BNC1 and IRF6 around the *SPRR1B* locus in keratinocytes. Red bar indicates primer locations.

**F.** QRT-PCR analysis of indicated mRNA transcripts 72 hours after induction of indicated shRNA construct (n=3) in keratinocytes. ***$p<0.001$. Error bars indicate +/- S.E.M.
