## Supplemental Table S7 for "PRMT1 Inhibition Selectively Targets BNC1-Dependent Proliferation, but not Migration in Squamous Cell Carcinoma"

### Homer Known Motif Enrichment Results (QC/homer/1_03HI_G5epTERT_BNC1_hg38)

Homer *de novo* Motif Results
Gene Ontology Enrichment Results
Known Motif Enrichment Results (txt file)
Total Target Sequences = 972, Total Background Sequences = 45293

| Rank | Motif | Name | P-value | log P-pvalue | q-value (Benjamini) | # Target Sequences with Motif | % of Targets Sequences with Motif | # Background Sequences with Motif | % of Background Sequences with Motif |
|---|---|---|---|---|---|---|---|---|---|
| 1 | 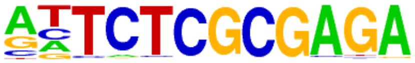 | GFX(?)/Promoter/Homer | 1e-363 | -8.367e+02 | 0.0000 | 273.0 | 28.09% | 262.7 | 0.58% |
| 2 | 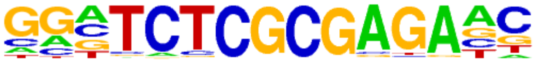 | ZBTB33(Zf)/GM12878-ZBTB33-ChIP-Seq(GSE32465)/Homer | 1e-357 | -8.240e+02 | 0.0000 | 345.0 | 35.49% | 690.6 | 1.53% |
| 3 | 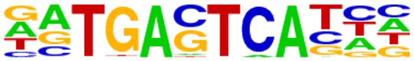 | Atf3(bZIP)/GBM-ATF3-ChIP-Seq(GSE33912)/Homer | 1e-103 | -2.372e+02 | 0.0000 | 194.0 | 19.96% | 1237.9 | 2.74% |
| 4 | 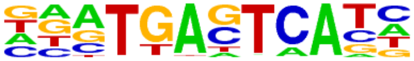 | Fra1(bZIP)/BT549-Fra1-ChIP-Seq(GSE46166)/Homer | 1e-96 | -2.222e+02 | 0.0000 | 176.0 | 18.11% | 1065.2 | 2.36% |
| 5 | 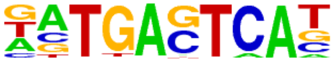 | BATF(bZIP)/Th17-BATF-ChIP-Seq(GSE39756)/Homer | 1e-93 | -2.153e+02 | 0.0000 | 183.0 | 18.83% | 1224.5 | 2.71% |
| 6 | 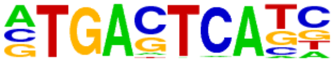 | AP-1(bZIP)/ThioMac-PU.1-ChIP-Seq(GSE21512)/Homer | 1e-92 | -2.127e+02 | 0.0000 | 197.0 | 20.27% | 1481.7 | 3.28% |
| 7 | 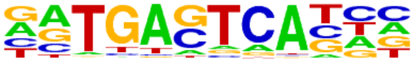 | Fosl2(bZIP)/3T3L1-Fosl2-ChIP-Seq(GSE56872)/Homer | 1e-91 | -2.103e+02 | 0.0000 | 148.0 | 15.23% | 743.2 | 1.64% |
| 8 | 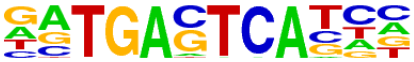 | Jun-AP1(bZIP)/K562-cJun-ChIP-Seq(GSE31477)/Homer | 1e-74 | -1.715e+02 | 0.0000 | 116.0 | 11.93% | 542.0 | 1.20% |
| 9 | 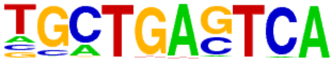 | Bach2(bZIP)/OCILy7-Bach2-ChIP-Seq(GSE44420)/Homer | 1e-46 | -1.080e+02 | 0.0000 | 85.0 | 8.74% | 496.6 | 1.10% |
| 10 | 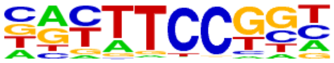 | Fli1(ETS)/CD8-FLI-ChIP-Seq(GSE20898)/Homer | 1e-27 | -6.271e+01 | 0.0000 | 287.0 | 29.53% | 7049.8 | 15.59% |
| 11 | 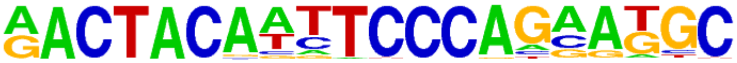 | GFY-Staf(?,Zf)/Promoter/Homer | 1e-26 | -6.191e+01 | 0.0000 | 66.0 | 6.79% | 568.2 | 1.26% |
| 12 | 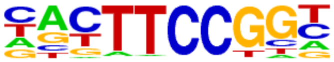 | Elk1(ETS)/Hela-Elk1-ChIP-Seq(GSE31477)/Homer | 1e-26 | -6.060e+01 | 0.0000 | 223.0 | 22.94% | 4913.8 | 10.87% |
| 13 | 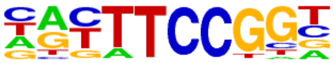 | Elk4(ETS)/Hela-Elk4-ChIP-Seq(GSE31477)/Homer | 1e-25 | -5.811e+01 | 0.0000 | 223.0 | 22.94% | 5007.9 | 11.07% |
| 14 | | GFY(?)/Promoter/Homer | 1e-23 | -5.484e+01 | 0.0000 | 62.0 | 6.38% | 570.6 | 1.26% |

| | | | | | | | | | |
|---|---|---|---|---|---|---|---|---|---|
| | 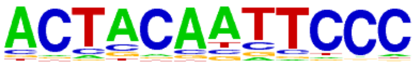 | | | | | | | | |
| 15 | 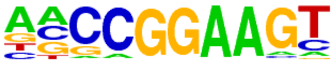 | ELF1(ETS)/Jurkat-ELF1-ChIP-Seq(SRA014231)/Homer | 1e-23 | -5.329e+01 | 0.0000 | 198.0 | 20.37% | 4356.5 | 9.63% |
| 16 | 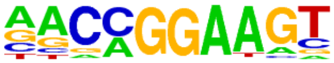 | ETV1(ETS)/GIST48-ETV1-ChIP-Seq(GSE22441)/Homer | 1e-21 | -5.025e+01 | 0.0000 | 279.0 | 28.70% | 7340.3 | 16.23% |
| 17 | 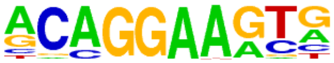 | ERG(ETS)/VCaP-ERG-ChIP-Seq(GSE14097)/Homer | 1e-20 | -4.747e+01 | 0.0000 | 275.0 | 28.29% | 7333.0 | 16.22% |
| 18 | 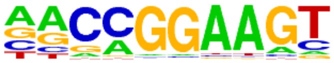 | GABPA(ETS)/Jurkat-GABPa-ChIP-Seq(GSE17954)/Homer | 1e-20 | -4.727e+01 | 0.0000 | 216.0 | 22.22% | 5212.3 | 11.53% |
| 19 | 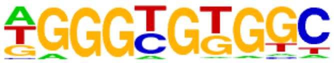 | KLF5(Zf)/LoVo-KLF5-ChIP-Seq(GSE49402)/Homer | 1e-20 | -4.721e+01 | 0.0000 | 374.0 | 38.48% | 11209.4 | 24.79% |
| 20 | 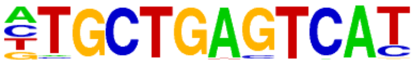 | Nrf2(bZIP)/Lymphoblast-Nrf2-ChIP-Seq(GSE37589)/Homer | 1e-19 | -4.482e+01 | 0.0000 | 27.0 | 2.78% | 106.6 | 0.24% |
| 21 | 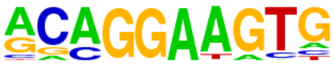 | ETS1(ETS)/Jurkat-ETS1-ChIP-Seq(GSE17954)/Homer | 1e-19 | -4.391e+01 | 0.0000 | 222.0 | 22.84% | 5577.4 | 12.33% |
| 22 | 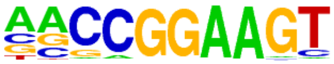 | ETS(ETS)/Promoter/Homer | 1e-17 | -4.000e+01 | 0.0000 | 130.0 | 13.37% | 2650.3 | 5.86% |
| 23 | 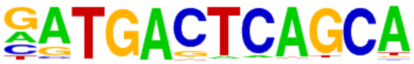 | NF-E2(bZIP)/K562-NFE2-ChIP-Seq(GSE31477)/Homer | 1e-17 | -3.980e+01 | 0.0000 | 27.0 | 2.78% | 130.2 | 0.29% |
| 24 | 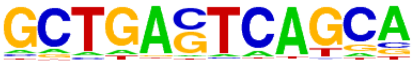 | MafK(bZIP)/C2C12-MafK-ChIP-Seq(GSE36030)/Homer | 1e-16 | -3.845e+01 | 0.0000 | 54.0 | 5.56% | 619.2 | 1.37% |
| 25 | 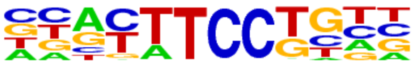 | Etv2(ETS)/ES-ER71-ChIP-Seq(GSE59402)/Homer(0.967) | 1e-15 | -3.670e+01 | 0.0000 | 174.0 | 17.90% | 4230.0 | 9.35% |
| 26 | 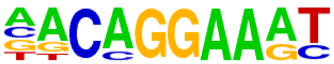 | EWS:FLI1-fusion(ETS)/SK_N_MC-EWS:FLI1-ChIP-Seq(SRA014231)/Homer | 1e-15 | -3.498e+01 | 0.0000 | 129.0 | 13.27% | 2798.4 | 6.19% |
| 27 | 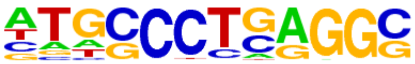 | AP-2alpha(AP2)/Hela-AP2alpha-ChIP-Seq(GSE31477)/Homer | 1e-13 | -3.206e+01 | 0.0000 | 241.0 | 24.79% | 6926.8 | 15.32% |
| 28 | 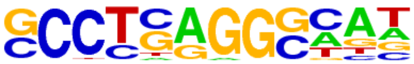 | AP-2gamma(AP2)/MCF7-TFAP2C-ChIP-Seq(GSE21234)/Homer | 1e-13 | -3.142e+01 | 0.0000 | 292.0 | 30.04% | 8964.5 | 19.82% |
| 29 | 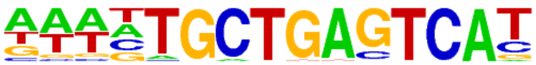 | Bach1(bZIP)/K562-Bach1-ChIP-Seq(GSE31477)/Homer | 1e-12 | -2.859e+01 | 0.0000 | 19.0 | 1.95% | 91.5 | 0.20% |
| 30 | 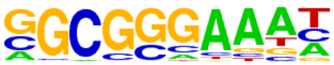 | E2F4(E2F)/K562-E2F4-ChIP-Seq(GSE31477)/Homer | 1e-12 | -2.843e+01 | 0.0000 | 200.0 | 20.58% | 5601.7 | 12.39% |
| 31 | | YY1(Zf)/Promoter/Homer | 1e-11 | -2.599e+01 | 0.0000 | 45.0 | 4.63% | 622.8 | 1.38% |

| | | | | | | | | | |
|---|---|---|---|---|---|---|---|---|---|
| | 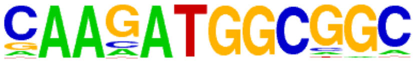 | | | | | | | | |
| 32 | 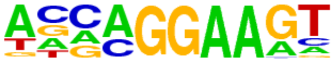 | EHF(ETS)/LoVo-EHF-ChIP-Seq(GSE49402)/Homer | 1e-11 | -2.589e+01 | 0.0000 | 185.0 | 19.03% | 5198.0 | 11.49% |
| 33 | 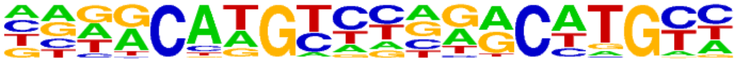 | p63(p53)/Keratinocyte-p63-ChIP-Seq(GSE17611)/Homer | 1e-10 | -2.476e+01 | 0.0000 | 63.0 | 6.48% | 1127.8 | 2.49% |
| 34 | 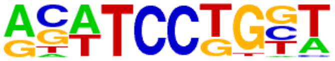 | SPDEF(ETS)/VCaP-SPDEF-ChIP-Seq(SRA014231)/Homer | 1e-10 | -2.356e+01 | 0.0000 | 153.0 | 15.74% | 4158.3 | 9.20% |
| 35 | 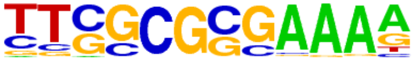 | E2F(E2F)/Hela-CellCycle-Expression/Homer | 1e-9 | -2.303e+01 | 0.0000 | 41.0 | 4.22% | 582.7 | 1.29% |
| 36 | 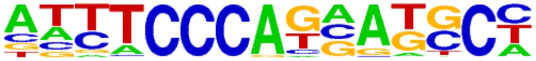 | ZNF143\|STAF(Zf)/CUTLL-ZNF143-ChIP-Seq(GSE29600)/Homer | 1e-9 | -2.171e+01 | 0.0000 | 76.0 | 7.82% | 1614.9 | 3.57% |
| 37 | 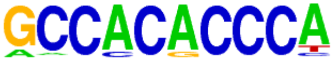 | Klf4(Zf)/mES-Klf4-ChIP-Seq(GSE11431)/Homer | 1e-9 | -2.078e+01 | 0.0000 | 103.0 | 10.60% | 2540.7 | 5.62% |
| 38 | 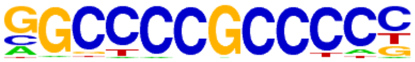 | Sp1(Zf)/Promoter/Homer | 1e-9 | -2.074e+01 | 0.0000 | 146.0 | 15.02% | 4072.2 | 9.00% |
| 39 | 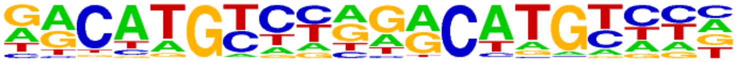 | p53(p53)/Saos-p53-ChIP-Seq(GSE15780)/Homer | 1e-8 | -2.038e+01 | 0.0000 | 23.0 | 2.37% | 222.8 | 0.49% |
| 40 | 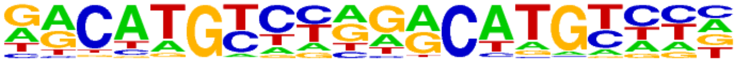 | p53(p53)/Saos-p53-ChIP-Seq/Homer | 1e-8 | -2.038e+01 | 0.0000 | 23.0 | 2.37% | 222.8 | 0.49% |
| 41 | 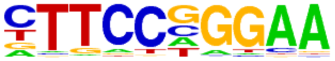 | Stat3(Stat)/mES-Stat3-ChIP-Seq(GSE11431)/Homer | 1e-8 | -1.988e+01 | 0.0000 | 87.0 | 8.95% | 2043.2 | 4.52% |
| 42 | 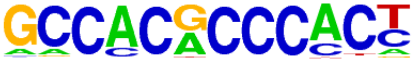 | Klf9(Zf)/GBM-Klf9-ChIP-Seq(GSE62211)/Homer | 1e-8 | -1.948e+01 | 0.0000 | 123.0 | 12.65% | 3311.8 | 7.32% |
| 43 | 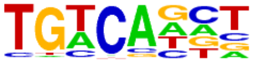 | Tgif2(Homeobox)/mES-Tgif2-ChIP-Seq(GSE55404)/Homer | 1e-8 | -1.931e+01 | 0.0000 | 279.0 | 28.70% | 9427.3 | 20.85% |
| 44 | 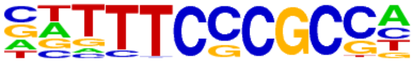 | E2F7(E2F)/Hela-E2F7-ChIP-Seq(GSE32673)/Homer | 1e-8 | -1.869e+01 | 0.0000 | 65.0 | 6.69% | 1382.6 | 3.06% |
| 45 | 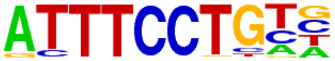 | EWS:ERG-fusion(ETS)/CADO_ES1-EWS:ERG-ChIP-Seq(SRA014231)/Homer | 1e-7 | -1.806e+01 | 0.0000 | 83.0 | 8.54% | 1992.4 | 4.41% |
| 46 | 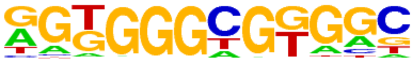 | KLF14(Zf)/HEK293-KLF14.GFP-ChIP-Seq(GSE58341)/Homer | 1e-7 | -1.706e+01 | 0.0000 | 406.0 | 41.77% | 15127.1 | 33.45% |
| 47 | 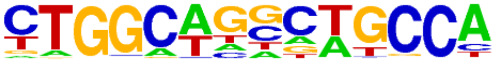 | Tlx?(NR)/NPC-H3K4me1-ChIP-Seq(GSE16256)/Homer | 1e-7 | -1.633e+01 | 0.0000 | 67.0 | 6.89% | 1540.9 | 3.41% |
| 48 | | Tbx5(T-box)/HL1-Tbx5.biotin-ChIP-Seq(GSE21529)/Homer | 1e-7 | -1.624e+01 | 0.0000 | 339.0 | 34.88% | 12294.6 | 27.19% |

| | | | | | | | | |
|---|---|---|---|---|---|---|---|---|
| | 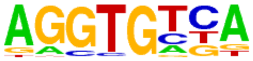 | | | | | | | |
| 49 | 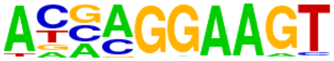 | ELF5(ETS)/T47D-ELF5-ChIP-Seq(GSE30407)/Homer | 1e-6 | -1.593e+01 | 0.0000 | 105.0 | 10.80% | 2873.2 | 6.35% |
| 50 | 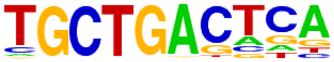 | MafA(bZIP)/Islet-MafA-ChIP-Seq(GSE30298)/Homer | 1e-6 | -1.523e+01 | 0.0000 | 100.0 | 10.29% | 2735.4 | 6.05% |
| 51 | 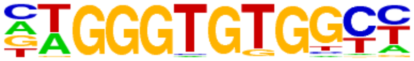 | EKLF(Zf)/Erythrocyte-Klf1-ChIP-Seq(GSE20478)/Homer | 1e-6 | -1.497e+01 | 0.0000 | 45.0 | 4.63% | 900.1 | 1.99% |
| 52 | 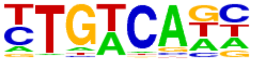 | Tgif1(Homeobox)/mES-Tgif1-ChIP-Seq(GSE55404)/Homer | 1e-6 | -1.496e+01 | 0.0000 | 237.0 | 24.38% | 8120.6 | 17.96% |
| 53 | 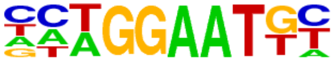 | TEAD(TEA)/Fibroblast-PU.1-ChIP-Seq(Unpublished)/Homer | 1e-6 | -1.420e+01 | 0.0000 | 56.0 | 5.76% | 1273.4 | 2.82% |
| 54 | 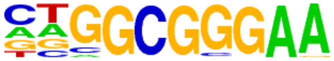 | E2F1(E2F)/Hela-E2F1-ChIP-Seq(GSE22478)/Homer | 1e-6 | -1.415e+01 | 0.0000 | 103.0 | 10.60% | 2912.6 | 6.44% |
| 55 | 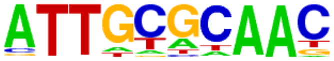 | CEBP(bZIP)/ThioMac-CEBPb-ChIP-Seq(GSE21512)/Homer | 1e-6 | -1.391e+01 | 0.0000 | 49.0 | 5.04% | 1061.5 | 2.35% |
| 56 | 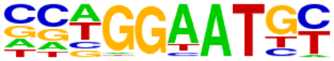 | TEAD4(TEA)/Tropoblast-Tead4-ChIP-Seq(GSE37350)/Homer | 1e-6 | -1.386e+01 | 0.0000 | 89.0 | 9.16% | 2422.2 | 5.36% |
| 57 | 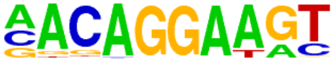 | Ets1-distal(ETS)/CD4+-PolII-ChIP-Seq(Barski_et_al.)/Homer | 1e-5 | -1.298e+01 | 0.0000 | 45.0 | 4.63% | 971.3 | 2.15% |
| 58 | 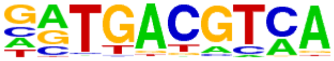 | Atf1(bZIP)/K562-ATF1-ChIP-Seq(GSE31477)/Homer | 1e-5 | -1.273e+01 | 0.0000 | 82.0 | 8.44% | 2239.8 | 4.95% |
| 59 | 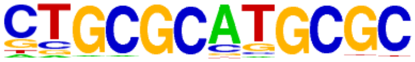 | NRF1(NRF)/MCF7-NRF1-ChIP-Seq(Unpublished)/Homer | 1e-5 | -1.261e+01 | 0.0000 | 80.0 | 8.23% | 2175.8 | 4.81% |
| 60 | 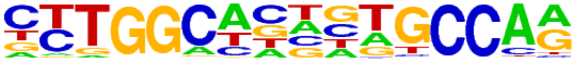 | NF1(CTF)/LNCAP-NF1-ChIP-Seq(Unpublished)/Homer | 1e-5 | -1.243e+01 | 0.0000 | 69.0 | 7.10% | 1796.5 | 3.97% |
| 61 | 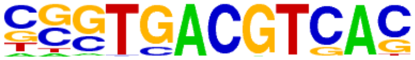 | CRE(bZIP)/Promoter/Homer | 1e-5 | -1.174e+01 | 0.0000 | 60.0 | 6.17% | 1521.6 | 3.36% |
| 62 | 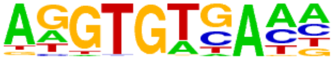 | Tbet(T-box)/CD8-Tbet-ChIP-Seq(GSE33802)/Homer | 1e-4 | -1.118e+01 | 0.0001 | 79.0 | 8.13% | 2228.9 | 4.93% |
| 63 | 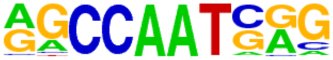 | NFY(CCAAT)/Promoter/Homer | 1e-4 | -1.111e+01 | 0.0001 | 108.0 | 11.11% | 3320.7 | 7.34% |
| 64 | 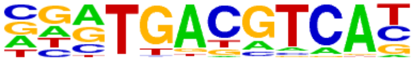 | Atf7(bZIP)/3T3L1-Atf7-ChIP-Seq(GSE56872)/Homer | 1e-4 | -1.087e+01 | 0.0001 | 60.0 | 6.17% | 1569.0 | 3.47% |
| 65 | | STAT1(Stat)/HelaS3-STAT1-ChIP-Seq(GSE12782)/Homer | 1e-4 | -1.015e+01 | 0.0002 | 33.0 | 3.40% | 703.4 | 1.56% |

| | | | | | | | | | |
|---|---|---|---|---|---|---|---|---|---|
| | 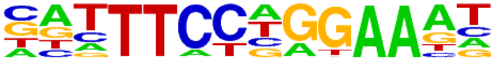 | | | | | | | | |
| 66 | 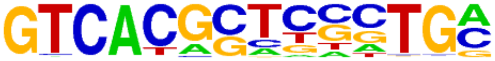 | PAX5(Paired,Homeobox),condensed/GM12878-PAX5-ChIP-Seq(GSE32465)/Homer | 1e-4 | -9.957e+00 | 0.0002 | 26.0 | 2.67% | 498.7 | 1.10% |
| 67 | 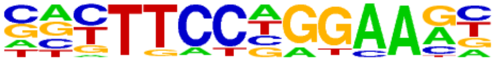 | Stat3+il21(Stat)/CD4-Stat3-ChIP-Seq(GSE19198)/Homer | 1e-4 | -9.703e+00 | 0.0003 | 81.0 | 8.33% | 2405.0 | 5.32% |
| 68 | 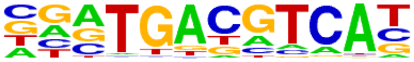 | Atf2(bZIP)/3T3L1-Atf2-ChIP-Seq(GSE56872)/Homer | 1e-4 | -9.621e+00 | 0.0003 | 45.0 | 4.63% | 1117.6 | 2.47% |
| 69 | 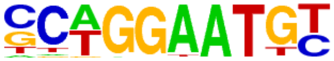 | TEAD2(TEA)/Py2T-Tead2-ChIP-Seq(GSE55709)/Homer | 1e-4 | -9.578e+00 | 0.0003 | 49.0 | 5.04% | 1256.2 | 2.78% |
| 70 | 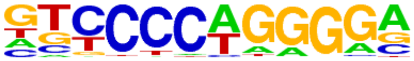 | EBF1(EBF)/Near-E2A-ChIP-Seq(GSE21512)/Homer | 1e-4 | -9.559e+00 | 0.0003 | 196.0 | 20.16% | 7027.1 | 15.54% |
| 71 | 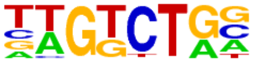 | Smad3(MAD)/NPC-Smad3-ChIP-Seq(GSE36673)/Homer | 1e-4 | -9.488e+00 | 0.0003 | 274.0 | 28.19% | 10362.3 | 22.91% |
| 72 | 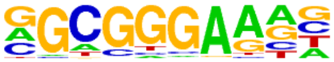 | E2F6(E2F)/Hela-E2F6-ChIP-Seq(GSE31477)/Homer | 1e-3 | -8.946e+00 | 0.0006 | 162.0 | 16.67% | 5689.9 | 12.58% |
| 73 | 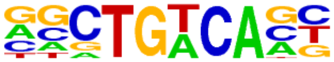 | Meis1(Homeobox)/MastCells-Meis1-ChIP-Seq(GSE48085)/Homer | 1e-3 | -8.901e+00 | 0.0006 | 164.0 | 16.87% | 5777.7 | 12.78% |
| 74 | 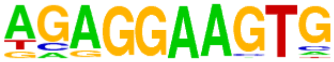 | PU.1(ETS)/ThioMac-PU.1-ChIP-Seq(GSE21512)/Homer | 1e-3 | -8.869e+00 | 0.0006 | 63.0 | 6.48% | 1797.6 | 3.98% |
| 75 | 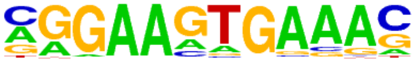 | PU.1-IRF(ETS:IRF)/Bcell-PU.1-ChIP-Seq(GSE21512)/Homer | 1e-3 | -8.563e+00 | 0.0008 | 127.0 | 13.07% | 4304.5 | 9.52% |
| 76 | 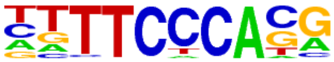 | Rbpj1(?)/Panc1-Rbpj1-ChIP-Seq(GSE47459)/Homer | 1e-3 | -8.473e+00 | 0.0009 | 151.0 | 15.53% | 5292.8 | 11.70% |
| 77 | 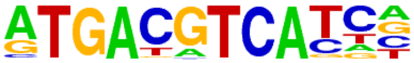 | JunD(bZIP)/K562-JunD-ChIP-Seq/Homer | 1e-3 | -8.232e+00 | 0.0011 | 21.0 | 2.16% | 405.0 | 0.90% |
| 78 | 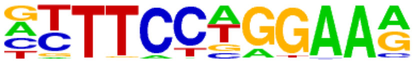 | STAT4(Stat)/CD4-Stat4-ChIP-Seq(GSE22104)/Homer | 1e-3 | -8.029e+00 | 0.0013 | 79.0 | 8.13% | 2461.1 | 5.44% |
| 79 | 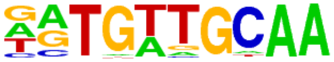 | CEBP:AP1(bZIP)/ThioMac-CEBPb-ChIP-Seq(GSE21512)/Homer | 1e-3 | -7.767e+00 | 0.0017 | 46.0 | 4.73% | 1253.3 | 2.77% |
| 80 | 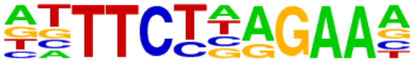 | STAT5(Stat)/mCD4+-Stat5-ChIP-Seq(GSE12346)/Homer | 1e-3 | -7.735e+00 | 0.0017 | 34.0 | 3.50% | 838.8 | 1.85% |
| 81 | 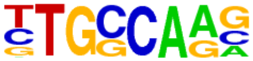 | NF1-halfsite(CTF)/LNCaP-NF1-ChIP-Seq(Unpublished)/Homer | 1e-3 | -7.724e+00 | 0.0017 | 203.0 | 20.88% | 7571.9 | 16.74% |
| 82 | | GRHL2(CP2)/HBE-GRHL2-ChIP-Seq(GSE46194)/Homer | 1e-3 | -7.221e+00 | 0.0028 | 29.0 | 2.98% | 695.2 | 1.54% |

| | | | | | | | | | |
|---|---|---|---|---|---|---|---|---|---|
| | 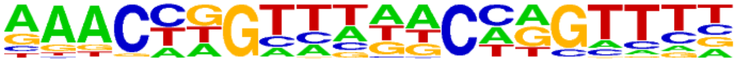 | | | | | | | | |
| 83 | 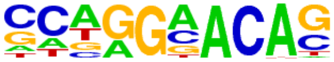 | AR-halfsite(NR)/LNCaP-AR-ChIP-Seq(GSE27824)/Homer | 1e-3 | -7.211e+00 | 0.0028 | 358.0 | 36.83% | 14458.3 | 31.97% |
| 84 | 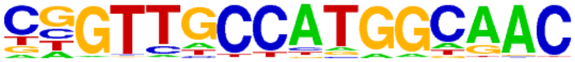 | RFX(HTH)/K562-RFX3-ChIP-Seq(SRA012198)/Homer | 1e-3 | -7.157e+00 | 0.0030 | 25.0 | 2.57% | 567.3 | 1.25% |
| 85 | 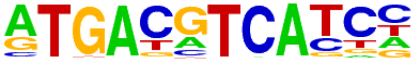 | c-Jun-CRE(bZIP)/K562-cJun-ChIP-Seq(GSE31477)/Homer | 1e-3 | -6.987e+00 | 0.0035 | 34.0 | 3.50% | 876.6 | 1.94% |
| 86 | 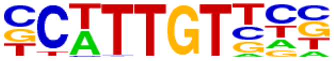 | Sox10(HMG)/SciaticNerve-Sox3-ChIP-Seq(GSE35132)/Homer | 1e-2 | -6.849e+00 | 0.0039 | 114.0 | 11.73% | 3968.4 | 8.78% |
| 87 | 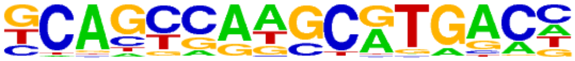 | PAX5(Paired,Homeobox)/GM12878-PAX5-ChIP-Seq(GSE32465)/Homer | 1e-2 | -6.520e+00 | 0.0054 | 59.0 | 6.07% | 1821.5 | 4.03% |
| 88 | 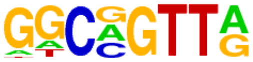 | MYB(HTH)/ERMYB-Myb-ChIPSeq(GSE22095)/Homer | 1e-2 | -6.465e+00 | 0.0056 | 180.0 | 18.52% | 6780.1 | 14.99% |
| 89 | 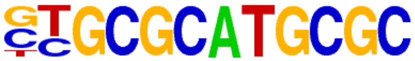 | NRF(NRF)/Promoter/Homer | 1e-2 | -6.116e+00 | 0.0079 | 68.0 | 7.00% | 2204.4 | 4.87% |
| 90 | 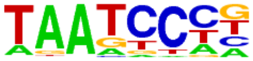 | Pitx1(Homeobox)/Chicken-Pitx1-ChIP-Seq(GSE38910)/Homer | 1e-2 | -6.031e+00 | 0.0085 | 259.0 | 26.65% | 10284.2 | 22.74% |
| 91 | 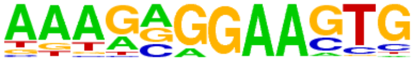 | SpiB(ETS)/OCILY3-SPIB-ChIP-Seq(GSE56857)/Homer | 1e-2 | -6.028e+00 | 0.0085 | 30.0 | 3.09% | 789.7 | 1.75% |
| 92 | 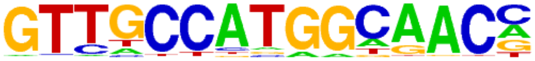 | Rfx2(HTH)/LoVo-RFX2-ChIP-Seq(GSE49402)/Homer | 1e-2 | -5.957e+00 | 0.0090 | 24.0 | 2.47% | 587.2 | 1.30% |
| 93 | 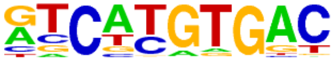 | MITF(bHLH)/MastCells-MITF-ChIP-Seq(GSE48085)/Homer | 1e-2 | -5.782e+00 | 0.0106 | 93.0 | 9.57% | 3238.4 | 7.16% |
| 94 | 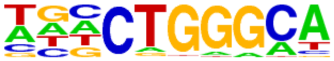 | ZNF416(Zf)/HEK293-ZNF416.GFP-ChIP-Seq(GSE58341)/Homer | 1e-2 | -5.747e+00 | 0.0108 | 162.0 | 16.67% | 6125.2 | 13.54% |
| 95 | 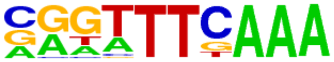 | CHR(?)/Hela-CellCycle-Expression/Homer | 1e-2 | -5.690e+00 | 0.0113 | 33.0 | 3.40% | 915.6 | 2.02% |
| 96 | 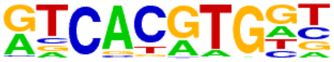 | Usf2(bHLH)/C2C12-Usf2-ChIP-Seq(GSE36030)/Homer | 1e-2 | -5.496e+00 | 0.0136 | 39.0 | 4.01% | 1147.3 | 2.54% |
| 97 | 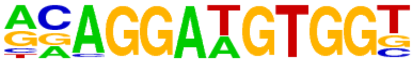 | ETS:RUNX(ETS,Runt)/Jurkat-RUNX1-ChIP-Seq(GSE17954)/Homer | 1e-2 | -5.477e+00 | 0.0138 | 21.0 | 2.16% | 509.3 | 1.13% |
| 98 | 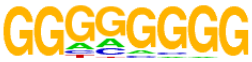 | Maz(Zf)/HepG2-Maz-ChIP-Seq(GSE31477)/Homer | 1e-2 | -5.398e+00 | 0.0147 | 305.0 | 31.38% | 12456.9 | 27.55% |
| 99 | | Bcl6(Zf)/Liver-Bcl6-ChIP-Seq(GSE31578)/Homer | 1e-2 | -5.212e+00 | 0.0176 | 110.0 | 11.32% | 4011.1 | 8.87% |

| | | | | | | | | | |
|---|---|---|---|---|---|---|---|---|---|
| | 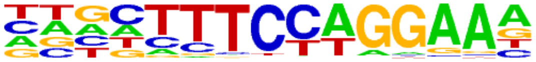 | | | | | | | | |
| 100 | 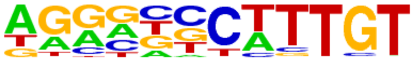 | Sox9(HMG)/Limb-SOX9-ChIP-Seq(GSE73225)/Homer | 1e-2 | -5.164e+00 | 0.0182 | 66.0 | 6.79% | 2219.6 | 4.91% |
| 101 | 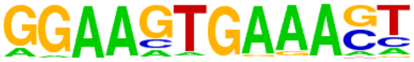 | PU.1:IRF8(ETS:IRF)/pDC-Irf8-ChIP-Seq(GSE66899)/Homer | 1e-2 | -5.090e+00 | 0.0195 | 19.0 | 1.95% | 460.1 | 1.02% |
| 102 | 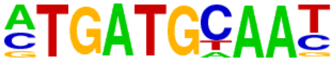 | Atf4(bZIP)/MEF-Atf4-ChIP-Seq(GSE35681)/Homer | 1e-2 | -5.063e+00 | 0.0198 | 18.0 | 1.85% | 428.8 | 0.95% |
| 103 | 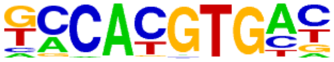 | NPAS2(bHLH)/Liver-NPAS2-ChIP-Seq(GSE39860)/Homer | 1e-2 | -5.043e+00 | 0.0200 | 115.0 | 11.83% | 4244.3 | 9.39% |
| 104 | 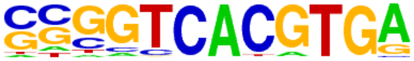 | E-box(bHLH)/Promoter/Homer | 1e-2 | -5.039e+00 | 0.0200 | 21.0 | 2.16% | 530.4 | 1.17% |
| 105 | 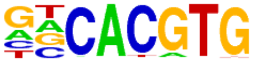 | BMAL1(bHLH)/Liver-Bmal1-ChIP-Seq(GSE39860)/Homer | 1e-2 | -4.975e+00 | 0.0210 | 153.0 | 15.74% | 5867.2 | 12.97% |
| 106 | 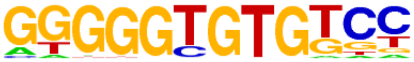 | KLF10(Zf)/HEK293-KLF10.GFP-ChIP-Seq(GSE58341)/Homer | 1e-2 | -4.751e+00 | 0.0260 | 96.0 | 9.88% | 3492.8 | 7.72% |
| 107 | 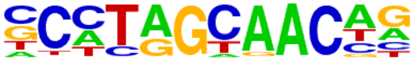 | Rfx5(HTH)/GM12878-Rfx5-ChIP-Seq(GSE31477)/Homer | 1e-2 | -4.745e+00 | 0.0260 | 37.0 | 3.81% | 1128.5 | 2.50% |
| 108 | 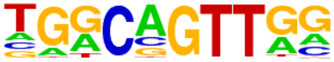 | AMYB(HTH)/Testes-AMYB-ChIP-Seq(GSE44588)/Homer | 1e-2 | -4.722e+00 | 0.0263 | 141.0 | 14.51% | 5396.5 | 11.93% |
| 109 | 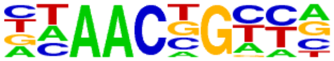 | BMYB(HTH)/Hela-BMYB-ChIP-Seq(GSE27030)/Homer | 1e-2 | -4.627e+00 | 0.0286 | 131.0 | 13.48% | 4985.2 | 11.02% |

### Homer Known Motif Enrichment Results (QC/homer/01_041O_SCC-13_BNC1_hg38)

[Homer *de novo* Motif Results](#)
[Gene Ontology Enrichment Results](#)
[Known Motif Enrichment Results (txt file)](#)
Total Target Sequences = 2341, Total Background Sequences = 46000

| Rank | Motif | Name | P-value | log P-pvalue | q-value (Benjamini) | # Target Sequences with Motif | % of Targets Sequences with Motif | # Background Sequences with Motif | % of Background Sequences with Motif |
|---|---|---|---|---|---|---|---|---|---|
| 1 | 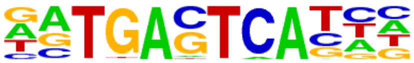 | Atf3(bZIP)/GBM-ATF3-ChIP-Seq(GSE33912)/Homer | 1e-431 | -9.940e+02 | 0.0000 | 704.0 | 30.07% | 1600.8 | 3.48% |
| 2 | 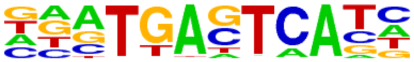 | Fra1(bZIP)/BT549-Fra1-ChIP-Seq(GSE46166)/Homer | 1e-424 | -9.775e+02 | 0.0000 | 657.0 | 28.06% | 1360.8 | 2.96% |
| 3 | 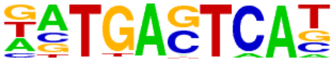 | BATF(bZIP)/Th17-BATF-ChIP-Seq(GSE39756)/Homer | 1e-411 | -9.470e+02 | 0.0000 | 686.0 | 29.30% | 1607.0 | 3.50% |
| 4 | 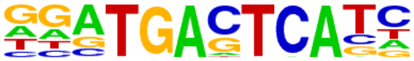 | Fra2(bZIP)/Striatum-Fra2-ChIP-Seq(GSE43429)/Homer | 1e-410 | -9.461e+02 | 0.0000 | 632.0 | 27.00% | 1288.8 | 2.81% |
| 5 | 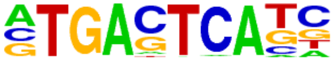 | AP-1(bZIP)/ThioMac-PU.1-ChIP-Seq(GSE21512)/Homer | 1e-407 | -9.390e+02 | 0.0000 | 739.0 | 31.57% | 1979.0 | 4.31% |
| 6 | 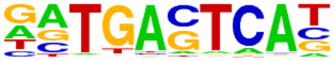 | JunB(bZIP)/DendriticCells-Junb-ChIP-Seq(GSE36099)/Homer | 1e-403 | -9.302e+02 | 0.0000 | 649.0 | 27.72% | 1423.4 | 3.10% |
| 7 | 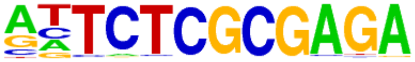 | GFX(?)/Promoter/Homer | 1e-376 | -8.677e+02 | 0.0000 | 284.0 | 12.13% | 106.9 | 0.23% |
| 8 | 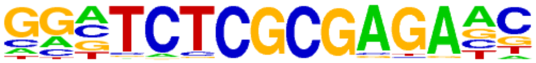 | ZBTB33(Zf)/GM12878-ZBTB33-ChIP-Seq(GSE32465)/Homer | 1e-371 | -8.565e+02 | 0.0000 | 387.0 | 16.53% | 350.6 | 0.76% |
| 9 | 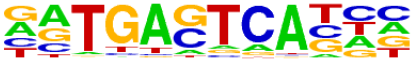 | Fosl2(bZIP)/3T3L1-Fosl2-ChIP-Seq(GSE56872)/Homer | 1e-363 | -8.372e+02 | 0.0000 | 538.0 | 22.98% | 1007.2 | 2.19% |
| 10 | 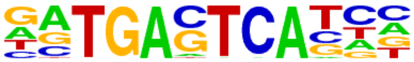 | Jun-AP1(bZIP)/K562-cJun-ChIP-Seq(GSE31477)/Homer | 1e-306 | -7.064e+02 | 0.0000 | 439.0 | 18.75% | 760.2 | 1.65% |
| 11 | 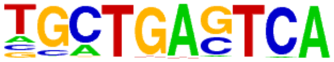 | Bach2(bZIP)/OCILy7-Bach2-ChIP-Seq(GSE44420)/Homer | 1e-179 | -4.126e+02 | 0.0000 | 306.0 | 13.07% | 687.4 | 1.50% |
| 12 | 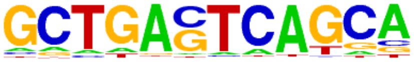 | MafK(bZIP)/C2C12-MafK-ChIP-Seq(GSE36030)/Homer | 1e-56 | -1.310e+02 | 0.0000 | 177.0 | 7.56% | 800.0 | 1.74% |
| 13 | 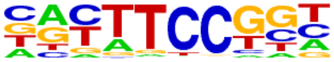 | Fli1(ETS)/CD8-FLI-ChIP-Seq(GSE20898)/Homer | 1e-54 | -1.253e+02 | 0.0000 | 585.0 | 24.99% | 5971.6 | 13.00% |
| 14 | | Nrf2(bZIP)/Lymphoblast-Nrf2-ChIP-Seq(GSE37589)/Homer | 1e-53 | -1.227e+02 | 0.0000 | 79.0 | 3.37% | 139.8 | 0.30% |

| | | | | | | | | | |
|---|---|---|---|---|---|---|---|---|---|
| | 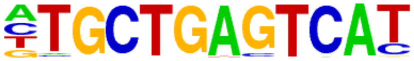 | | | | | | | | |
| 15 | 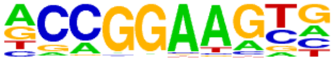 | ETV4(ETS)/HepG2-ETV4-ChIP-Seq(ENCODE)/Homer | 1e-50 | -1.161e+02 | 0.0000 | 613.0 | 26.19% | 6570.8 | 14.30% |
| 16 | 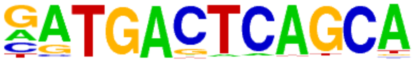 | NF-E2(bZIP)/K562-NFE2-ChIP-Seq(GSE31477)/Homer | 1e-49 | -1.130e+02 | 0.0000 | 84.0 | 3.59% | 186.1 | 0.40% |
| 17 | 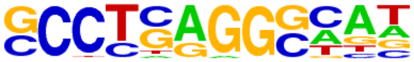 | AP-2gamma(AP2)/MCF7-TFAP2C-ChIP-Seq(GSE21234)/Homer | 1e-48 | -1.106e+02 | 0.0000 | 746.0 | 31.87% | 8790.9 | 19.14% |
| 18 | 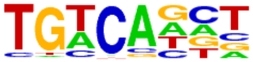 | Tgif2(Homeobox)/mES-Tgif2-ChIP-Seq(GSE55404)/Homer | 1e-45 | -1.055e+02 | 0.0000 | 967.0 | 41.31% | 12659.1 | 27.56% |
| 19 | 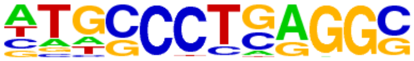 | AP-2alpha(AP2)/Hela-AP2alpha-ChIP-Seq(GSE31477)/Homer | 1e-45 | -1.051e+02 | 0.0000 | 625.0 | 26.70% | 6987.5 | 15.21% |
| 20 | 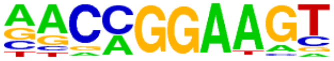 | ETV1(ETS)/GIST48-ETV1-ChIP-Seq(GSE22441)/Homer | 1e-43 | -1.005e+02 | 0.0000 | 612.0 | 26.14% | 6887.5 | 14.99% |
| 21 | 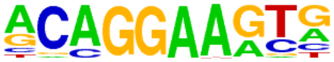 | ERG(ETS)/VCaP-ERG-ChIP-Seq(GSE14097)/Homer | 1e-43 | -1.003e+02 | 0.0000 | 644.0 | 27.51% | 7394.1 | 16.10% |
| 22 | 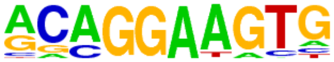 | ETS1(ETS)/Jurkat-ETS1-ChIP-Seq(GSE17954)/Homer | 1e-38 | -8.972e+01 | 0.0000 | 488.0 | 20.85% | 5219.4 | 11.36% |
| 23 | 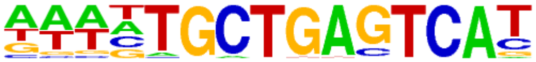 | Bach1(bZIP)/K562-Bach1-ChIP-Seq(GSE31477)/Homer | 1e-38 | -8.952e+01 | 0.0000 | 63.0 | 2.69% | 129.3 | 0.28% |
| 24 | 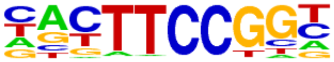 | Elk1(ETS)/Hela-Elk1-ChIP-Seq(GSE31477)/Homer | 1e-38 | -8.924e+01 | 0.0000 | 383.0 | 16.36% | 3700.7 | 8.06% |
| 25 | 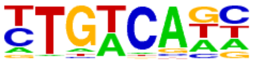 | Tgif1(Homeobox)/mES-Tgif1-ChIP-Seq(GSE55404)/Homer | 1e-37 | -8.744e+01 | 0.0000 | 857.0 | 36.61% | 11273.0 | 24.54% |
| 26 | 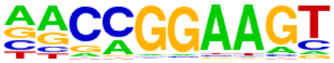 | GABPA(ETS)/Jurkat-GABPa-ChIP-Seq(GSE17954)/Homer | 1e-36 | -8.461e+01 | 0.0000 | 444.0 | 18.97% | 4668.5 | 10.16% |
| 27 | 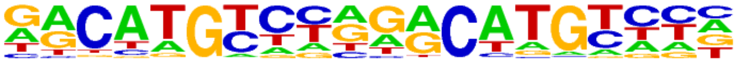 | p53(p53)/Saos-p53-ChIP-Seq(GSE15780)/Homer | 1e-36 | -8.363e+01 | 0.0000 | 80.0 | 3.42% | 250.5 | 0.55% |
| 28 | 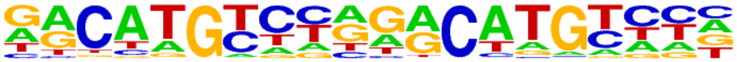 | p53(p53)/Saos-p53-ChIP-Seq/Homer | 1e-36 | -8.363e+01 | 0.0000 | 80.0 | 3.42% | 250.5 | 0.55% |
| 29 | 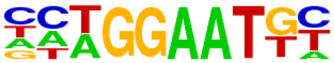 | TEAD(TEA)/Fibroblast-PU.1-ChIP-Seq(Unpublished)/Homer | 1e-36 | -8.362e+01 | 0.0000 | 216.0 | 9.23% | 1595.9 | 3.47% |
| 30 | 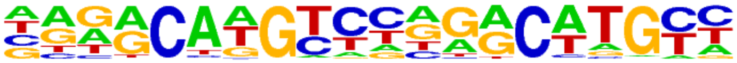 | p73(p53)/Trachea-p73-ChIP-Seq(PRJNA310161)/Homer | 1e-36 | -8.349e+01 | 0.0000 | 62.0 | 2.65% | 138.1 | 0.30% |
| 31 | | Etv2(ETS)/ES-ER71-ChIP-Seq(GSE59402)/Homer | 1e-35 | -8.181e+01 | 0.0000 | 406.0 | 17.34% | 4166.2 | 9.07% |

| | | | | | | | | | |
|---|---|---|---|---|---|---|---|---|---|
| | 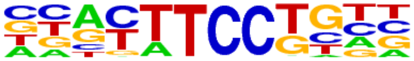 | | | | | | | | |
| 32 | 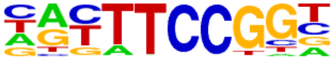 | Elk4(ETS)/Hela-Elk4-ChIP-Seq(GSE31477)/Homer | 1e-35 | -8.113e+01 | 0.0000 | 374.0 | 15.98% | 3718.2 | 8.09% |
| 33 | 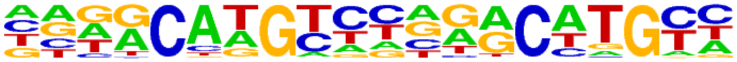 | p63(p53)/Keratinocyte-p63-ChIP-Seq(GSE17611)/Homer | 1e-35 | -8.088e+01 | 0.0000 | 195.0 | 8.33% | 1379.7 | 3.00% |
| 34 | 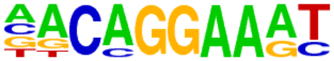 | EWS:FLI1-fusion(ETS)/SK_N_MC-EWS:FLI1-ChIP-Seq(SRA014231)/Homer | 1e-35 | -8.085e+01 | 0.0000 | 294.0 | 12.56% | 2619.0 | 5.70% |
| 35 | 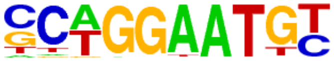 | TEAD2(TEA)/Py2T-Tead2-ChIP-Seq(GSE55709)/Homer | 1e-34 | -7.942e+01 | 0.0000 | 207.0 | 8.84% | 1536.1 | 3.34% |
| 36 | 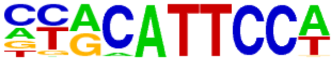 | TEAD1(TEAD)/HepG2-TEAD1-ChIP-Seq(Encode)/Homer | 1e-34 | -7.908e+01 | 0.0000 | 306.0 | 13.07% | 2807.3 | 6.11% |
| 37 | 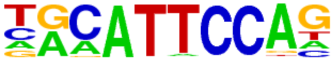 | TEAD3(TEA)/HepG2-TEAD3-ChIP-Seq(Encode)/Homer | 1e-34 | -7.843e+01 | 0.0000 | 330.0 | 14.10% | 3147.8 | 6.85% |
| 38 | 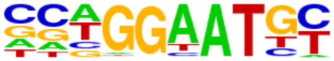 | TEAD4(TEA)/Tropoblast-Tead4-ChIP-Seq(GSE37350)/Homer | 1e-32 | -7.495e+01 | 0.0000 | 306.0 | 13.07% | 2875.4 | 6.26% |
| 39 | 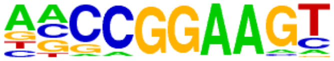 | ELF1(ETS)/Jurkat-ELF1-ChIP-Seq(SRA014231)/Homer | 1e-31 | -7.300e+01 | 0.0000 | 337.0 | 14.40% | 3342.7 | 7.28% |
| 40 | 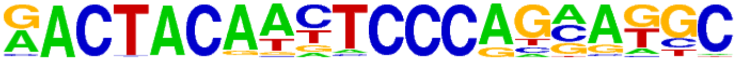 | Ronin(THAP)/ES-Thap11-ChIP-Seq(GSE51522)/Homer | 1e-29 | -6.827e+01 | 0.0000 | 75.0 | 3.20% | 276.8 | 0.60% |
| 41 | 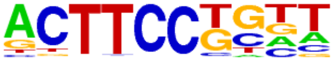 | Elf4(ETS)/BMDM-Elf4-ChIP-Seq(GSE88699)/Homer | 1e-27 | -6.342e+01 | 0.0000 | 418.0 | 17.86% | 4742.5 | 10.32% |
| 42 | 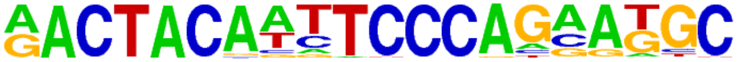 | GFY-Staf(?,Zf)/Promoter/Homer | 1e-26 | -6.035e+01 | 0.0000 | 80.0 | 3.42% | 357.5 | 0.78% |
| 43 | 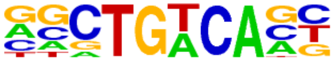 | Meis1(Homeobox)/MastCells-Meis1-ChIP-Seq(GSE48085)/Homer | 1e-25 | -5.863e+01 | 0.0000 | 603.0 | 25.76% | 7848.2 | 17.08% |
| 44 | 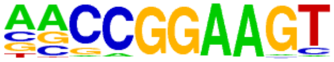 | ETS(ETS)/Promoter/Homer | 1e-25 | -5.813e+01 | 0.0000 | 221.0 | 9.44% | 2001.7 | 4.36% |
| 45 | 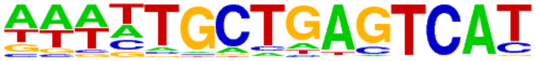 | NFE2L2(bZIP)/HepG2-NFE2L2-ChIP-Seq(Encode)/Homer | 1e-23 | -5.317e+01 | 0.0000 | 43.0 | 1.84% | 110.5 | 0.24% |
| 46 | 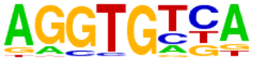 | Tbx5(T-box)/HL1-Tbx5.biotin-ChIP-Seq(GSE21529)/Homer | 1e-22 | -5.150e+01 | 0.0000 | 992.0 | 42.38% | 14989.8 | 32.63% |
| 47 | 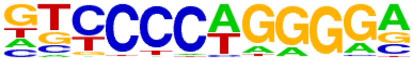 | EBF1(EBF)/Near-E2A-ChIP-Seq(GSE21512)/Homer | 1e-22 | -5.144e+01 | 0.0000 | 561.0 | 23.96% | 7380.6 | 16.07% |
| 48 | | KLF5(Zf)/LoVo-KLF5-ChIP-Seq(GSE49402)/Homer | 1e-22 | -5.129e+01 | 0.0000 | 723.0 | 30.88% | 10157.0 | 22.11% |

| | | | | | | | | | |
|---|---|---|---|---|---|---|---|---|---|
| | 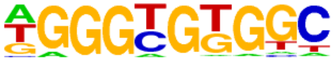 | | | | | | | | |
| 49 | 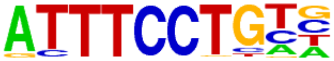 | EWS:ERG-fusion(ETS)/CADO_ES1-EWS:ERG-ChIP-Seq(SRA014231)/Homer | 1e-21 | -4.942e+01 | 0.0000 | 226.0 | 9.65% | 2219.4 | 4.83% |
| 50 | 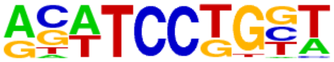 | SPDEF(ETS)/VCaP-SPDEF-ChIP-Seq(SRA014231)/Homer | 1e-21 | -4.919e+01 | 0.0000 | 363.0 | 15.51% | 4250.8 | 9.25% |
| 51 | 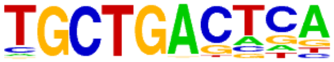 | MafA(bZIP)/Islet-MafA-ChIP-Seq(GSE30298)/Homer | 1e-20 | -4.806e+01 | 0.0000 | 315.0 | 13.46% | 3544.5 | 7.72% |
| 52 | 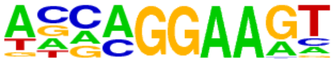 | EHF(ETS)/LoVo-EHF-ChIP-Seq(GSE49402)/Homer | 1e-20 | -4.730e+01 | 0.0000 | 417.0 | 17.81% | 5153.5 | 11.22% |
| 53 | 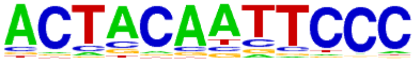 | GFY(?)/Promoter/Homer | 1e-20 | -4.619e+01 | 0.0000 | 73.0 | 3.12% | 383.9 | 0.84% |
| 54 | 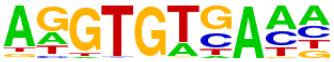 | Tbet(T-box)/CD8-Tbet-ChIP-Seq(GSE33802)/Homer | 1e-16 | -3.897e+01 | 0.0000 | 278.0 | 11.88% | 3209.0 | 6.99% |
| 55 | 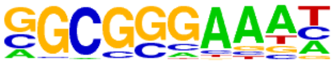 | E2F4(E2F)/K562-E2F4-ChIP-Seq(GSE31477)/Homer | 1e-16 | -3.832e+01 | 0.0000 | 311.0 | 13.28% | 3736.7 | 8.13% |
| 56 | 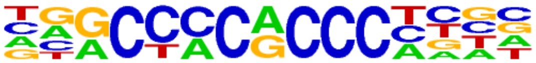 | KLF3(Zf)/MEF-Klf3-ChIP-Seq(GSE44748)/Homer | 1e-15 | -3.678e+01 | 0.0000 | 305.0 | 13.03% | 3686.4 | 8.02% |
| 57 | 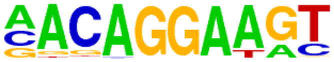 | Ets1-distal(ETS)/CD4+-PolII-ChIP-Seq(Barski_et_al.)/Homer | 1e-15 | -3.588e+01 | 0.0000 | 131.0 | 5.60% | 1167.0 | 2.54% |
| 58 | 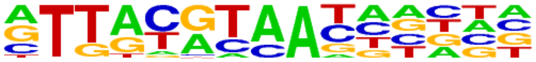 | NFIL3(bZIP)/HepG2-NFIL3-ChIP-Seq(Encode)/Homer | 1e-15 | -3.543e+01 | 0.0000 | 123.0 | 5.25% | 1069.7 | 2.33% |
| 59 | 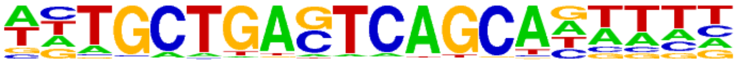 | MafB(bZIP)/BMM-Mafb-ChIP-Seq(GSE75722)/Homer | 1e-15 | -3.523e+01 | 0.0000 | 128.0 | 5.47% | 1137.7 | 2.48% |
| 60 | 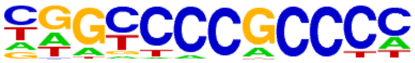 | Sp2(Zf)/HEK293-Sp2.eGFP-ChIP-Seq(Encode)/Homer | 1e-15 | -3.505e+01 | 0.0000 | 814.0 | 34.77% | 12497.0 | 27.20% |
| 61 | 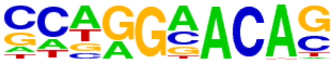 | AR-halfsite(NR)/LNCaP-AR-ChIP-Seq(GSE27824)/Homer | 1e-14 | -3.257e+01 | 0.0000 | 1070.0 | 45.71% | 17405.7 | 37.89% |
| 62 | 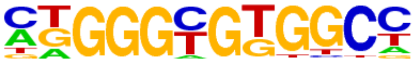 | KLF6(Zf)/PDAC-KLF6-ChIP-Seq(GSE64557)/Homer | 1e-13 | -3.044e+01 | 0.0000 | 580.0 | 24.78% | 8530.4 | 18.57% |
| 63 | 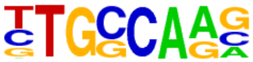 | NF1-halfsite(CTF)/LNCaP-NF1-ChIP-Seq(Unpublished)/Homer | 1e-13 | -3.014e+01 | 0.0000 | 605.0 | 25.84% | 8985.3 | 19.56% |
| 64 | 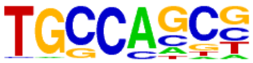 | HIC1(Zf)/Treg-ZBTB29-ChIP-Seq(GSE99889)/Homer | 1e-12 | -2.983e+01 | 0.0000 | 811.0 | 34.64% | 12718.2 | 27.69% |
| 65 | | MITF(bHLH)/MastCells-MITF-ChIP-Seq(GSE48085)/Homer | 1e-12 | -2.962e+01 | 0.0000 | 309.0 | 13.20% | 3971.3 | 8.64% |

| | | | | | | | | | |
|---|---|---|---|---|---|---|---|---|---|
| | 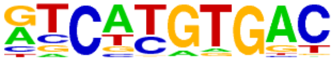 | | | | | | | | |
| 66 | 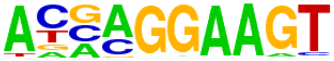 | ELF5(ETS)/T47D-ELF5-ChIP-Seq(GSE30407)/Homer | 1e-12 | -2.894e+01 | 0.0000 | 233.0 | 9.95% | 2789.5 | 6.07% |
| 67 | 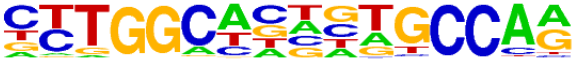 | NF1(CTF)/LNCAP-NF1-ChIP-Seq(Unpublished)/Homer | 1e-12 | -2.893e+01 | 0.0000 | 199.0 | 8.50% | 2270.2 | 4.94% |
| 68 | 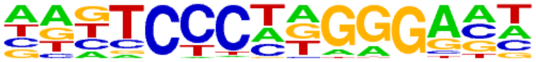 | EBF2(EBF)/BrownAdipose-EBF2-ChIP-Seq(GSE97114)/Homer | 1e-12 | -2.813e+01 | 0.0000 | 378.0 | 16.15% | 5160.8 | 11.23% |
| 69 | 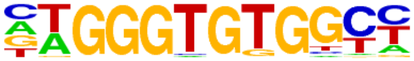 | EKLF(Zf)/Erythrocyte-Klf1-ChIP-Seq(GSE20478)/Homer | 1e-12 | -2.784e+01 | 0.0000 | 119.0 | 5.08% | 1141.4 | 2.48% |
| 70 | 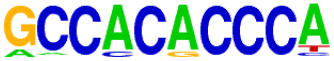 | Klf4(Zf)/mES-Klf4-ChIP-Seq(GSE11431)/Homer | 1e-11 | -2.720e+01 | 0.0000 | 215.0 | 9.18% | 2560.2 | 5.57% |
| 71 | 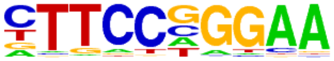 | Stat3(Stat)/mES-Stat3-ChIP-Seq(GSE11431)/Homer | 1e-11 | -2.617e+01 | 0.0000 | 176.0 | 7.52% | 1995.8 | 4.34% |
| 72 | 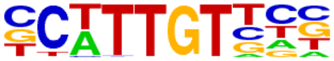 | Sox10(HMG)/SciaticNerve-Sox3-ChIP-Seq(GSE35132)/Homer | 1e-10 | -2.528e+01 | 0.0000 | 333.0 | 14.22% | 4521.0 | 9.84% |
| 73 | 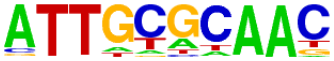 | CEBP(bZIP)/ThioMac-CEBPb-ChIP-Seq(GSE21512)/Homer | 1e-10 | -2.462e+01 | 0.0000 | 117.0 | 5.00% | 1174.4 | 2.56% |
| 74 | 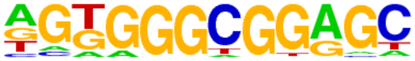 | Sp5(Zf)/mES-Sp5.Flag-ChIP-Seq(GSE72989)/Homer | 1e-10 | -2.416e+01 | 0.0000 | 533.0 | 22.77% | 8012.8 | 17.44% |
| 75 | 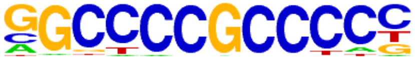 | Sp1(Zf)/Promoter/Homer | 1e-10 | -2.380e+01 | 0.0000 | 217.0 | 9.27% | 2692.9 | 5.86% |
| 76 | 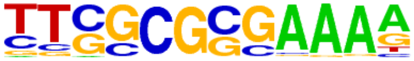 | E2F(E2F)/Hela-CellCycle-Expression/Homer | 1e-10 | -2.305e+01 | 0.0000 | 52.0 | 2.22% | 362.5 | 0.79% |
| 77 | 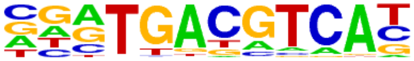 | Atf7(bZIP)/3T3L1-Atf7-ChIP-Seq(GSE56872)/Homer | 1e-9 | -2.285e+01 | 0.0000 | 160.0 | 6.83% | 1840.8 | 4.01% |
| 78 | 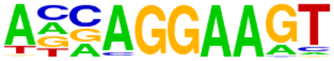 | ELF3(ETS)/PDAC-ELF3-ChIP-Seq(GSE64557)/Homer | 1e-9 | -2.205e+01 | 0.0000 | 209.0 | 8.93% | 2621.4 | 5.71% |
| 79 | 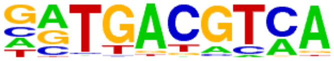 | Atf1(bZIP)/K562-ATF1-ChIP-Seq(GSE31477)/Homer | 1e-9 | -2.140e+01 | 0.0000 | 198.0 | 8.46% | 2468.1 | 5.37% |
| 80 | 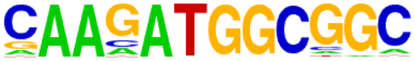 | YY1(Zf)/Promoter/Homer | 1e-9 | -2.085e+01 | 0.0000 | 58.0 | 2.48% | 458.3 | 1.00% |
| 81 | 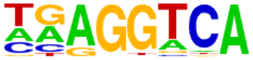 | THRb(NR)/Liver-NR1A2-ChIP-Seq(GSE52613)/Homer | 1e-8 | -2.009e+01 | 0.0000 | 1063.0 | 45.41% | 18095.8 | 39.39% |
| 82 | | ZNF143\|STAF(Zf)/CUTLL-ZNF143-ChIP-Seq(GSE29600)/Homer | 1e-8 | -1.962e+01 | 0.0000 | 142.0 | 6.07% | 1654.3 | 3.60% |

| | | | | | | | | | |
|---|---|---|---|---|---|---|---|---|---|
| | 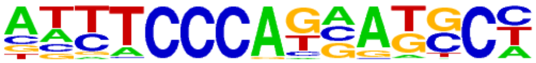 | | | | | | | | |
| 83 | 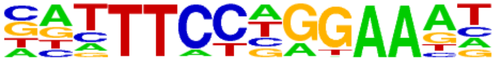 | STAT1(Stat)/HelaS3-STAT1-ChIP-Seq(GSE12782)/Homer | 1e-8 | -1.940e+01 | 0.0000 | 73.0 | 3.12% | 671.0 | 1.46% |
| 84 | 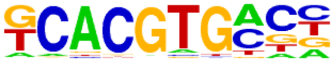 | bHLHE41(bHLH)/proB-Bhlhe41-ChIP-Seq(GSE93764)/Homer | 1e-8 | -1.866e+01 | 0.0000 | 447.0 | 19.09% | 6789.2 | 14.78% |
| 85 | 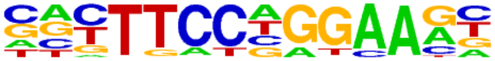 | Stat3+il21(Stat)/CD4-Stat3-ChIP-Seq(GSE19198)/Homer | 1e-7 | -1.831e+01 | 0.0000 | 194.0 | 8.29% | 2507.6 | 5.46% |
| 86 | 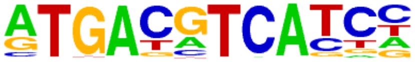 | c-Jun-CRE(bZIP)/K562-cJun-ChIP-Seq(GSE31477)/Homer | 1e-7 | -1.793e+01 | 0.0000 | 103.0 | 4.40% | 1117.7 | 2.43% |
| 87 | 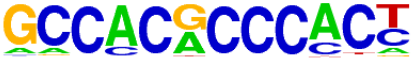 | Klf9(Zf)/GBM-Klf9-ChIP-Seq(GSE62211)/Homer | 1e-7 | -1.792e+01 | 0.0000 | 215.0 | 9.18% | 2861.2 | 6.23% |
| 88 | 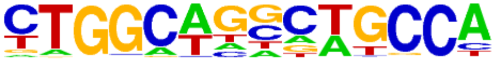 | Tlx?(NR)/NPC-H3K4me1-ChIP-Seq(GSE16256)/Homer | 1e-7 | -1.782e+01 | 0.0000 | 173.0 | 7.39% | 2189.4 | 4.77% |
| 89 | 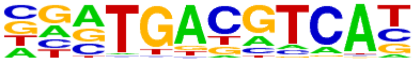 | Atf2(bZIP)/3T3L1-Atf2-ChIP-Seq(GSE56872)/Homer | 1e-7 | -1.730e+01 | 0.0000 | 118.0 | 5.04% | 1354.6 | 2.95% |
| 90 | 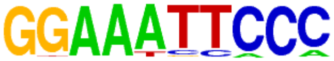 | NFkB-p65-Rel(RHD)/ThioMac-LPS-Expression(GSE23622)/Homer | 1e-7 | -1.725e+01 | 0.0000 | 31.0 | 1.32% | 189.5 | 0.41% |
| 91 | 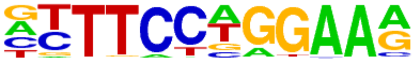 | STAT4(Stat)/CD4-Stat4-ChIP-Seq(GSE22104)/Homer | 1e-7 | -1.719e+01 | 0.0000 | 193.0 | 8.24% | 2531.5 | 5.51% |
| 92 | 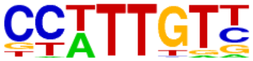 | Sox3(HMG)/NPC-Sox3-ChIP-Seq(GSE33059)/Homer | 1e-7 | -1.716e+01 | 0.0000 | 309.0 | 13.20% | 4467.5 | 9.72% |
| 93 | 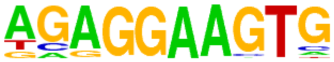 | PU.1(ETS)/ThioMac-PU.1-ChIP-Seq(GSE21512)/Homer | 1e-7 | -1.618e+01 | 0.0000 | 156.0 | 6.66% | 1974.0 | 4.30% |
| 94 | 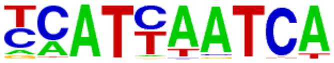 | Pdx1(Homeobox)/Islet-Pdx1-ChIP-Seq(SRA008281)/Homer | 1e-6 | -1.603e+01 | 0.0000 | 138.0 | 5.89% | 1696.8 | 3.69% |
| 95 | 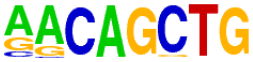 | MyoG(bHLH)/C2C12-MyoG-ChIP-Seq(GSE36024)/Homer | 1e-6 | -1.599e+01 | 0.0000 | 402.0 | 17.17% | 6147.9 | 13.38% |
| 96 | 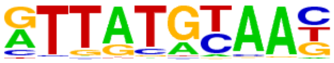 | HLF(bZIP)/HSC-HLF.Flag-ChIP-Seq(GSE69817)/Homer | 1e-6 | -1.582e+01 | 0.0000 | 99.0 | 4.23% | 1109.2 | 2.41% |
| 97 | 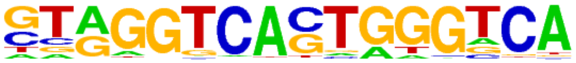 | Reverb(NR),DR2/RAW-Reverba.biotin-ChIP-Seq(GSE45914)/Homer | 1e-6 | -1.423e+01 | 0.0000 | 68.0 | 2.90% | 697.2 | 1.52% |
| 98 | 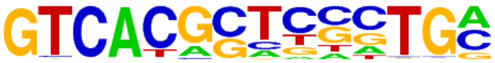 | PAX5(Paired,Homeobox),condensed/GM12878-PAX5-ChIP-Seq(GSE32465)/Homer | 1e-6 | -1.409e+01 | 0.0000 | 59.0 | 2.52% | 575.7 | 1.25% |
| 99 | | E2F1(E2F)/Hela-E2F1-ChIP-Seq(GSE22478)/Homer | 1e-5 | -1.336e+01 | 0.0000 | 152.0 | 6.49% | 2009.7 | 4.37% |

| | | | | | | | | | |
|---|---|---|---|---|---|---|---|---|---|
| | 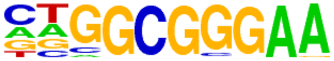 | | | | | | | | |
| 100 | 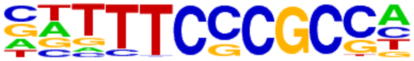 | E2F7(E2F)/Hela-E2F7-ChIP-Seq(GSE32673)/Homer | 1e-5 | -1.333e+01 | 0.0000 | 84.0 | 3.59% | 949.3 | 2.07% |
| 101 | 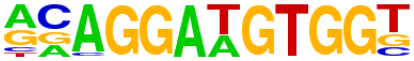 | ETS:RUNX(ETS,Runt)/Jurkat-RUNX1-ChIP-Seq(GSE17954)/Homer | 1e-5 | -1.315e+01 | 0.0000 | 59.0 | 2.52% | 593.6 | 1.29% |
| 102 | 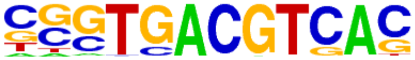 | CRE(bZIP)/Promoter/Homer | 1e-5 | -1.311e+01 | 0.0000 | 111.0 | 4.74% | 1366.5 | 2.97% |
| 103 | 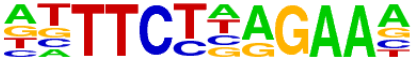 | STAT5(Stat)/mCD4+-Stat5-ChIP-Seq(GSE12346)/Homer | 1e-5 | -1.307e+01 | 0.0000 | 75.0 | 3.20% | 823.4 | 1.79% |
| 104 | 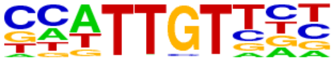 | Sox6(HMG)/Myotubes-Sox6-ChIP-Seq(GSE32627)/Homer | 1e-5 | -1.301e+01 | 0.0000 | 254.0 | 10.85% | 3732.8 | 8.13% |
| 105 | 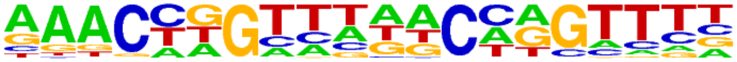 | GRHL2(CP2)/HBE-GRHL2-ChIP-Seq(GSE46194)/Homer | 1e-5 | -1.284e+01 | 0.0000 | 76.0 | 3.25% | 843.2 | 1.84% |
| 106 | 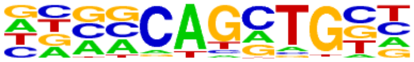 | Ascl1(bHLH)/NeuralTubes-Ascl1-ChIP-Seq(GSE55840)/Homer | 1e-5 | -1.281e+01 | 0.0000 | 589.0 | 25.16% | 9748.0 | 21.22% |
| 107 | 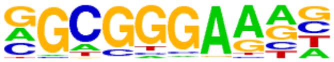 | E2F6(E2F)/Hela-E2F6-ChIP-Seq(GSE31477)/Homer | 1e-5 | -1.234e+01 | 0.0000 | 267.0 | 11.41% | 3992.3 | 8.69% |
| 108 | 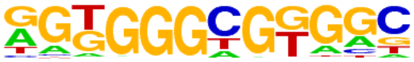 | KLF14(Zf)/HEK293-KLF14.GFP-ChIP-Seq(GSE58341)/Homer | 1e-5 | -1.221e+01 | 0.0000 | 779.0 | 33.28% | 13348.3 | 29.06% |
| 109 | 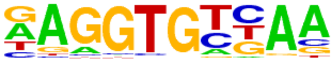 | Tbx6(T-box)/ESC-Tbx6-ChIP-Seq(GSE93524)/Homer | 1e-5 | -1.217e+01 | 0.0000 | 274.0 | 11.70% | 4123.2 | 8.98% |
| 110 | 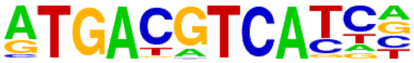 | JunD(bZIP)/K562-JunD-ChIP-Seq/Homer | 1e-5 | -1.196e+01 | 0.0000 | 43.0 | 1.84% | 397.9 | 0.87% |
| 111 | 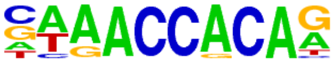 | RUNX(Runt)/HPC7-Runx1-ChIP-Seq(GSE22178)/Homer | 1e-5 | -1.193e+01 | 0.0000 | 188.0 | 8.03% | 2663.6 | 5.80% |
| 112 | 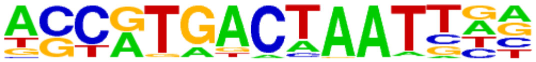 | PAX3:FKHR-fusion(Paired,Homeobox)/Rh4-PAX3:FKHR-ChIP-Seq(GSE19063)/Homer | 1e-5 | -1.183e+01 | 0.0000 | 34.0 | 1.45% | 283.3 | 0.62% |
| 113 | 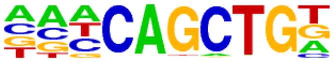 | Ap4(bHLH)/AML-Tfap4-ChIP-Seq(GSE45738)/Homer | 1e-5 | -1.175e+01 | 0.0000 | 417.0 | 17.81% | 6688.6 | 14.56% |
| 114 | 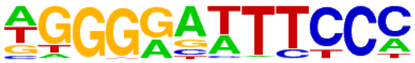 | NFkB-p65(RHD)/GM12787-p65-ChIP-Seq(GSE19485)/Homer | 1e-5 | -1.168e+01 | 0.0000 | 145.0 | 6.19% | 1960.2 | 4.27% |
| 115 | 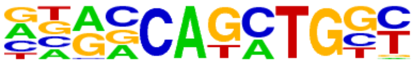 | Atoh1(bHLH)/Cerebellum-Atoh1-ChIP-Seq(GSE22111)/Homer | 1e-4 | -1.145e+01 | 0.0000 | 354.0 | 15.12% | 5578.4 | 12.14% |
| 116 | | NeuroG2(bHLH)/Fibroblast-NeuroG2-ChIP-Seq(GSE75910)/Homer | 1e-4 | -1.099e+01 | 0.0001 | 429.0 | 18.33% | 6961.6 | 15.15% |

| | | | | | | | | | |
|---|---|---|---|---|---|---|---|---|---|
| | 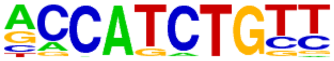 | | | | | | | | |
| 117 | 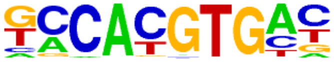 | NPAS2(bHLH)/Liver-NPAS2-ChIP-Seq(GSE39860)/Homer | 1e-4 | -1.060e+01 | 0.0001 | 336.0 | 14.35% | 5314.2 | 11.57% |
| 118 | 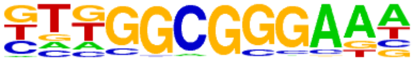 | E2F3(E2F)/MEF-E2F3-ChIP-Seq(GSE71376)/Homer | 1e-4 | -1.055e+01 | 0.0001 | 301.0 | 12.86% | 4695.7 | 10.22% |
| 119 | 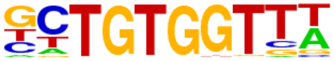 | RUNX-AML(Runt)/CD4+-PolII-ChIP-Seq(Barski_et_al.)/Homer | 1e-4 | -1.023e+01 | 0.0001 | 180.0 | 7.69% | 2609.1 | 5.68% |
| 120 | 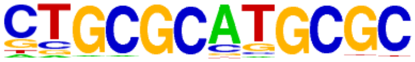 | NRF1(NRF)/MCF7-NRF1-ChIP-Seq(Unpublished)/Homer | 1e-4 | -1.015e+01 | 0.0001 | 108.0 | 4.61% | 1418.3 | 3.09% |
| 121 | 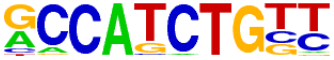 | NeuroD1(bHLH)/Islet-NeuroD1-ChIP-Seq(GSE30298)/Homer | 1e-4 | -1.009e+01 | 0.0001 | 258.0 | 11.02% | 3966.4 | 8.63% |
| 122 | 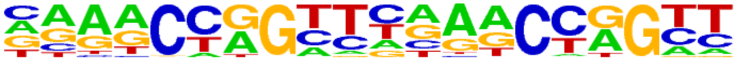 | Tcfcp2l1(CP2)/mES-Tcfcp2l1-ChIP-Seq(GSE11431)/Homer | 1e-4 | -1.002e+01 | 0.0002 | 53.0 | 2.26% | 574.8 | 1.25% |
| 123 | 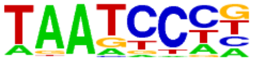 | Pitx1(Homeobox)/Chicken-Pitx1-ChIP-Seq(GSE38910)/Homer | 1e-4 | -9.833e+00 | 0.0002 | 639.0 | 27.30% | 10940.5 | 23.82% |
| 124 | 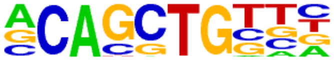 | Ptf1a(bHLH)/Panc1-Ptf1a-ChIP-Seq(GSE47459)/Homer | 1e-4 | -9.770e+00 | 0.0002 | 884.0 | 37.76% | 15589.8 | 33.94% |
| 125 | 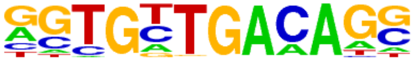 | Tbx20(T-box)/Heart-Tbx20-ChIP-Seq(GSE29636)/Homer | 1e-4 | -9.763e+00 | 0.0002 | 74.0 | 3.16% | 895.4 | 1.95% |
| 126 | 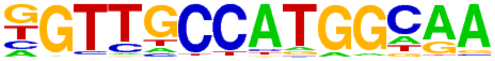 | Rfx1(HTH)/NPC-H3K4me1-ChIP-Seq(GSE16256)/Homer | 1e-4 | -9.660e+00 | 0.0002 | 71.0 | 3.03% | 852.5 | 1.86% |
| 127 | 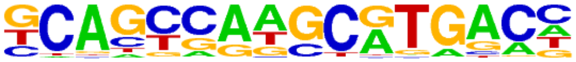 | PAX5(Paired,Homeobox)/GM12878-PAX5-ChIP-Seq(GSE32465)/Homer | 1e-4 | -9.480e+00 | 0.0003 | 139.0 | 5.94% | 1954.5 | 4.25% |
| 128 | 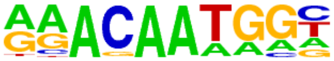 | Sox15(HMG)/CPA-Sox15-ChIP-Seq(GSE62909)/Homer | 1e-4 | -9.237e+00 | 0.0003 | 165.0 | 7.05% | 2404.9 | 5.24% |
| 129 | 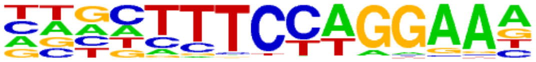 | Bcl6(Zf)/Liver-Bcl6-ChIP-Seq(GSE31578)/Homer | 1e-3 | -9.188e+00 | 0.0003 | 291.0 | 12.43% | 4609.4 | 10.03% |
| 130 | 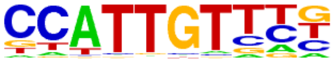 | Sox17(HMG)/Endoderm-Sox17-ChIP-Seq(GSE61475)/Homer | 1e-3 | -9.043e+00 | 0.0004 | 126.0 | 5.38% | 1757.1 | 3.82% |
| 131 | 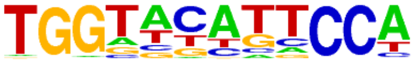 | PRDM10(Zf)/HEK293-PRDM10.eGFP-ChIP-Seq(Encode)/Homer | 1e-3 | -9.002e+00 | 0.0004 | 164.0 | 7.01% | 2399.7 | 5.22% |
| 132 | 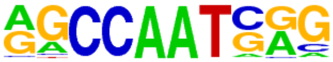 | NFY(CCAAT)/Promoter/Homer | 1e-3 | -8.853e+00 | 0.0005 | 200.0 | 8.54% | 3027.3 | 6.59% |
| 133 | | Rfx2(HTH)/LoVo-RFX2-ChIP-Seq(GSE49402)/Homer | 1e-3 | -8.789e+00 | 0.0005 | 41.0 | 1.75% | 429.9 | 0.94% |

| | | | | | | | | | |
|---|---|---|---|---|---|---|---|---|---|
| | 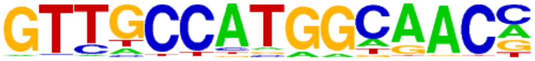 | | | | | | | | |
| 134 | 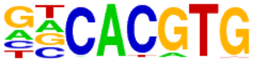 | BMAL1(bHLH)/Liver-Bmal1-ChIP-Seq(GSE39860)/Homer | 1e-3 | -8.752e+00 | 0.0005 | 461.0 | 19.69% | 7732.4 | 16.83% |
| 135 | 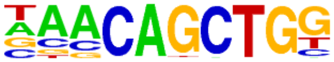 | Tcf21(bHLH)/ArterySmoothMuscle-Tcf21-ChIP-Seq(GSE61369)/Homer | 1e-3 | -8.706e+00 | 0.0005 | 322.0 | 13.75% | 5200.5 | 11.32% |
| 136 | 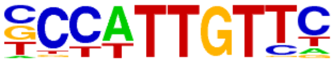 | Sox2(HMG)/mES-Sox2-ChIP-Seq(GSE11431)/Homer | 1e-3 | -8.505e+00 | 0.0006 | 149.0 | 6.36% | 2169.4 | 4.72% |
| 137 | 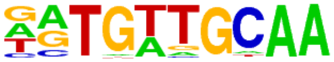 | CEBP:AP1(bZIP)/ThioMac-CEBPb-ChIP-Seq(GSE21512)/Homer | 1e-3 | -8.490e+00 | 0.0006 | 112.0 | 4.78% | 1549.6 | 3.37% |
| 138 | 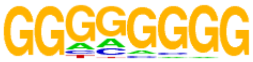 | Maz(Zf)/HepG2-Maz-ChIP-Seq(GSE31477)/Homer | 1e-3 | -8.437e+00 | 0.0007 | 607.0 | 25.93% | 10480.2 | 22.81% |
| 139 | 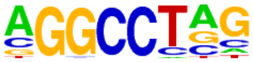 | ZNF711(Zf)/SHSY5Y-ZNF711-ChIP-Seq(GSE20673)/Homer | 1e-3 | -8.413e+00 | 0.0007 | 800.0 | 34.17% | 14136.1 | 30.77% |
| 140 | 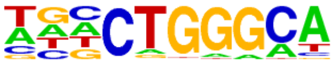 | ZNF416(Zf)/HEK293-ZNF416.GFP-ChIP-Seq(GSE58341)/Homer | 1e-3 | -8.232e+00 | 0.0008 | 455.0 | 19.44% | 7669.4 | 16.70% |
| 141 | 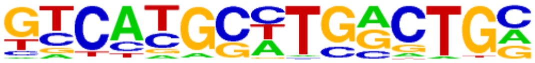 | Pax8(Paired,Homeobox)/Thyroid-Pax8-ChIP-Seq(GSE26938)/Homer | 1e-3 | -7.921e+00 | 0.0011 | 131.0 | 5.60% | 1893.3 | 4.12% |
| 142 | 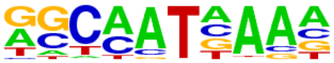 | Hoxa9(Homeobox)/ChickenMSG-Hoxa9.Flag-ChIP-Seq(GSE86088)/Homer | 1e-3 | -7.844e+00 | 0.0012 | 390.0 | 16.66% | 6508.2 | 14.17% |
| 143 | 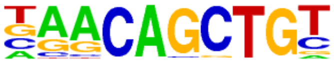 | Myf5(bHLH)/GM-Myf5-ChIP-Seq(GSE24852)/Homer | 1e-3 | -7.567e+00 | 0.0015 | 232.0 | 9.91% | 3672.6 | 7.99% |
| 144 | 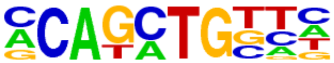 | Twist2(bHLH)/Myoblast-Twist2.Ty1-ChIP-Seq(GSE127998)/Homer | 1e-3 | -7.426e+00 | 0.0018 | 493.0 | 21.06% | 8453.5 | 18.40% |
| 145 | 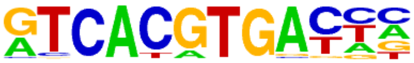 | TFE3(bHLH)/MEF-TFE3-ChIP-Seq(GSE75757)/Homer | 1e-3 | -7.392e+00 | 0.0018 | 34.0 | 1.45% | 359.8 | 0.78% |
| 146 | 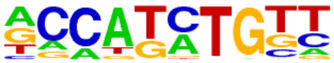 | Olig2(bHLH)/Neuron-Olig2-ChIP-Seq(GSE30882)/Homer | 1e-3 | -7.277e+00 | 0.0020 | 445.0 | 19.01% | 7576.7 | 16.49% |
| 147 | 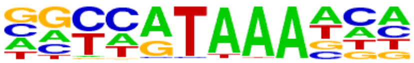 | CDX4(Homeobox)/ZebrafishEmbryos-Cdx4.Myc-ChIP-Seq(GSE48254)/Homer | 1e-3 | -7.253e+00 | 0.0021 | 81.0 | 3.46% | 1091.4 | 2.38% |
| 148 | 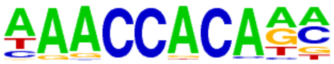 | RUNX1(Runt)/Jurkat-RUNX1-ChIP-Seq(GSE29180)/Homer | 1e-3 | -7.232e+00 | 0.0021 | 221.0 | 9.44% | 3500.4 | 7.62% |
| 149 | 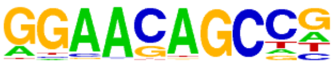 | ZNF341(Zf)/EBV-ZNF341-ChIP-Seq(GSE113194)/Homer | 1e-3 | -7.226e+00 | 0.0021 | 234.0 | 10.00% | 3732.3 | 8.12% |
| 150 | | TCF4(bHLH)/SHSY5Y-TCF4-ChIP-Seq(GSE96915)/Homer | 1e-3 | -7.136e+00 | 0.0023 | 422.0 | 18.03% | 7164.5 | 15.60% |

| | | | | | | | | | |
|---|---|---|---|---|---|---|---|---|---|
| | 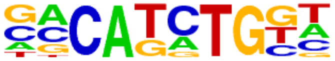 | | | | | | | | |
| 151 | 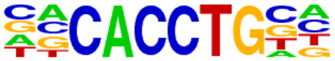 | E2A(bHLH),near_PU.1/Bcell-PU.1-ChIP-Seq(GSE21512)/Homer | 1e-3 | -7.039e+00 | 0.0025 | 553.0 | 23.62% | 9617.4 | 20.94% |
| 152 | 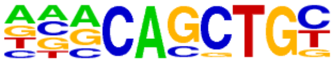 | E2A(bHLH)/proBcell-E2A-ChIP-Seq(GSE21978)/Homer | 1e-3 | -7.006e+00 | 0.0026 | 561.0 | 23.96% | 9771.4 | 21.27% |
| 153 | 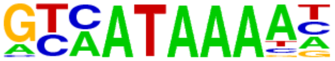 | Cdx2(Homeobox)/mES-Cdx2-ChIP-Seq(GSE14586)/Homer | 1e-3 | -6.915e+00 | 0.0028 | 56.0 | 2.39% | 703.4 | 1.53% |
| 154 | 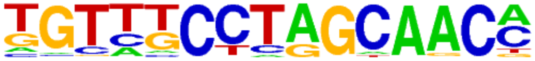 | Rfx6(HTH)/Min6b1-Rfx6.HA-ChIP-Seq(GSE62844)/Homer | 1e-2 | -6.821e+00 | 0.0030 | 306.0 | 13.07% | 5063.8 | 11.02% |
| 155 | 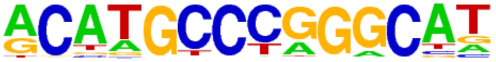 | p53(p53)/mES-cMyc-ChIP-Seq(GSE11431)/Homer | 1e-2 | -6.775e+00 | 0.0032 | 13.0 | 0.56% | 92.7 | 0.20% |
| 156 | 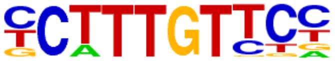 | Sox4(HMG)/proB-Sox4-ChIP-Seq(GSE50066)/Homer | 1e-2 | -6.713e+00 | 0.0033 | 151.0 | 6.45% | 2302.4 | 5.01% |
| 157 | 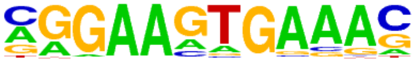 | PU.1-IRF(ETS:IRF)/Bcell-PU.1-ChIP-Seq(GSE21512)/Homer | 1e-2 | -6.697e+00 | 0.0034 | 253.0 | 10.81% | 4113.8 | 8.96% |
| 158 | 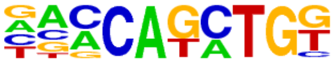 | BHLHA15(bHLH)/NIH3T3-BHLHB8.HA-ChIP-Seq(GSE119782)/Homer | 1e-2 | -6.692e+00 | 0.0034 | 402.0 | 17.17% | 6838.5 | 14.89% |
| 159 | 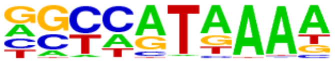 | Hoxd11(Homeobox)/ChickenMSG-Hoxd11.Flag-ChIP-Seq(GSE86088)/Homer | 1e-2 | -6.677e+00 | 0.0034 | 288.0 | 12.30% | 4748.3 | 10.34% |
| 160 | 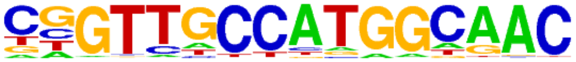 | RFX(HTH)/K562-RFX3-ChIP-Seq(SRA012198)/Homer | 1e-2 | -6.609e+00 | 0.0036 | 36.0 | 1.54% | 406.7 | 0.89% |
| 161 | 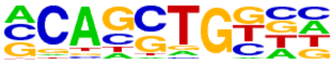 | HEB(bHLH)/mES-Heb-ChIP-Seq(GSE53233)/Homer | 1e-2 | -6.560e+00 | 0.0038 | 699.0 | 29.86% | 12436.7 | 27.07% |
| 162 | 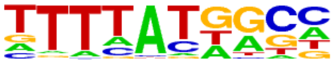 | Hoxa11(Homeobox)/ChickenMSG-Hoxa11.Flag-ChIP-Seq(GSE86088)/Homer | 1e-2 | -6.534e+00 | 0.0038 | 251.0 | 10.72% | 4090.5 | 8.90% |
| 163 | 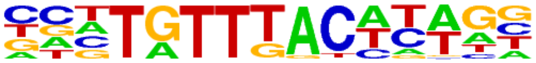 | Foxa3(Forkhead)/Liver-Foxa3-ChIP-Seq(GSE77670)/Homer | 1e-2 | -6.521e+00 | 0.0039 | 39.0 | 1.67% | 453.3 | 0.99% |
| 164 | 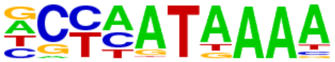 | Hoxd13(Homeobox)/ChickenMSG-Hoxd13.Flag-ChIP-Seq(GSE86088)/Homer | 1e-2 | -6.464e+00 | 0.0041 | 146.0 | 6.24% | 2230.5 | 4.86% |
| 165 | 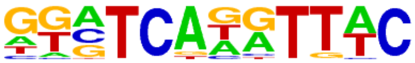 | Six1(Homeobox)/Myoblast-Six1-ChIP-Chip(GSE20150)/Homer | 1e-2 | -6.378e+00 | 0.0044 | 44.0 | 1.88% | 533.2 | 1.16% |
| 166 | 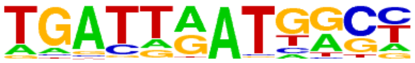 | Hoxb4(Homeobox)/ES-Hoxb4-ChIP-Seq(GSE34014)/Homer | 1e-2 | -6.111e+00 | 0.0057 | 32.0 | 1.37% | 359.5 | 0.78% |
| 167 | | HOXB13(Homeobox)/ProstateTumor-HOXB13-ChIP-Seq(GSE56288)/Homer | 1e-2 | -6.074e+00 | 0.0059 | 98.0 | 4.19% | 1426.0 | 3.10% |

| | | | | | | | | | |
|---|---|---|---|---|---|---|---|---|---|
| | 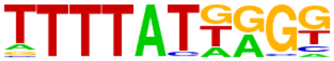 | | | | | | | | |
| 168 | 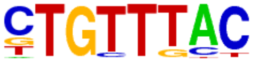 | Foxo1(Forkhead)/RAW-Foxo1-ChIP-Seq(Fan_et_al.)/Homer | 1e-2 | -6.059e+00 | 0.0060 | 332.0 | 14.18% | 5609.7 | 12.21% |
| 169 | 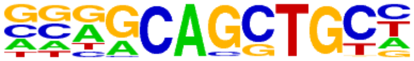 | Ascl2(bHLH)/ESC-Ascl2-ChIP-Seq(GSE97712)/Homer | 1e-2 | -6.047e+00 | 0.0060 | 441.0 | 18.84% | 7631.4 | 16.61% |
| 170 | 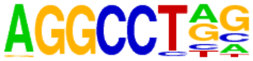 | ZFX(Zf)/mES-Zfx-ChIP-Seq(GSE11431)/Homer | 1e-2 | -6.036e+00 | 0.0060 | 542.0 | 23.15% | 9528.0 | 20.74% |
| 171 | 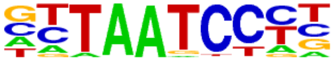 | Otx2(Homeobox)/EpiLC-Otx2-ChIP-Seq(GSE56098)/Homer | 1e-2 | -5.899e+00 | 0.0069 | 93.0 | 3.97% | 1350.6 | 2.94% |
| 172 | 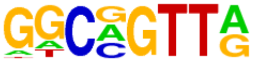 | MYB(HTH)/ERMYB-Myb-ChIPSeq(GSE22095)/Homer | 1e-2 | -5.882e+00 | 0.0069 | 371.0 | 15.85% | 6347.1 | 13.82% |
| 173 | 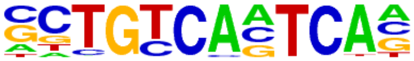 | Pknox1(Homeobox)/ES-Prep1-ChIP-Seq(GSE63282)/Homer | 1e-2 | -5.880e+00 | 0.0069 | 75.0 | 3.20% | 1049.7 | 2.29% |
| 174 | 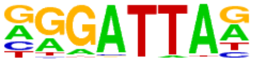 | GSC(Homeobox)/FrogEmbryos-GSC-ChIP-Seq(DRA000576)/Homer | 1e-2 | -5.856e+00 | 0.0070 | 124.0 | 5.30% | 1884.4 | 4.10% |
| 175 | 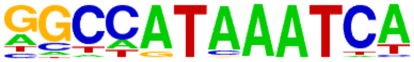 | Hoxc9(Homeobox)/Ainv15-Hoxc9-ChIP-Seq(GSE21812)/Homer | 1e-2 | -5.855e+00 | 0.0070 | 53.0 | 2.26% | 691.0 | 1.50% |
| 176 | 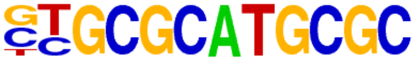 | NRF(NRF)/Promoter/Homer | 1e-2 | -5.831e+00 | 0.0071 | 106.0 | 4.53% | 1575.4 | 3.43% |
| 177 | 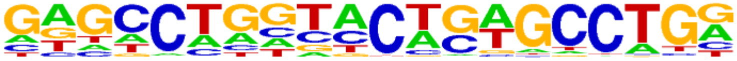 | ZNF322(Zf)/HEK293-ZNF322.GFP-ChIP-Seq(GSE58341)/Homer | 1e-2 | -5.765e+00 | 0.0076 | 123.0 | 5.25% | 1872.4 | 4.08% |
| 178 | 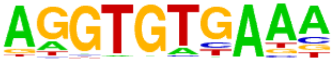 | Tbx21(T-box)/GM12878-TBX21-ChIP-Seq(Encode)/Homer | 1e-2 | -5.675e+00 | 0.0082 | 191.0 | 8.16% | 3079.3 | 6.70% |
| 179 | 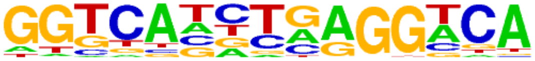 | THRa(NR)/C17.2-THRa-ChIP-Seq(GSE38347)/Homer | 1e-2 | -5.593e+00 | 0.0089 | 144.0 | 6.15% | 2250.9 | 4.90% |
| 180 | 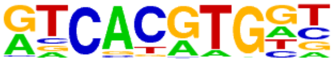 | Usf2(bHLH)/C2C12-Usf2-ChIP-Seq(GSE36030)/Homer | 1e-2 | -5.489e+00 | 0.0098 | 95.0 | 4.06% | 1405.9 | 3.06% |
| 181 | 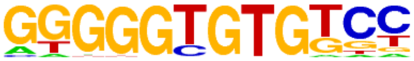 | KLF10(Zf)/HEK293-KLF10.GFP-ChIP-Seq(GSE58341)/Homer | 1e-2 | -5.485e+00 | 0.0098 | 226.0 | 9.65% | 3725.5 | 8.11% |
| 182 | 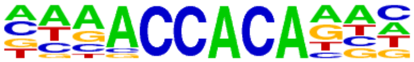 | RUNX2(Runt)/PCa-RUNX2-ChIP-Seq(GSE33889)/Homer | 1e-2 | -5.365e+00 | 0.0110 | 170.0 | 7.26% | 2727.4 | 5.94% |
| 183 | 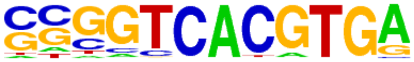 | E-box(bHLH)/Promoter/Homer | 1e-2 | -5.337e+00 | 0.0112 | 41.0 | 1.75% | 518.6 | 1.13% |
| 184 | | PBX2(Homeobox)/K562-PBX2-ChIP-Seq(Encode)/Homer | 1e-2 | -5.281e+00 | 0.0118 | 94.0 | 4.02% | 1399.3 | 3.05% |

| | | | | | | | | | |
|---|---|---|---|---|---|---|---|---|---|
| | 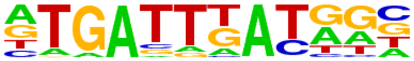 | | | | | | | | |
| 185 | 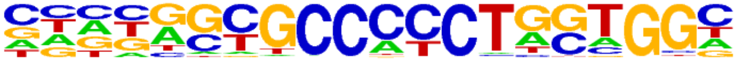 | BORIS(Zf)/K562-CTCFL-ChIP-Seq(GSE32465)/Homer | 1e-2 | -5.139e+00 | 0.0136 | 103.0 | 4.40% | 1562.5 | 3.40% |
| 186 | 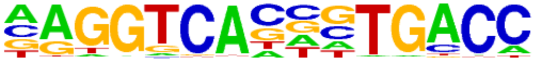 | ERE(NR),IR3/MCF7-ERa-ChIP-Seq(Unpublished)/Homer | 1e-2 | -5.039e+00 | 0.0149 | 87.0 | 3.72% | 1292.9 | 2.81% |
| 187 | 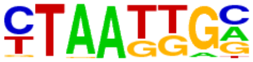 | Isl1(Homeobox)/Neuron-Isl1-ChIP-Seq(GSE31456)/Homer | 1e-2 | -4.964e+00 | 0.0160 | 245.0 | 10.47% | 4118.0 | 8.96% |
| 188 | 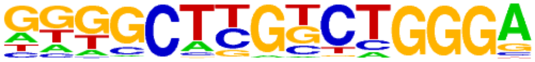 | Zfp809(Zf)/ES-Zfp809-ChIP-Seq(GSE70799)/Homer | 1e-2 | -4.804e+00 | 0.0187 | 117.0 | 5.00% | 1827.6 | 3.98% |
| 189 | 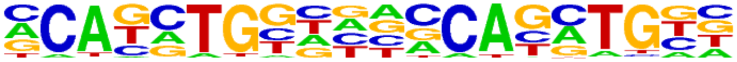 | Twist(bHLH)/HMLE-TWIST1-ChIP-Seq(Chang_et_al)/Homer | 1e-2 | -4.734e+00 | 0.0199 | 46.0 | 1.96% | 619.8 | 1.35% |
| 190 | 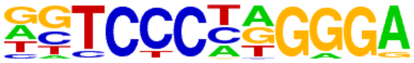 | EBF(EBF)/proBcell-EBF-ChIP-Seq(GSE21978)/Homer | 1e-2 | -4.720e+00 | 0.0201 | 85.0 | 3.63% | 1275.4 | 2.78% |
| 191 | 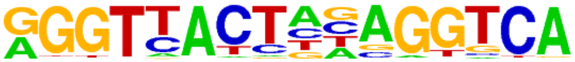 | LXRE(NR),DR4/RAW-LXRb.biotin-ChIP-Seq(GSE21512)/Homer | 1e-2 | -4.701e+00 | 0.0204 | 16.0 | 0.68% | 159.8 | 0.35% |
| 192 | 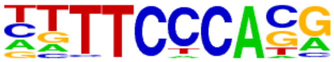 | Rbpj1(?)/Panc1-Rbpj1-ChIP-Seq(GSE47459)/Homer | 1e-2 | -4.680e+00 | 0.0207 | 338.0 | 14.44% | 5866.6 | 12.77% |
| 193 | 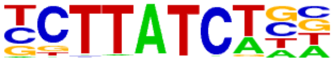 | Gata6(Zf)/HUG1N-GATA6-ChIP-Seq(GSE51936)/Homer | 1e-2 | -4.665e+00 | 0.0209 | 92.0 | 3.93% | 1399.7 | 3.05% |
| 194 | 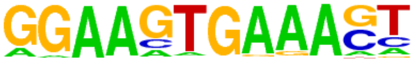 | PU.1:IRF8(ETS:IRF)/pDC-Irf8-ChIP-Seq(GSE66899)/Homer | 1e-2 | -4.629e+00 | 0.0215 | 36.0 | 1.54% | 462.6 | 1.01% |
