## Supplemental Table S10 for "PRMT1 Inhibition Selectively Targets BNC1-Dependent Proliferation, but not Migration in Squamous Cell Carcinoma"

### Homer Known Motif Enrichment Results (QC/homer/09_0937_SCC-13_Fra1_hg38)

[Homer *de novo* Motif Results]
[Gene Ontology Enrichment Results]
[Known Motif Enrichment Results (txt file)]
Total Target Sequences = 1959, Total Background Sequences = 43686

| Rank | Motif | Name | P-value | log P-pvalue | q-value (Benjamini) | # Target Sequences with Motif | % of Targets Sequences with Motif | # Background Sequences with Motif | % of Background Sequences with Motif |
|---|---|---|---|---|---|---|---|---|---|
| 1 | 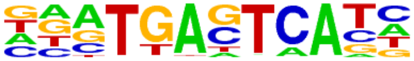 | Fra1(bZIP)/BT549-Fra1-ChIP-Seq(GSE46166)/Homer | 1e-1632 | -3.759e+03 | 0.0000 | 1603.0 | 81.83% | 2380.9 | 5.45% |
| 2 | 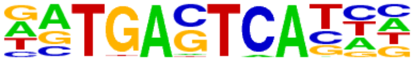 | Atf3(bZIP)/GBM-ATF3-ChIP-Seq(GSE33912)/Homer | 1e-1603 | -3.693e+03 | 0.0000 | 1651.0 | 84.28% | 2823.1 | 6.47% |
| 3 | 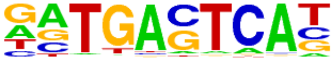 | JunB(bZIP)/DendriticCells-Junb-ChIP-Seq(GSE36099)/Homer | 1e-1585 | -3.651e+03 | 0.0000 | 1586.0 | 80.96% | 2432.5 | 5.57% |
| 4 | 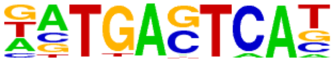 | BATF(bZIP)/Th17-BATF-ChIP-Seq(GSE39756)/Homer | 1e-1581 | -3.641e+03 | 0.0000 | 1630.0 | 83.21% | 2757.3 | 6.32% |
| 5 | 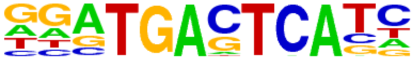 | Fra2(bZIP)/Striatum-Fra2-ChIP-Seq(GSE43429)/Homer | 1e-1566 | -3.608e+03 | 0.0000 | 1529.0 | 78.05% | 2136.8 | 4.89% |
| 6 | 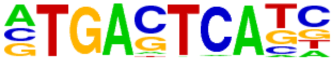 | AP-1(bZIP)/ThioMac-PU.1-ChIP-Seq(GSE21512)/Homer | 1e-1502 | -3.461e+03 | 0.0000 | 1660.0 | 84.74% | 3333.1 | 7.64% |
| 7 | 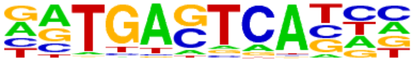 | Fosl2(bZIP)/3T3L1-Fosl2-ChIP-Seq(GSE56872)/Homer | 1e-1497 | -3.448e+03 | 0.0000 | 1381.0 | 70.50% | 1548.6 | 3.55% |
| 8 | 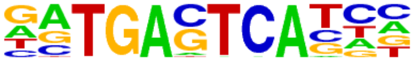 | Jun-AP1(bZIP)/K562-cJun-ChIP-Seq(GSE31477)/Homer | 1e-1364 | -3.143e+03 | 0.0000 | 1210.0 | 61.77% | 1129.9 | 2.59% |
| 9 | 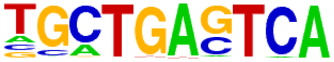 | Bach2(bZIP)/OCILy7-Bach2-ChIP-Seq(GSE44420)/Homer | 1e-688 | -1.585e+03 | 0.0000 | 761.0 | 38.85% | 1016.8 | 2.33% |
| 10 | 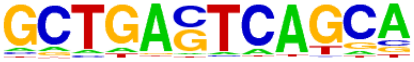 | MafK(bZIP)/C2C12-MafK-ChIP-Seq(GSE36030)/Homer | 1e-246 | -5.666e+02 | 0.0000 | 409.0 | 20.88% | 1038.7 | 2.38% |
| 11 | 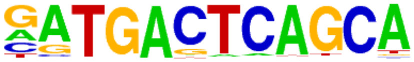 | NF-E2(bZIP)/K562-NFE2-ChIP-Seq(GSE31477)/Homer | 1e-217 | -5.010e+02 | 0.0000 | 246.0 | 12.56% | 299.1 | 0.69% |
| 12 | 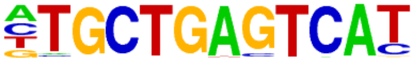 | Nrf2(bZIP)/Lymphoblast-Nrf2-ChIP-Seq(GSE37589)/Homer | 1e-184 | -4.237e+02 | 0.0000 | 203.0 | 10.36% | 232.6 | 0.53% |
| 13 | 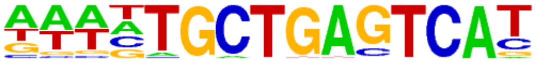 | Bach1(bZIP)/K562-Bach1-ChIP-Seq(GSE31477)/Homer | 1e-145 | -3.352e+02 | 0.0000 | 185.0 | 9.44% | 282.5 | 0.65% |
| 14 | | NFE2L2(bZIP)/HepG2-NFE2L2-ChIP-Seq(Encode)/Homer | 1e-128 | -2.957e+02 | 0.0000 | 150.0 | 7.66% | 192.0 | 0.44% |

| | | | | | | | | | |
|---|---|---|---|---|---|---|---|---|---|
| | 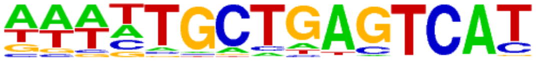 | | | | | | | | |
| 15 | 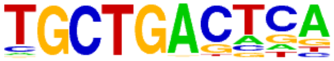 | MafA(bZIP)/Islet-MafA-ChIP-Seq(GSE30298)/Homer | 1e-111 | -2.557e+02 | 0.0000 | 513.0 | 26.19% | 3856.6 | 8.83% |
| 16 | 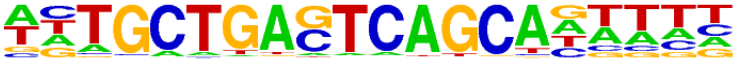 | MafB(bZIP)/BMM-Mafb-ChIP-Seq(GSE75722)/Homer | 1e-83 | -1.917e+02 | 0.0000 | 290.0 | 14.80% | 1684.7 | 3.86% |
| 17 | 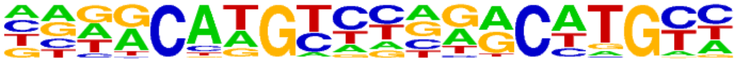 | p63(p53)/Keratinocyte-p63-ChIP-Seq(GSE17611)/Homer | 1e-38 | -8.755e+01 | 0.0000 | 177.0 | 9.04% | 1270.2 | 2.91% |
| 18 | 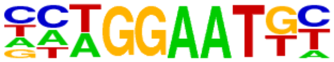 | TEAD(TEA)/Fibroblast-PU.1-ChIP-Seq(Unpublished)/Homer | 1e-31 | -7.352e+01 | 0.0000 | 242.0 | 12.35% | 2337.3 | 5.35% |
| 19 | 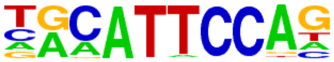 | TEAD3(TEA)/HepG2-TEAD3-ChIP-Seq(Encode)/Homer | 1e-31 | -7.326e+01 | 0.0000 | 359.0 | 18.33% | 4181.1 | 9.58% |
| 20 | 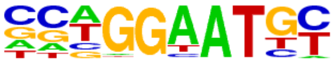 | TEAD4(TEA)/Tropoblast-Tead4-ChIP-Seq(GSE37350)/Homer | 1e-31 | -7.200e+01 | 0.0000 | 301.0 | 15.36% | 3268.0 | 7.49% |
| 21 | 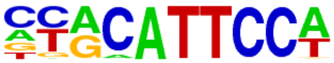 | TEAD1(TEAD)/HepG2-TEAD1-ChIP-Seq(Encode)/Homer | 1e-31 | -7.172e+01 | 0.0000 | 317.0 | 16.18% | 3528.8 | 8.08% |
| 22 | 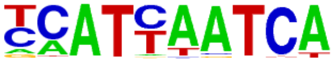 | Pdx1(Homeobox)/Islet-Pdx1-ChIP-Seq(SRA008281)/Homer | 1e-30 | -7.082e+01 | 0.0000 | 314.0 | 16.03% | 3498.2 | 8.01% |
| 23 | 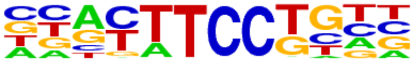 | Etv2(ETS)/ES-ER71-ChIP-Seq(GSE59402)/Homer | 1e-29 | -6.826e+01 | 0.0000 | 314.0 | 16.03% | 3551.1 | 8.13% |
| 24 | 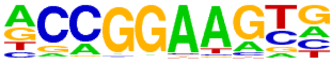 | ETV4(ETS)/HepG2-ETV4-ChIP-Seq(ENCODE)/Homer | 1e-27 | -6.424e+01 | 0.0000 | 361.0 | 18.43% | 4423.3 | 10.13% |
| 25 | 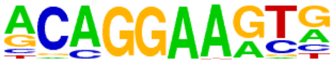 | ERG(ETS)/VCaP-ERG-ChIP-Seq(GSE14097)/Homer | 1e-26 | -6.070e+01 | 0.0000 | 472.0 | 24.09% | 6481.0 | 14.85% |
| 26 | 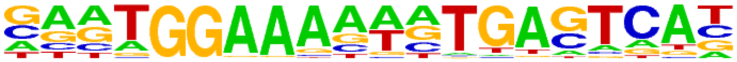 | NFAT:AP1(RHD,bZIP)/Jurkat-NFATC1-ChIP-Seq(Jolma_et_al.)/Homer | 1e-26 | -6.007e+01 | 0.0000 | 92.0 | 4.70% | 532.1 | 1.22% |
| 27 | 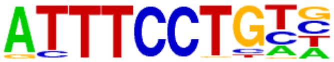 | EWS:ERG-fusion(ETS)/CADO_ES1-EWS:ERG-ChIP-Seq(SRA014231)/Homer | 1e-25 | -5.913e+01 | 0.0000 | 241.0 | 12.30% | 2574.1 | 5.90% |
| 28 | 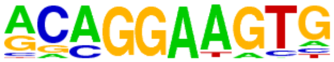 | ETS1(ETS)/Jurkat-ETS1-ChIP-Seq(GSE17954)/Homer | 1e-25 | -5.906e+01 | 0.0000 | 341.0 | 17.41% | 4210.7 | 9.65% |
| 29 | 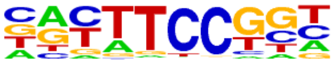 | Fli1(ETS)/CD8-FLI-ChIP-Seq(GSE20898)/Homer | 1e-24 | -5.730e+01 | 0.0000 | 340.0 | 17.36% | 4237.1 | 9.71% |
| 30 | 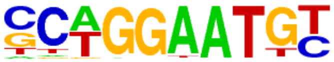 | TEAD2(TEA)/Py2T-Tead2-ChIP-Seq(GSE55709)/Homer | 1e-23 | -5.477e+01 | 0.0000 | 191.0 | 9.75% | 1893.0 | 4.34% |
| 31 | | p53(p53)/Saos-p53-ChIP-Seq(GSE15780)/Homer | 1e-23 | -5.372e+01 | 0.0000 | 67.0 | 3.42% | 322.0 | 0.74% |

| | | | | | | | | | |
|---|---|---|---|---|---|---|---|---|---|
| | 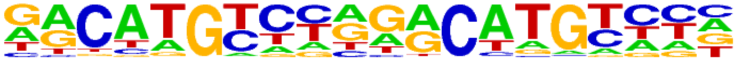 | | | | | | | | |
| 32 | 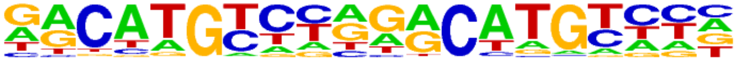 | p53(p53)/Saos-p53-ChIP-Seq/Homer | 1e-23 | -5.372e+01 | 0.0000 | 67.0 | 3.42% | 322.0 | 0.74% |
| 33 | 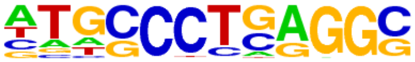 | AP-2alpha(AP2)/Hela-AP2alpha-ChIP-Seq(GSE31477)/Homer | 1e-23 | -5.334e+01 | 0.0000 | 312.0 | 15.93% | 3860.8 | 8.84% |
| 34 | 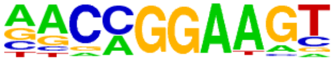 | ETV1(ETS)/GIST48-ETV1-ChIP-Seq(GSE22441)/Homer | 1e-22 | -5.244e+01 | 0.0000 | 400.0 | 20.42% | 5425.8 | 12.43% |
| 35 | 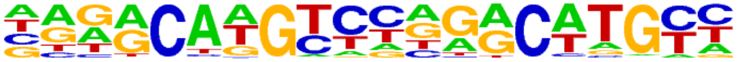 | p73(p53)/Trachea-p73-ChIP-Seq(PRJNA310161)/Homer | 1e-20 | -4.809e+01 | 0.0000 | 44.0 | 2.25% | 151.0 | 0.35% |
| 36 | 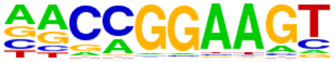 | GABPA(ETS)/Jurkat-GABPa-ChIP-Seq(GSE17954)/Homer | 1e-20 | -4.706e+01 | 0.0000 | 270.0 | 13.78% | 3308.9 | 7.58% |
| 37 | 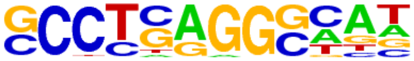 | AP-2gamma(AP2)/MCF7-TFAP2C-ChIP-Seq(GSE21234)/Homer | 1e-20 | -4.652e+01 | 0.0000 | 361.0 | 18.43% | 4906.0 | 11.24% |
| 38 | 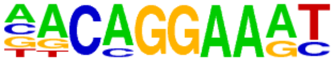 | EWS:FLI1-fusion(ETS)/SK_N_MC-EWS:FLI1-ChIP-Seq(SRA014231)/Homer | 1e-19 | -4.445e+01 | 0.0000 | 190.0 | 9.70% | 2067.6 | 4.74% |
| 39 | 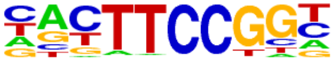 | Elk1(ETS)/Hela-Elk1-ChIP-Seq(GSE31477)/Homer | 1e-19 | -4.411e+01 | 0.0000 | 176.0 | 8.98% | 1857.0 | 4.25% |
| 40 | 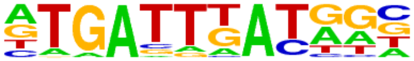 | PBX2(Homeobox)/K562-PBX2-ChIP-Seq(Encode)/Homer | 1e-16 | -3.697e+01 | 0.0000 | 232.0 | 11.84% | 2927.3 | 6.71% |
| 41 | 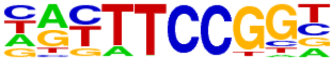 | Elk4(ETS)/Hela-Elk4-ChIP-Seq(GSE31477)/Homer | 1e-12 | -2.971e+01 | 0.0000 | 155.0 | 7.91% | 1830.7 | 4.19% |
| 42 | 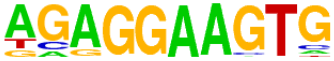 | PU.1(ETS)/ThioMac-PU.1-ChIP-Seq(GSE21512)/Homer | 1e-11 | -2.759e+01 | 0.0000 | 157.0 | 8.01% | 1915.3 | 4.39% |
| 43 | 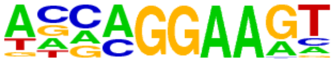 | EHF(ETS)/LoVo-EHF-ChIP-Seq(GSE49402)/Homer | 1e-11 | -2.715e+01 | 0.0000 | 317.0 | 16.18% | 4778.3 | 10.95% |
| 44 | 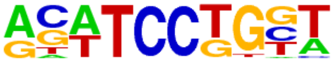 | SPDEF(ETS)/VCaP-SPDEF-ChIP-Seq(SRA014231)/Homer | 1e-11 | -2.564e+01 | 0.0000 | 269.0 | 13.73% | 3946.6 | 9.04% |
| 45 | 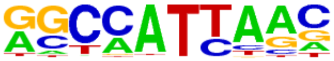 | Nanog(Homeobox)/mES-Nanog-ChIP-Seq(GSE11724)/Homer | 1e-10 | -2.529e+01 | 0.0000 | 918.0 | 46.86% | 17188.6 | 39.38% |
| 46 | 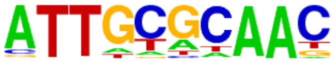 | CEBP(bZIP)/ThioMac-CEBPb-ChIP-Seq(GSE21512)/Homer | 1e-10 | -2.471e+01 | 0.0000 | 145.0 | 7.40% | 1790.9 | 4.10% |
| 47 | 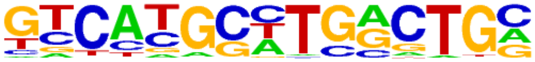 | Pax8(Paired,Homeobox)/Thyroid-Pax8-ChIP-Seq(GSE26938)/Homer | 1e-10 | -2.438e+01 | 0.0000 | 123.0 | 6.28% | 1438.6 | 3.30% |
| 48 | | Elf4(ETS)/BMDM-Elf4-ChIP-Seq(GSE88699)/Homer | 1e-10 | -2.391e+01 | 0.0000 | 259.0 | 13.22% | 3829.6 | 8.77% |

| | | | | | | | | | |
|---|---|---|---|---|---|---|---|---|---|
| | 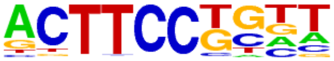 | | | | | | | | |
| 49 | 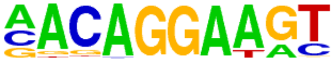 | Ets1-distal(ETS)/CD4+-PolII-ChIP-Seq(Barski_et_al.)/Homer | 1e-10 | -2.363e+01 | 0.0000 | 113.0 | 5.77% | 1295.2 | 2.97% |
| 50 | 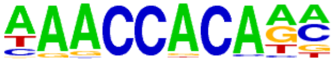 | RUNX1(Runt)/Jurkat-RUNX1-ChIP-Seq(GSE29180)/Homer | 1e-9 | -2.300e+01 | 0.0000 | 270.0 | 13.78% | 4070.2 | 9.32% |
| 51 | 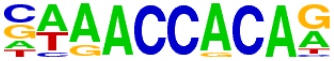 | RUNX(Runt)/HPC7-Runx1-ChIP-Seq(GSE22178)/Homer | 1e-9 | -2.192e+01 | 0.0000 | 207.0 | 10.57% | 2955.5 | 6.77% |
| 52 | 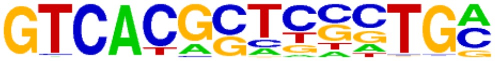 | PAX5(Paired,Homeobox),condensed/GM12878-PAX5-ChIP-Seq(GSE32465)/Homer | 1e-9 | -2.189e+01 | 0.0000 | 58.0 | 2.96% | 506.3 | 1.16% |
| 53 | 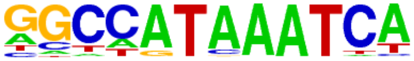 | Hoxc9(Homeobox)/Ainv15-Hoxc9-ChIP-Seq(GSE21812)/Homer | 1e-9 | -2.132e+01 | 0.0000 | 115.0 | 5.87% | 1382.3 | 3.17% |
| 54 | 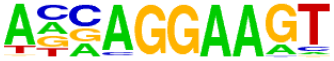 | ELF3(ETS)/PDAC-ELF3-ChIP-Seq(GSE64557)/Homer | 1e-8 | -1.971e+01 | 0.0000 | 189.0 | 9.65% | 2711.8 | 6.21% |
| 55 | 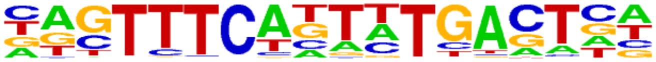 | bZIP:IRF(bZIP,IRF)/Th17-BatF-ChIP-Seq(GSE39756)/Homer | 1e-8 | -1.910e+01 | 0.0000 | 99.0 | 5.05% | 1176.6 | 2.70% |
| 56 | 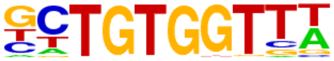 | RUNX-AML(Runt)/CD4+-PolII-ChIP-Seq(Barski_et_al.)/Homer | 1e-7 | -1.815e+01 | 0.0000 | 205.0 | 10.46% | 3064.4 | 7.02% |
| 57 | 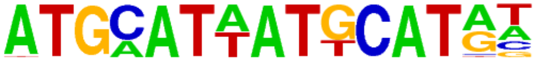 | OCT:OCT-short(POU,Homeobox)/NPC-OCT6-ChIP-Seq(GSE43916)/Homer | 1e-7 | -1.804e+01 | 0.0000 | 159.0 | 8.12% | 2233.6 | 5.12% |
| 58 | 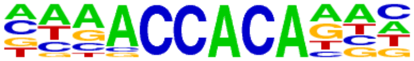 | RUNX2(Runt)/PCa-RUNX2-ChIP-Seq(GSE33889)/Homer | 1e-7 | -1.791e+01 | 0.0000 | 219.0 | 11.18% | 3334.6 | 7.64% |
| 59 | 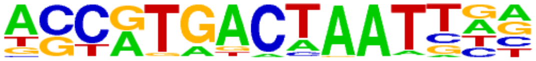 | PAX3:FKHR-fusion(Paired,Homeobox)/Rh4-PAX3:FKHR-ChIP-Seq(GSE19063)/Homer | 1e-7 | -1.783e+01 | 0.0000 | 61.0 | 3.11% | 612.3 | 1.40% |
| 60 | 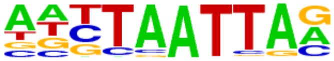 | Lhx3(Homeobox)/Neuron-Lhx3-ChIP-Seq(GSE31456)/Homer | 1e-7 | -1.782e+01 | 0.0000 | 308.0 | 15.72% | 5037.2 | 11.54% |
| 61 | 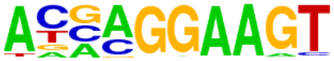 | ELF5(ETS)/T47D-ELF5-ChIP-Seq(GSE30407)/Homer | 1e-7 | -1.771e+01 | 0.0000 | 177.0 | 9.04% | 2569.5 | 5.89% |
| 62 | 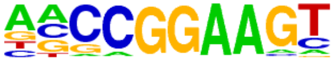 | ELF1(ETS)/Jurkat-ELF1-ChIP-Seq(SRA014231)/Homer | 1e-7 | -1.740e+01 | 0.0000 | 129.0 | 6.58% | 1727.7 | 3.96% |
| 63 | 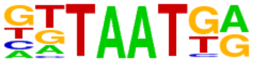 | Nkx6.1(Homeobox)/Islet-Nkx6.1-ChIP-Seq(GSE40975)/Homer | 1e-7 | -1.676e+01 | 0.0000 | 487.0 | 24.86% | 8689.8 | 19.91% |
| 64 | 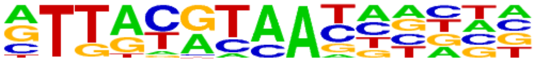 | NFIL3(bZIP)/HepG2-NFIL3-ChIP-Seq(Encode)/Homer | 1e-7 | -1.675e+01 | 0.0000 | 127.0 | 6.48% | 1713.1 | 3.92% |
| 65 | | Hoxb4(Homeobox)/ES-Hoxb4-ChIP-Seq(GSE34014)/Homer | 1e-7 | -1.652e+01 | 0.0000 | 69.0 | 3.52% | 758.3 | 1.74% |

| | | | | | | | | | |
|---|---|---|---|---|---|---|---|---|---|
| | 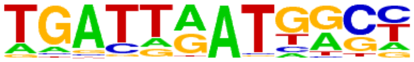 | | | | | | | | |
| 66 | 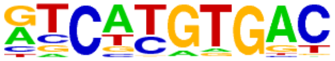 | MITF(bHLH)/MastCells-MITF-ChIP-Seq(GSE48085)/Homer | 1e-6 | -1.597e+01 | 0.0000 | 256.0 | 13.07% | 4127.3 | 9.45% |
| 67 | 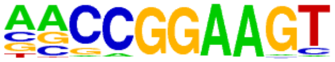 | ETS(ETS)/Promoter/Homer | 1e-6 | -1.564e+01 | 0.0000 | 86.0 | 4.39% | 1050.1 | 2.41% |
| 68 | 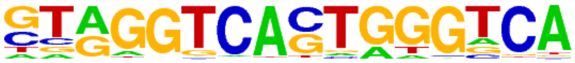 | Reverb(NR),DR2/RAW-Reverba.biotin-ChIP-Seq(GSE45914)/Homer | 1e-6 | -1.532e+01 | 0.0000 | 63.0 | 3.22% | 690.6 | 1.58% |
| 69 | 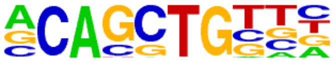 | Ptf1a(bHLH)/Panc1-Ptf1a-ChIP-Seq(GSE47459)/Homer | 1e-6 | -1.509e+01 | 0.0000 | 727.0 | 37.11% | 13861.6 | 31.75% |
| 70 | 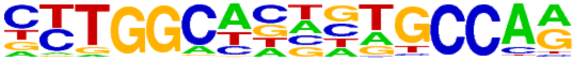 | NF1(CTF)/LNCAP-NF1-ChIP-Seq(Unpublished)/Homer | 1e-6 | -1.501e+01 | 0.0000 | 121.0 | 6.18% | 1665.4 | 3.82% |
| 71 | 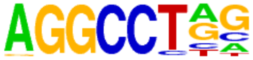 | ZFX(Zf)/mES-Zfx-ChIP-Seq(GSE11431)/Homer | 1e-6 | -1.490e+01 | 0.0000 | 365.0 | 18.63% | 6337.8 | 14.52% |
| 72 | 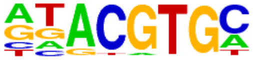 | HIF-1b(HLH)/T47D-HIF1b-ChIP-Seq(GSE59937)/Homer | 1e-6 | -1.457e+01 | 0.0000 | 261.0 | 13.32% | 4298.6 | 9.85% |
| 73 | 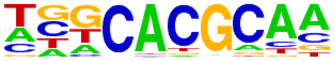 | Arnt:Ahr(bHLH)/MCF7-Arnt-ChIP-Seq(Lo_et_al.)/Homer | 1e-6 | -1.456e+01 | 0.0000 | 161.0 | 8.22% | 2403.0 | 5.50% |
| 74 | 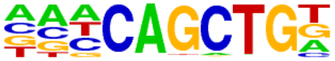 | Ap4(bHLH)/AML-Tfap4-ChIP-Seq(GSE45738)/Homer | 1e-6 | -1.452e+01 | 0.0000 | 344.0 | 17.56% | 5940.3 | 13.61% |
| 75 | 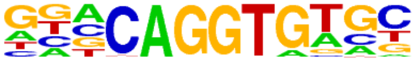 | ZEB2(Zf)/SNU398-ZEB2-ChIP-Seq(GSE103048)/Homer | 1e-6 | -1.430e+01 | 0.0000 | 326.0 | 16.64% | 5595.3 | 12.82% |
| 76 | 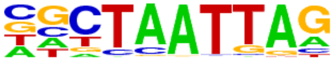 | LXH9(Homeobox)/Hct116-LXH9.V5-ChIP-Seq(GSE116822)/Homer | 1e-6 | -1.394e+01 | 0.0000 | 276.0 | 14.09% | 4627.9 | 10.60% |
| 77 | 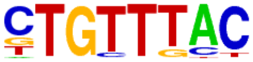 | Foxo1(Forkhead)/RAW-Foxo1-ChIP-Seq(Fan_et_al.)/Homer | 1e-5 | -1.344e+01 | 0.0000 | 371.0 | 18.94% | 6556.8 | 15.02% |
| 78 | 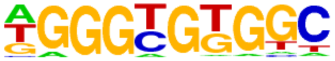 | KLF5(Zf)/LoVo-KLF5-ChIP-Seq(GSE49402)/Homer | 1e-5 | -1.325e+01 | 0.0000 | 297.0 | 15.16% | 5084.3 | 11.65% |
| 79 | 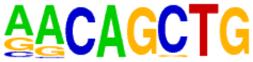 | MyoG(bHLH)/C2C12-MyoG-ChIP-Seq(GSE36024)/Homer | 1e-5 | -1.317e+01 | 0.0000 | 306.0 | 15.62% | 5268.2 | 12.07% |
| 80 | 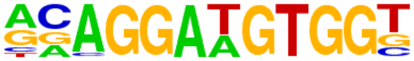 | ETS:RUNX(ETS,Runt)/Jurkat-RUNX1-ChIP-Seq(GSE17954)/Homer | 1e-5 | -1.285e+01 | 0.0000 | 40.0 | 2.04% | 391.5 | 0.90% |
| 81 | 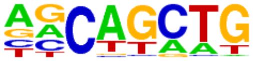 | SCL(bHLH)/HPC7-Scl-ChIP-Seq(GSE13511)/Homer | 1e-5 | -1.250e+01 | 0.0000 | 1069.0 | 54.57% | 21602.4 | 49.49% |
| 82 | | ZNF711(Zf)/SHSY5Y-ZNF711-ChIP-Seq(GSE20673)/Homer | 1e-5 | -1.223e+01 | 0.0000 | 457.0 | 23.33% | 8412.6 | 19.27% |

| | | | | | | | | | |
|---|---|---|---|---|---|---|---|---|---|
| | 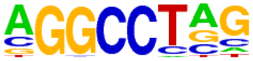 | | | | | | | | |
| 83 | 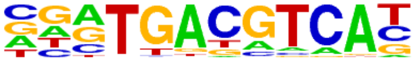 | Atf7(bZIP)/3T3L1-Atf7-ChIP-Seq(GSE56872)/Homer | 1e-5 | -1.218e+01 | 0.0000 | 139.0 | 7.10% | 2097.3 | 4.80% |
| 84 | 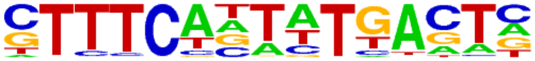 | IRF:BATF(IRF:bZIP)/pDC-Irf8-ChIP-Seq(GSE66899)/Homer | 1e-5 | -1.154e+01 | 0.0000 | 33.0 | 1.68% | 312.6 | 0.72% |
| 85 | 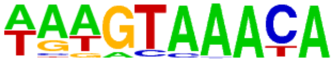 | FOXA1(Forkhead)/LNCAP-FOXA1-ChIP-Seq(GSE27824)/Homer | 1e-4 | -1.142e+01 | 0.0001 | 188.0 | 9.60% | 3058.7 | 7.01% |
| 86 | 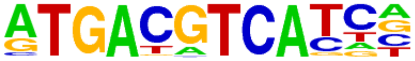 | JunD(bZIP)/K562-JunD-ChIP-Seq/Homer | 1e-4 | -1.139e+01 | 0.0001 | 38.0 | 1.94% | 387.3 | 0.89% |
| 87 | 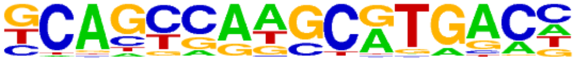 | PAX5(Paired,Homeobox)/GM12878-PAX5-ChIP-Seq(GSE32465)/Homer | 1e-4 | -1.073e+01 | 0.0001 | 108.0 | 5.51% | 1589.3 | 3.64% |
| 88 | 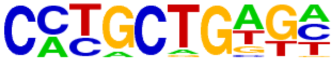 | Zic(Zf)/Cerebellum-ZIC1.2-ChIP-Seq(GSE60731)/Homer | 1e-4 | -1.025e+01 | 0.0002 | 233.0 | 11.89% | 4006.5 | 9.18% |
| 89 | 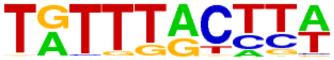 | FOXM1(Forkhead)/MCF7-FOXM1-ChIP-Seq(GSE72977)/Homer | 1e-4 | -1.025e+01 | 0.0002 | 161.0 | 8.22% | 2602.2 | 5.96% |
| 90 | 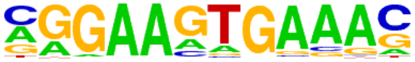 | PU.1-IRF(ETS:IRF)/Bcell-PU.1-ChIP-Seq(GSE21512)/Homer | 1e-4 | -1.010e+01 | 0.0002 | 251.0 | 12.81% | 4374.8 | 10.02% |
| 91 | 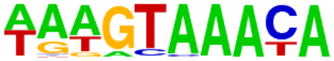 | FOXA1(Forkhead)/MCF7-FOXA1-ChIP-Seq(GSE26831)/Homer | 1e-4 | -9.984e+00 | 0.0002 | 147.0 | 7.50% | 2349.9 | 5.38% |
| 92 | 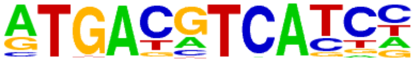 | c-Jun-CRE(bZIP)/K562-cJun-ChIP-Seq(GSE31477)/Homer | 1e-4 | -9.807e+00 | 0.0003 | 95.0 | 4.85% | 1390.6 | 3.19% |
| 93 | 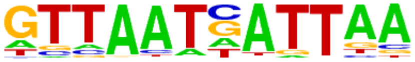 | HNF1b(Homeobox)/PDAC-HNF1B-ChIP-Seq(GSE64557)/Homer | 1e-4 | -9.782e+00 | 0.0003 | 29.0 | 1.48% | 283.6 | 0.65% |
| 94 | 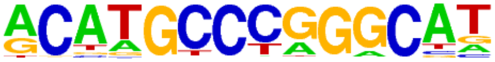 | p53(p53)/mES-cMyc-ChIP-Seq(GSE11431)/Homer | 1e-4 | -9.731e+00 | 0.0003 | 10.0 | 0.51% | 46.4 | 0.11% |
| 95 | 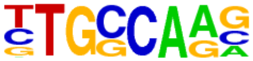 | NF1-halfsite(CTF)/LNCaP-NF1-ChIP-Seq(Unpublished)/Homer | 1e-4 | -9.562e+00 | 0.0003 | 430.0 | 21.95% | 8082.1 | 18.51% |
| 96 | 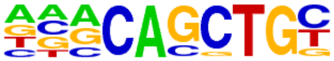 | E2A(bHLH)/proBcell-E2A-ChIP-Seq(GSE21978)/Homer | 1e-4 | -9.530e+00 | 0.0003 | 416.0 | 21.24% | 7793.2 | 17.85% |
| 97 | 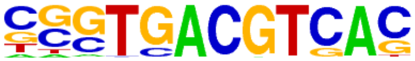 | CRE(bZIP)/Promoter/Homer | 1e-4 | -9.520e+00 | 0.0003 | 68.0 | 3.47% | 921.5 | 2.11% |
| 98 | 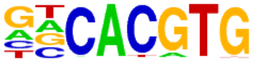 | BMAL1(bHLH)/Liver-Bmal1-ChIP-Seq(GSE39860)/Homer | 1e-4 | -9.487e+00 | 0.0003 | 459.0 | 23.43% | 8696.2 | 19.92% |
| 99 | | Tbet(T-box)/CD8-Tbet-ChIP-Seq(GSE33802)/Homer | 1e-4 | -9.447e+00 | 0.0003 | 262.0 | 13.37% | 4641.6 | 10.63% |

| | | | | | | | | | |
|---|---|---|---|---|---|---|---|---|---|
| | 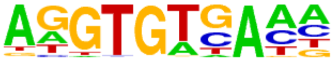 | | | | | | | | |
| 100 | 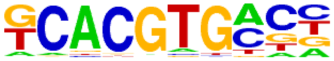 | bHLHE41(bHLH)/proB-Bhlhe41-ChIP-Seq(GSE93764)/Homer | 1e-4 | -9.218e+00 | 0.0004 | 268.0 | 13.68% | 4780.0 | 10.95% |
| 101 | 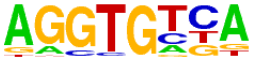 | Tbx5(T-box)/HL1-Tbx5.biotin-ChIP-Seq(GSE21529)/Homer | 1e-3 | -9.169e+00 | 0.0004 | 777.0 | 39.66% | 15539.3 | 35.60% |
| 102 | 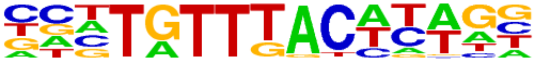 | Foxa3(Forkhead)/Liver-Foxa3-ChIP-Seq(GSE77670)/Homer | 1e-3 | -9.087e+00 | 0.0005 | 62.0 | 3.16% | 831.9 | 1.91% |
| 103 | 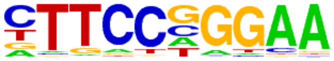 | Stat3(Stat)/mES-Stat3-ChIP-Seq(GSE11431)/Homer | 1e-3 | -8.985e+00 | 0.0005 | 109.0 | 5.56% | 1682.6 | 3.85% |
| 104 | 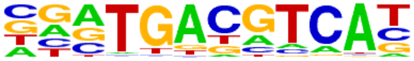 | Atf2(bZIP)/3T3L1-Atf2-ChIP-Seq(GSE56872)/Homer | 1e-3 | -8.853e+00 | 0.0006 | 100.0 | 5.10% | 1521.3 | 3.48% |
| 105 | 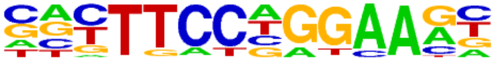 | Stat3+il21(Stat)/CD4-Stat3-ChIP-Seq(GSE19198)/Homer | 1e-3 | -8.239e+00 | 0.0011 | 142.0 | 7.25% | 2347.2 | 5.38% |
| 106 | 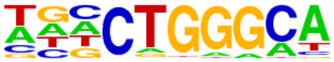 | ZNF416(Zf)/HEK293-ZNF416.GFP-ChIP-Seq(GSE58341)/Homer | 1e-3 | -8.220e+00 | 0.0011 | 340.0 | 17.36% | 6335.8 | 14.51% |
| 107 | 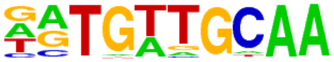 | CEBP:AP1(bZIP)/ThioMac-CEBPb-ChIP-Seq(GSE21512)/Homer | 1e-3 | -8.177e+00 | 0.0011 | 153.0 | 7.81% | 2564.5 | 5.87% |
| 108 | 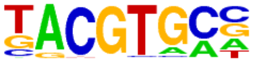 | HIF-1a(bHLH)/MCF7-HIF1a-ChIP-Seq(GSE28352)/Homer | 1e-3 | -8.068e+00 | 0.0012 | 67.0 | 3.42% | 953.6 | 2.18% |
| 109 | 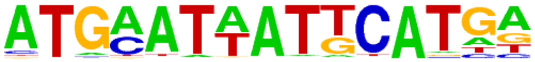 | OCT:OCT(POU,Homeobox,IR1)/NPC-Brn2-ChIP-Seq(GSE35496)/Homer | 1e-3 | -8.023e+00 | 0.0013 | 7.0 | 0.36% | 28.7 | 0.07% |
| 110 | 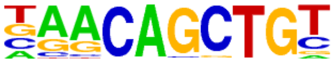 | Myf5(bHLH)/GM-Myf5-ChIP-Seq(GSE24852)/Homer | 1e-3 | -7.972e+00 | 0.0013 | 193.0 | 9.85% | 3365.6 | 7.71% |
| 111 | 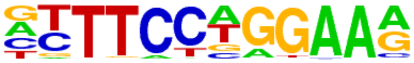 | STAT4(Stat)/CD4-Stat4-ChIP-Seq(GSE22104)/Homer | 1e-3 | -7.938e+00 | 0.0014 | 173.0 | 8.83% | 2971.0 | 6.81% |
| 112 | 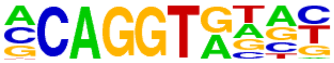 | ZEB1(Zf)/PDAC-ZEB1-ChIP-Seq(GSE64557)/Homer | 1e-3 | -7.775e+00 | 0.0016 | 481.0 | 24.55% | 9336.3 | 21.39% |
| 113 | 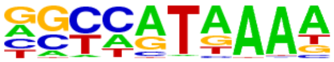 | Hoxd11(Homeobox)/ChickenMSG-Hoxd11.Flag-ChIP-Seq(GSE86088)/Homer | 1e-3 | -7.772e+00 | 0.0016 | 406.0 | 20.72% | 7754.1 | 17.76% |
| 114 | 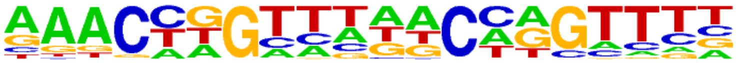 | GRHL2(CP2)/HBE-GRHL2-ChIP-Seq(GSE46194)/Homer | 1e-3 | -7.768e+00 | 0.0016 | 72.0 | 3.68% | 1054.8 | 2.42% |
| 115 | 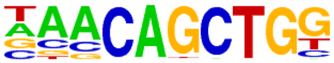 | Tcf21(bHLH)/ArterySmoothMuscle-Tcf21-ChIP-Seq(GSE61369)/Homer | 1e-3 | -7.740e+00 | 0.0016 | 248.0 | 12.66% | 4489.5 | 10.28% |
| 116 | | NPAS2(bHLH)/Liver-NPAS2-ChIP-Seq(GSE39860)/Homer | 1e-3 | -7.734e+00 | 0.0016 | 285.0 | 14.55% | 5245.2 | 12.02% |

| | | | | | | | | | |
|---|---|---|---|---|---|---|---|---|---|
| | 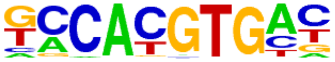 | | | | | | | | |
| 117 | 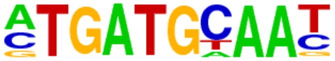 | Atf4(bZIP)/MEF-Atf4-ChIP-Seq(GSE35681)/Homer | 1e-3 | -7.676e+00 | 0.0017 | 63.0 | 3.22% | 896.2 | 2.05% |
| 118 | 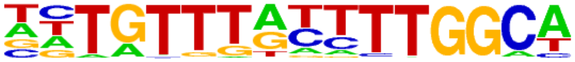 | NF1:FOXA1(CTF,Forkhead)/LNCAP-FOXA1-ChIP-Seq(GSE27824)/Homer | 1e-3 | -7.535e+00 | 0.0019 | 13.0 | 0.66% | 96.8 | 0.22% |
| 119 | 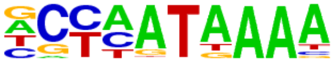 | Hoxd13(Homeobox)/ChickenMSG-Hoxd13.Flag-ChIP-Seq(GSE86088)/Homer | 1e-3 | -7.527e+00 | 0.0019 | 247.0 | 12.61% | 4486.2 | 10.28% |
| 120 | 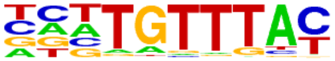 | FOXK1(Forkhead)/HEK293-FOXK1-ChIP-Seq(GSE51673)/Homer | 1e-3 | -7.349e+00 | 0.0023 | 170.0 | 8.68% | 2951.4 | 6.76% |
| 121 | 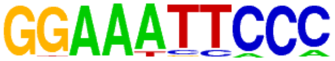 | NFkB-p65-Rel(RHD)/ThioMac-LPS-Expression(GSE23622)/Homer | 1e-3 | -7.191e+00 | 0.0027 | 19.0 | 0.97% | 181.5 | 0.42% |
| 122 | 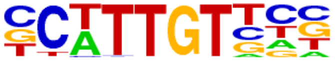 | Sox10(HMG)/SciaticNerve-Sox3-ChIP-Seq(GSE35132)/Homer | 1e-3 | -7.185e+00 | 0.0027 | 298.0 | 15.21% | 5562.7 | 12.74% |
| 123 | 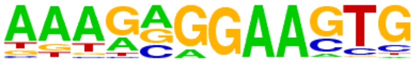 | SpiB(ETS)/OCILY3-SPIB-ChIP-Seq(GSE56857)/Homer | 1e-3 | -7.165e+00 | 0.0027 | 56.0 | 2.86% | 790.9 | 1.81% |
| 124 | 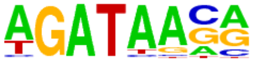 | GATA3(Zf)/iTreg-Gata3-ChIP-Seq(GSE20898)/Homer | 1e-3 | -7.041e+00 | 0.0030 | 235.0 | 12.00% | 4282.5 | 9.81% |
| 125 | 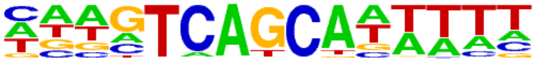 | MafF(bZIP)/HepG2-MafF-ChIP-Seq(GSE31477)/Homer | 1e-3 | -6.982e+00 | 0.0032 | 60.0 | 3.06% | 868.6 | 1.99% |
| 126 | 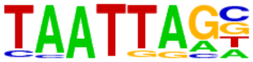 | Lhx2(Homeobox)/HFSC-Lhx2-ChIP-Seq(GSE48068)/Homer | 1e-2 | -6.859e+00 | 0.0036 | 179.0 | 9.14% | 3165.4 | 7.25% |
| 127 | 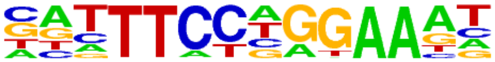 | STAT1(Stat)/HelaS3-STAT1-ChIP-Seq(GSE12782)/Homer | 1e-2 | -6.777e+00 | 0.0038 | 56.0 | 2.86% | 804.6 | 1.84% |
| 128 | 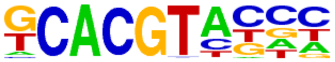 | HIF2a(bHLH)/785_O-HIF2a-ChIP-Seq(GSE34871)/Homer | 1e-2 | -6.562e+00 | 0.0047 | 87.0 | 4.44% | 1385.4 | 3.17% |
| 129 | 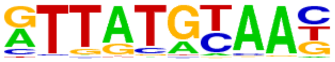 | HLF(bZIP)/HSC-HLF.Flag-ChIP-Seq(GSE69817)/Homer | 1e-2 | -6.481e+00 | 0.0051 | 128.0 | 6.53% | 2181.0 | 5.00% |
| 130 | 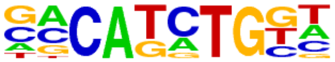 | TCF4(bHLH)/SHSY5Y-TCF4-ChIP-Seq(GSE96915)/Homer | 1e-2 | -6.466e+00 | 0.0051 | 378.0 | 19.30% | 7305.1 | 16.73% |
| 131 | 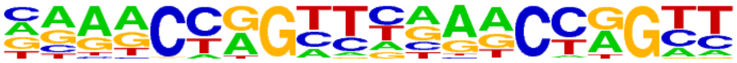 | Tcfcp2l1(CP2)/mES-Tcfcp2l1-ChIP-Seq(GSE11431)/Homer | 1e-2 | -6.447e+00 | 0.0052 | 35.0 | 1.79% | 449.5 | 1.03% |
| 132 | 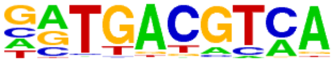 | Atf1(bZIP)/K562-ATF1-ChIP-Seq(GSE31477)/Homer | 1e-2 | -6.367e+00 | 0.0056 | 163.0 | 8.32% | 2881.7 | 6.60% |
| 133 | | HEB(bHLH)/mES-Heb-ChIP-Seq(GSE53233)/Homer | 1e-2 | -6.332e+00 | 0.0057 | 509.0 | 25.98% | 10104.5 | 23.15% |

| | | | | | | | | | |
|---|---|---|---|---|---|---|---|---|---|
| | 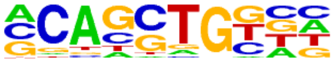 | | | | | | | | |
| 134 | 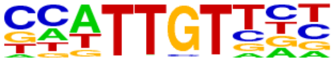 | Sox6(HMG)/Myotubes-Sox6-ChIP-Seq(GSE32627)/Homer | 1e-2 | -6.300e+00 | 0.0059 | 264.0 | 13.48% | 4943.8 | 11.33% |
| 135 | 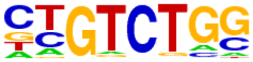 | Smad2(MAD)/ES-SMAD2-ChIP-Seq(GSE29422)/Homer | 1e-2 | -6.292e+00 | 0.0059 | 371.0 | 18.94% | 7177.3 | 16.44% |
| 136 | 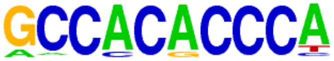 | Klf4(Zf)/mES-Klf4-ChIP-Seq(GSE11431)/Homer | 1e-2 | -6.228e+00 | 0.0062 | 87.0 | 4.44% | 1402.3 | 3.21% |
| 137 | 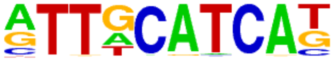 | Chop(bZIP)/MEF-Chop-ChIP-Seq(GSE35681)/Homer | 1e-2 | -6.223e+00 | 0.0062 | 51.0 | 2.60% | 735.8 | 1.69% |
| 138 | 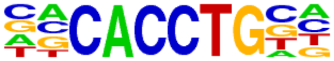 | E2A(bHLH),near_PU.1/Bcell-PU.1-ChIP-Seq(GSE21512)/Homer | 1e-2 | -6.217e+00 | 0.0062 | 423.0 | 21.59% | 8285.6 | 18.98% |
| 139 | 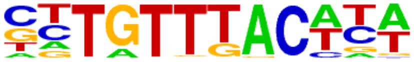 | Foxa2(Forkhead)/Liver-Foxa2-ChIP-Seq(GSE25694)/Homer | 1e-2 | -6.136e+00 | 0.0067 | 137.0 | 6.99% | 2380.4 | 5.45% |
| 140 | 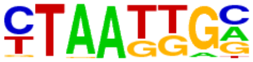 | Isl1(Homeobox)/Neuron-Isl1-ChIP-Seq(GSE31456)/Homer | 1e-2 | -6.134e+00 | 0.0067 | 337.0 | 17.20% | 6480.2 | 14.84% |
| 141 | 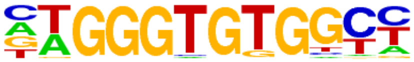 | EKLF(Zf)/Erythrocyte-Klf1-ChIP-Seq(GSE20478)/Homer | 1e-2 | -6.129e+00 | 0.0067 | 57.0 | 2.91% | 847.6 | 1.94% |
| 142 | 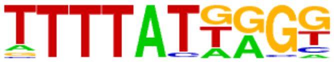 | HOXB13(Homeobox)/ProstateTumor-HOXB13-ChIP-Seq(GSE56288)/Homer | 1e-2 | -5.893e+00 | 0.0083 | 150.0 | 7.66% | 2656.6 | 6.09% |
| 143 | 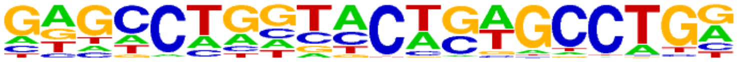 | ZNF322(Zf)/HEK293-ZNF322.GFP-ChIP-Seq(GSE58341)/Homer | 1e-2 | -5.839e+00 | 0.0087 | 89.0 | 4.54% | 1461.2 | 3.35% |
| 144 | 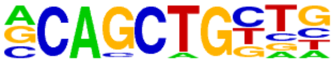 | Tcf12(bHLH)/GM12878-Tcf12-ChIP-Seq(GSE32465)/Homer | 1e-2 | -5.789e+00 | 0.0091 | 257.0 | 13.12% | 4848.7 | 11.11% |
| 145 | 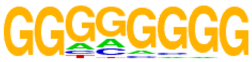 | Maz(Zf)/HepG2-Maz-ChIP-Seq(GSE31477)/Homer | 1e-2 | -5.674e+00 | 0.0101 | 255.0 | 13.02% | 4818.4 | 11.04% |
| 146 | 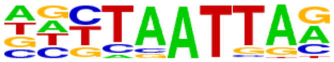 | Lhx1(Homeobox)/EmbryoCarcinoma-Lhx1-ChIP-Seq(GSE70957)/Homer | 1e-2 | -5.565e+00 | 0.0112 | 165.0 | 8.42% | 2983.6 | 6.83% |
| 147 | 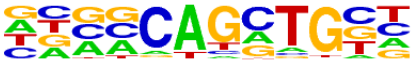 | Ascl1(bHLH)/NeuralTubes-Ascl1-ChIP-Seq(GSE55840)/Homer | 1e-2 | -5.525e+00 | 0.0116 | 405.0 | 20.67% | 7988.2 | 18.30% |
| 148 | 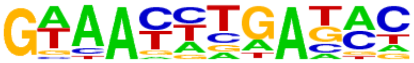 | Six2(Homeobox)/NephronProgenitor-Six2-ChIP-Seq(GSE39837)/Homer | 1e-2 | -5.464e+00 | 0.0123 | 168.0 | 8.58% | 3052.1 | 6.99% |
| 149 | 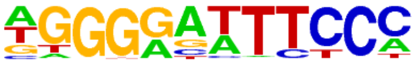 | NFkB-p65(RHD)/GM12787-p65-ChIP-Seq(GSE19485)/Homer | 1e-2 | -5.458e+00 | 0.0123 | 108.0 | 5.51% | 1853.3 | 4.25% |
| 150 | | Tlx?(NR)/NPC-H3K4me1-ChIP-Seq(GSE16256)/Homer | 1e-2 | -5.453e+00 | 0.0123 | 109.0 | 5.56% | 1873.9 | 4.29% |

| | | | | | | | | | |
|---|---|---|---|---|---|---|---|---|---|
| | 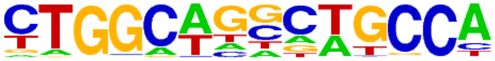 | | | | | | | | |
| 151 | 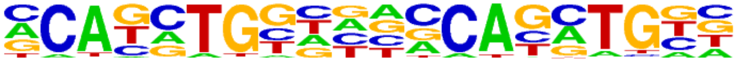 | Twist(bHLH)/HMLE-TWIST1-ChIP-Seq(Chang_et_al)/Homer | 1e-2 | -5.322e+00 | 0.0138 | 44.0 | 2.25% | 643.4 | 1.47% |
| 152 | 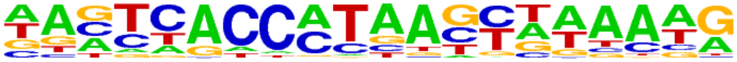 | PSE(SNAPc)/K562-mStart-Seq/Homer | 1e-2 | -5.169e+00 | 0.0160 | 90.0 | 4.59% | 1519.4 | 3.48% |
| 153 | 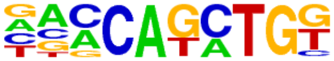 | BHLHA15(bHLH)/NIH3T3-BHLHB8.HA-ChIP-Seq(GSE119782)/Homer | 1e-2 | -5.075e+00 | 0.0175 | 333.0 | 17.00% | 6518.4 | 14.93% |
| 154 | 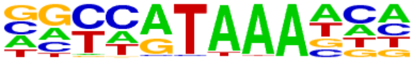 | CDX4(Homeobox)/ZebrafishEmbryos-Cdx4.Myc-ChIP-Seq(GSE48254)/Homer | 1e-2 | -4.989e+00 | 0.0189 | 125.0 | 6.38% | 2223.5 | 5.09% |
| 155 | 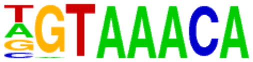 | Foxo3(Forkhead)/U2OS-Foxo3-ChIP-Seq(E-MTAB-2701)/Homer | 1e-2 | -4.969e+00 | 0.0192 | 113.0 | 5.77% | 1985.7 | 4.55% |
| 156 | 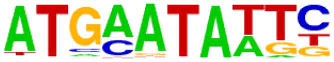 | Brn2(POU,Homeobox)/NPC-Brn2-ChIP-Seq(GSE35496)/Homer | 1e-2 | -4.967e+00 | 0.0192 | 21.0 | 1.07% | 256.0 | 0.59% |
| 157 | 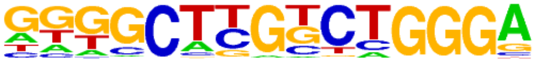 | Zfp809(Zf)/ES-Zfp809-ChIP-Seq(GSE70799)/Homer | 1e-2 | -4.898e+00 | 0.0203 | 57.0 | 2.91% | 901.6 | 2.07% |
| 158 | 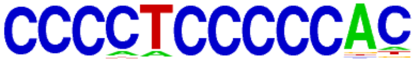 | Zfp281(Zf)/ES-Zfp281-ChIP-Seq(GSE81042)/Homer | 1e-2 | -4.842e+00 | 0.0214 | 37.0 | 1.89% | 535.5 | 1.23% |
| 159 | 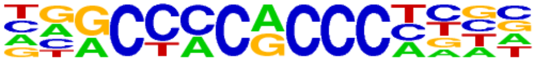 | KLF3(Zf)/MEF-Klf3-ChIP-Seq(GSE44748)/Homer | 1e-2 | -4.834e+00 | 0.0214 | 87.0 | 4.44% | 1481.1 | 3.39% |
| 160 | 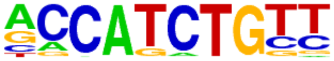 | NeuroG2(bHLH)/Fibroblast-NeuroG2-ChIP-Seq(GSE75910)/Homer | 1e-2 | -4.681e+00 | 0.0248 | 357.0 | 18.22% | 7077.8 | 16.21% |
| 161 | 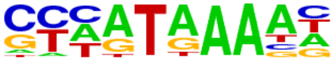 | Hoxa13(Homeobox)/ChickenMSG-Hoxa13.Flag-ChIP-Seq(GSE86088)/Homer | 1e-2 | -4.608e+00 | 0.0265 | 378.0 | 19.30% | 7535.0 | 17.26% |

### Homer Known Motif Enrichment Results (QC/homer/07_0938_SCC-13_JunB_hg38)

Homer *de novo* Motif Results
Gene Ontology Enrichment Results
Known Motif Enrichment Results (txt file)
Total Target Sequences = 1900, Total Background Sequences = 43424

| Rank | Motif | Name | P-value | log P-pvalue | q-value (Benjamini) | # Target Sequences with Motif | % of Targets Sequences with Motif | # Background Sequences with Motif | % of Background Sequences with Motif |
|---|---|---|---|---|---|---|---|---|---|
| 1 | 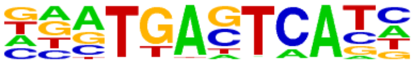 | Fra1(bZIP)/BT549-Fra1-ChIP-Seq(GSE46166)/Homer | 1e-1529 | -3.521e+03 | 0.0000 | 1561.0 | 82.16% | 2614.1 | 6.02% |
| 2 | 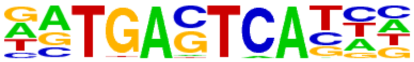 | Atf3(bZIP)/GBM-ATF3-ChIP-Seq(GSE33912)/Homer | 1e-1485 | -3.421e+03 | 0.0000 | 1608.0 | 84.63% | 3168.8 | 7.29% |
| 3 | 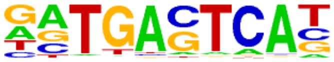 | JunB(bZIP)/DendriticCells-Junb-ChIP-Seq(GSE36099)/Homer | 1e-1476 | -3.401e+03 | 0.0000 | 1542.0 | 81.16% | 2683.9 | 6.18% |
| 4 | 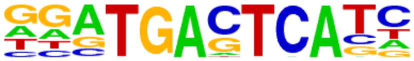 | Fra2(bZIP)/Striatum-Fra2-ChIP-Seq(GSE43429)/Homer | 1e-1476 | -3.400e+03 | 0.0000 | 1488.0 | 78.32% | 2308.7 | 5.31% |
| 5 | 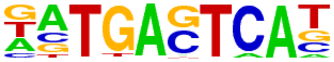 | BATF(bZIP)/Th17-BATF-ChIP-Seq(GSE39756)/Homer | 1e-1462 | -3.368e+03 | 0.0000 | 1592.0 | 83.79% | 3139.9 | 7.23% |
| 6 | 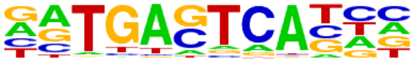 | Fosl2(bZIP)/3T3L1-Fosl2-ChIP-Seq(GSE56872)/Homer | 1e-1411 | -3.251e+03 | 0.0000 | 1326.0 | 69.79% | 1587.0 | 3.65% |
| 7 | 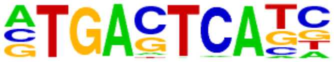 | AP-1(bZIP)/ThioMac-PU.1-ChIP-Seq(GSE21512)/Homer | 1e-1399 | -3.221e+03 | 0.0000 | 1614.0 | 84.95% | 3650.0 | 8.40% |
| 8 | 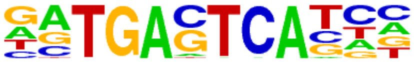 | Jun-AP1(bZIP)/K562-cJun-ChIP-Seq(GSE31477)/Homer | 1e-1278 | -2.945e+03 | 0.0000 | 1148.0 | 60.42% | 1123.9 | 2.59% |
| 9 | 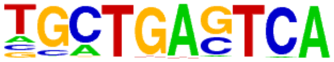 | Bach2(bZIP)/OCILy7-Bach2-ChIP-Seq(GSE44420)/Homer | 1e-661 | -1.523e+03 | 0.0000 | 716.0 | 37.68% | 931.0 | 2.14% |
| 10 | 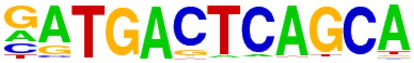 | NF-E2(bZIP)/K562-NFE2-ChIP-Seq(GSE31477)/Homer | 1e-261 | -6.012e+02 | 0.0000 | 258.0 | 13.58% | 238.8 | 0.55% |
| 11 | 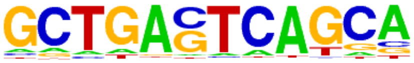 | MafK(bZIP)/C2C12-MafK-ChIP-Seq(GSE36030)/Homer | 1e-242 | -5.586e+02 | 0.0000 | 376.0 | 19.79% | 873.9 | 2.01% |
| 12 | 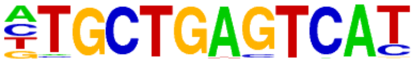 | Nrf2(bZIP)/Lymphoblast-Nrf2-ChIP-Seq(GSE37589)/Homer | 1e-218 | -5.023e+02 | 0.0000 | 213.0 | 11.21% | 190.0 | 0.44% |
| 13 | 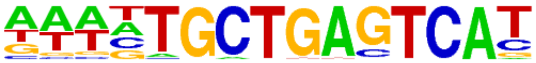 | Bach1(bZIP)/K562-Bach1-ChIP-Seq(GSE31477)/Homer | 1e-200 | -4.607e+02 | 0.0000 | 212.0 | 11.16% | 228.2 | 0.53% |
| 14 | | NFE2L2(bZIP)/HepG2-NFE2L2-ChIP-Seq(Encode)/Homer | 1e-144 | -3.332e+02 | 0.0000 | 158.0 | 8.32% | 180.3 | 0.41% |

| | | | | | | | | | |
|---|---|---|---|---|---|---|---|---|---|
| | 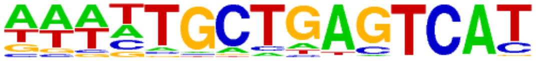 | | | | | | | | |
| 15 | 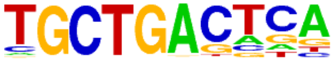 | MafA(bZIP)/Islet-MafA-ChIP-Seq(GSE30298)/Homer | 1e-109 | -2.514e+02 | 0.0000 | 487.0 | 25.63% | 3659.2 | 8.42% |
| 16 | 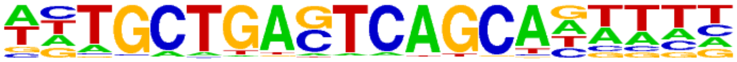 | MafB(bZIP)/BMM-Mafb-ChIP-Seq(GSE75722)/Homer | 1e-103 | -2.381e+02 | 0.0000 | 304.0 | 16.00% | 1558.7 | 3.59% |
| 17 | 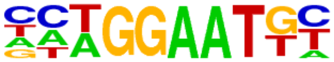 | TEAD(TEA)/Fibroblast-PU.1-ChIP-Seq(Unpublished)/Homer | 1e-38 | -8.931e+01 | 0.0000 | 260.0 | 13.68% | 2423.7 | 5.58% |
| 18 | 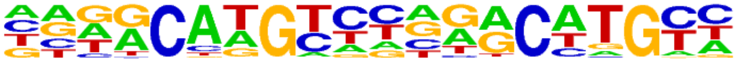 | p63(p53)/Keratinocyte-p63-ChIP-Seq(GSE17611)/Homer | 1e-33 | -7.816e+01 | 0.0000 | 167.0 | 8.79% | 1278.7 | 2.94% |
| 19 | 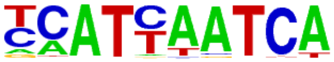 | Pdx1(Homeobox)/Islet-Pdx1-ChIP-Seq(SRA008281)/Homer | 1e-33 | -7.649e+01 | 0.0000 | 347.0 | 18.26% | 4024.1 | 9.26% |
| 20 | 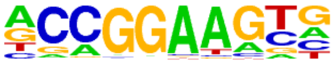 | ETV4(ETS)/HepG2-ETV4-ChIP-Seq(ENCODE)/Homer | 1e-33 | -7.640e+01 | 0.0000 | 340.0 | 17.89% | 3909.5 | 9.00% |
| 21 | 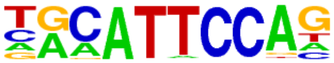 | TEAD3(TEA)/HepG2-TEAD3-ChIP-Seq(Encode)/Homer | 1e-32 | -7.592e+01 | 0.0000 | 367.0 | 19.32% | 4374.6 | 10.07% |
| 22 | 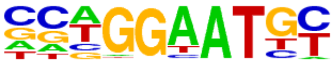 | TEAD4(TEA)/Tropoblast-Tead4-ChIP-Seq(GSE37350)/Homer | 1e-31 | -7.259e+01 | 0.0000 | 294.0 | 15.47% | 3232.4 | 7.44% |
| 23 | 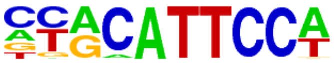 | TEAD1(TEAD)/HepG2-TEAD1-ChIP-Seq(Encode)/Homer | 1e-31 | -7.185e+01 | 0.0000 | 321.0 | 16.89% | 3689.2 | 8.49% |
| 24 | 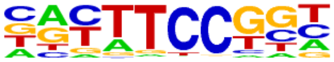 | Fli1(ETS)/CD8-FLI-ChIP-Seq(GSE20898)/Homer | 1e-30 | -7.071e+01 | 0.0000 | 332.0 | 17.47% | 3897.2 | 8.97% |
| 25 | 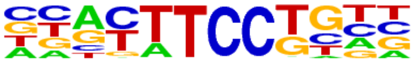 | Etv2(ETS)/ES-ER71-ChIP-Seq(GSE59402)/Homer | 1e-30 | -6.915e+01 | 0.0000 | 301.0 | 15.84% | 3415.7 | 7.86% |
| 26 | 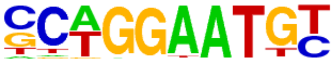 | TEAD2(TEA)/Py2T-Tead2-ChIP-Seq(GSE55709)/Homer | 1e-29 | -6.814e+01 | 0.0000 | 202.0 | 10.63% | 1891.9 | 4.35% |
| 27 | 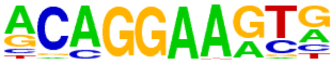 | ERG(ETS)/VCaP-ERG-ChIP-Seq(GSE14097)/Homer | 1e-29 | -6.803e+01 | 0.0000 | 458.0 | 24.11% | 6185.9 | 14.24% |
| 28 | 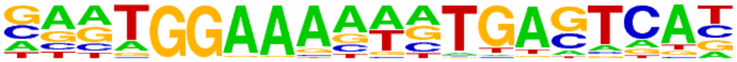 | NFAT:AP1(RHD,bZIP)/Jurkat-NFATC1-ChIP-Seq(Jolma_et_al.)/Homer | 1e-26 | -6.204e+01 | 0.0000 | 102.0 | 5.37% | 642.1 | 1.48% |
| 29 | 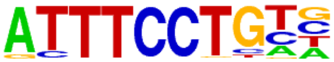 | EWS:ERG-fusion(ETS)/CADO_ES1-EWS:ERG-ChIP-Seq(SRA014231)/Homer | 1e-26 | -6.057e+01 | 0.0000 | 248.0 | 13.05% | 2728.8 | 6.28% |
| 30 | 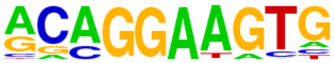 | ETS1(ETS)/Jurkat-ETS1-ChIP-Seq(GSE17954)/Homer | 1e-25 | -5.957e+01 | 0.0000 | 320.0 | 16.84% | 3949.8 | 9.09% |
| 31 | | ETV1(ETS)/GIST48-ETV1-ChIP-Seq(GSE22441)/Homer | 1e-23 | -5.503e+01 | 0.0000 | 378.0 | 19.89% | 5095.7 | 11.73% |

| | | | | | | | | | |
|---|---|---|---|---|---|---|---|---|---|
| | 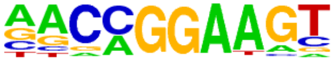 | | | | | | | | |
| 32 | 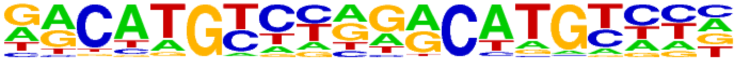 | p53(p53)/Saos-p53-ChIP-Seq(GSE15780)/Homer | 1e-22 | -5.193e+01 | 0.0000 | 66.0 | 3.47% | 332.7 | 0.77% |
| 33 | 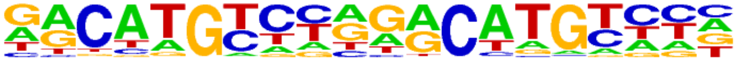 | p53(p53)/Saos-p53-ChIP-Seq/Homer | 1e-22 | -5.193e+01 | 0.0000 | 66.0 | 3.47% | 332.7 | 0.77% |
| 34 | 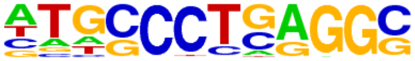 | AP-2alpha(AP2)/Hela-AP2alpha-ChIP-Seq(GSE31477)/Homer | 1e-22 | -5.122e+01 | 0.0000 | 251.0 | 13.21% | 2975.7 | 6.85% |
| 35 | 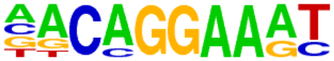 | EWS:FLI1-fusion(ETS)/SK_N_MC-EWS:FLI1-ChIP-Seq(SRA014231)/Homer | 1e-22 | -5.097e+01 | 0.0000 | 192.0 | 10.11% | 2028.0 | 4.67% |
| 36 | 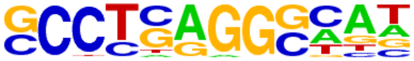 | AP-2gamma(AP2)/MCF7-TFAP2C-ChIP-Seq(GSE21234)/Homer | 1e-20 | -4.678e+01 | 0.0000 | 298.0 | 15.68% | 3895.1 | 8.97% |
| 37 | 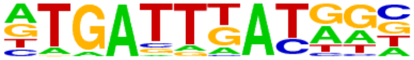 | PBX2(Homeobox)/K562-PBX2-ChIP-Seq(Encode)/Homer | 1e-19 | -4.575e+01 | 0.0000 | 259.0 | 13.63% | 3242.2 | 7.46% |
| 38 | 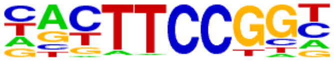 | Elk1(ETS)/Hela-Elk1-ChIP-Seq(GSE31477)/Homer | 1e-19 | -4.457e+01 | 0.0000 | 155.0 | 8.16% | 1574.8 | 3.62% |
| 39 | 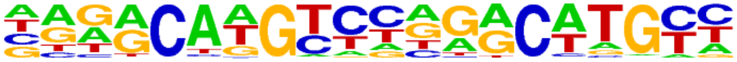 | p73(p53)/Trachea-p73-ChIP-Seq(PRJNA310161)/Homer | 1e-18 | -4.196e+01 | 0.0000 | 39.0 | 2.05% | 141.2 | 0.32% |
| 40 | 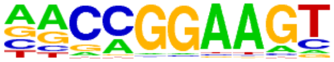 | GABPA(ETS)/Jurkat-GABPa-ChIP-Seq(GSE17954)/Homer | 1e-16 | -3.849e+01 | 0.0000 | 240.0 | 12.63% | 3104.2 | 7.15% |
| 41 | 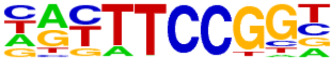 | Elk4(ETS)/Hela-Elk4-ChIP-Seq(GSE31477)/Homer | 1e-14 | -3.268e+01 | 0.0000 | 141.0 | 7.42% | 1583.9 | 3.65% |
| 42 | 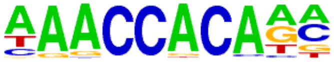 | RUNX1(Runt)/Jurkat-RUNX1-ChIP-Seq(GSE29180)/Homer | 1e-14 | -3.241e+01 | 0.0000 | 285.0 | 15.00% | 4102.7 | 9.44% |
| 43 | 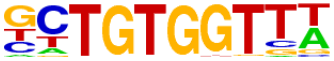 | RUNX-AML(Runt)/CD4+-PolII-ChIP-Seq(Barski_et_al.)/Homer | 1e-13 | -3.120e+01 | 0.0000 | 225.0 | 11.84% | 3050.8 | 7.02% |
| 44 | 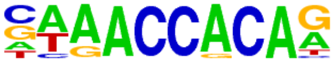 | RUNX(Runt)/HPC7-Runx1-ChIP-Seq(GSE22178)/Homer | 1e-13 | -3.012e+01 | 0.0000 | 218.0 | 11.47% | 2958.9 | 6.81% |
| 45 | 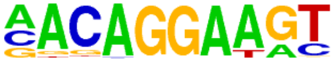 | Ets1-distal(ETS)/CD4+-PolII-ChIP-Seq(Barski_et_al.)/Homer | 1e-12 | -2.868e+01 | 0.0000 | 113.0 | 5.95% | 1222.9 | 2.81% |
| 46 | 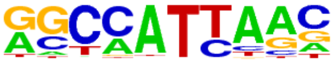 | Nanog(Homeobox)/mES-Nanog-ChIP-Seq(GSE11724)/Homer | 1e-12 | -2.818e+01 | 0.0000 | 929.0 | 48.89% | 17721.0 | 40.79% |
| 47 | 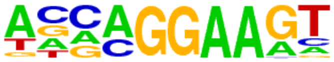 | EHF(ETS)/LoVo-EHF-ChIP-Seq(GSE49402)/Homer | 1e-12 | -2.778e+01 | 0.0000 | 311.0 | 16.37% | 4766.6 | 10.97% |
| 48 | | CEBP(bZIP)/ThioMac-CEBPb-ChIP-Seq(GSE21512)/Homer | 1e-11 | -2.721e+01 | 0.0000 | 157.0 | 8.26% | 1976.2 | 4.55% |

| | | | | | | | | | |
|---|---|---|---|---|---|---|---|---|---|
| | 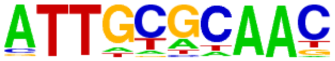 | | | | | | | | |
| 49 | 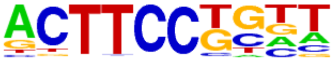 | Elf4(ETS)/BMDM-Elf4-ChIP-Seq(GSE88699)/Homer | 1e-11 | -2.704e+01 | 0.0000 | 254.0 | 13.37% | 3718.1 | 8.56% |
| 50 | 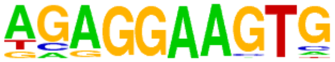 | PU.1(ETS)/ThioMac-PU.1-ChIP-Seq(GSE21512)/Homer | 1e-11 | -2.584e+01 | 0.0000 | 150.0 | 7.89% | 1893.9 | 4.36% |
| 51 | 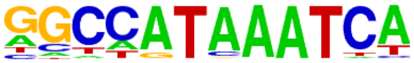 | Hoxc9(Homeobox)/Ainv15-Hoxc9-ChIP-Seq(GSE21812)/Homer | 1e-10 | -2.452e+01 | 0.0000 | 131.0 | 6.89% | 1606.1 | 3.70% |
| 52 | 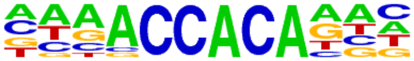 | RUNX2(Runt)/PCa-RUNX2-ChIP-Seq(GSE33889)/Homer | 1e-10 | -2.449e+01 | 0.0000 | 229.0 | 12.05% | 3347.6 | 7.71% |
| 53 | 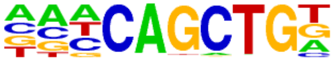 | Ap4(bHLH)/AML-Tfap4-ChIP-Seq(GSE45738)/Homer | 1e-10 | -2.380e+01 | 0.0000 | 331.0 | 17.42% | 5328.6 | 12.27% |
| 54 | 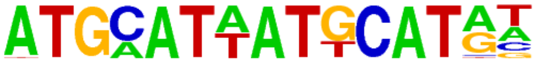 | OCT:OCT-short(POU,Homeobox)/NPC-OCT6-ChIP-Seq(GSE43916)/Homer | 1e-10 | -2.362e+01 | 0.0000 | 192.0 | 10.11% | 2699.2 | 6.21% |
| 55 | 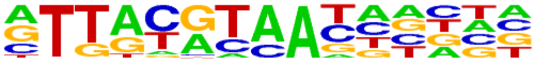 | NFIL3(bZIP)/HepG2-NFIL3-ChIP-Seq(Encode)/Homer | 1e-10 | -2.351e+01 | 0.0000 | 150.0 | 7.89% | 1957.7 | 4.51% |
| 56 | 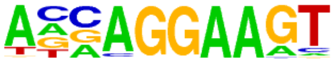 | ELF3(ETS)/PDAC-ELF3-ChIP-Seq(GSE64557)/Homer | 1e-10 | -2.309e+01 | 0.0000 | 191.0 | 10.05% | 2699.5 | 6.21% |
| 57 | 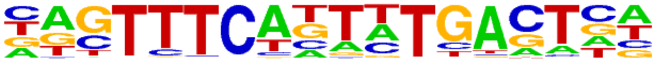 | bZIP:IRF(bZIP,IRF)/Th17-BatF-ChIP-Seq(GSE39756)/Homer | 1e-9 | -2.295e+01 | 0.0000 | 118.0 | 6.21% | 1428.7 | 3.29% |
| 58 | 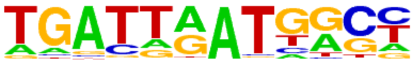 | Hoxb4(Homeobox)/ES-Hoxb4-ChIP-Seq(GSE34014)/Homer | 1e-9 | -2.220e+01 | 0.0000 | 77.0 | 4.05% | 791.0 | 1.82% |
| 59 | 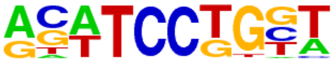 | SPDEF(ETS)/VCaP-SPDEF-ChIP-Seq(SRA014231)/Homer | 1e-8 | -2.008e+01 | 0.0000 | 248.0 | 13.05% | 3884.6 | 8.94% |
| 60 | 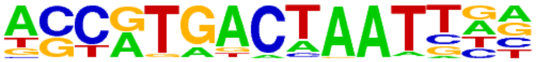 | PAX3:FKHR-fusion(Paired,Homeobox)/Rh4-PAX3:FKHR-ChIP-Seq(GSE19063)/Homer | 1e-8 | -1.884e+01 | 0.0000 | 68.0 | 3.58% | 716.9 | 1.65% |
| 61 | 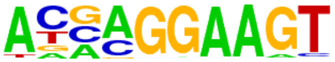 | ELF5(ETS)/T47D-ELF5-ChIP-Seq(GSE30407)/Homer | 1e-7 | -1.822e+01 | 0.0000 | 176.0 | 9.26% | 2599.6 | 5.98% |
| 62 | 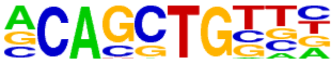 | Ptf1a(bHLH)/Panc1-Ptf1a-ChIP-Seq(GSE47459)/Homer | 1e-7 | -1.811e+01 | 0.0000 | 658.0 | 34.63% | 12483.2 | 28.73% |
| 63 | 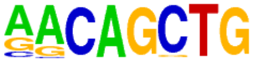 | MyoG(bHLH)/C2C12-MyoG-ChIP-Seq(GSE36024)/Homer | 1e-7 | -1.787e+01 | 0.0000 | 278.0 | 14.63% | 4574.5 | 10.53% |
| 64 | 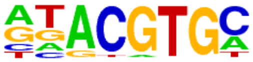 | HIF-1b(HLH)/T47D-HIF1b-ChIP-Seq(GSE59937)/Homer | 1e-7 | -1.783e+01 | 0.0000 | 247.0 | 13.00% | 3968.6 | 9.13% |
| 65 | | MITF(bHLH)/MastCells-MITF-ChIP-Seq(GSE48085)/Homer | 1e-7 | -1.753e+01 | 0.0000 | 246.0 | 12.95% | 3963.5 | 9.12% |

| | | | | | | | | | |
|---|---|---|---|---|---|---|---|---|---|
| | 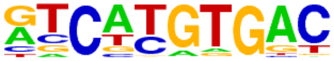 | | | | | | | | |
| 66 | 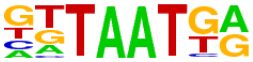 | Nkx6.1(Homeobox)/Islet-Nkx6.1-ChIP-Seq(GSE40975)/Homer | 1e-6 | -1.551e+01 | 0.0000 | 533.0 | 28.05% | 9995.1 | 23.01% |
| 67 | 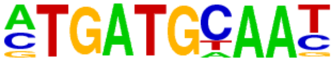 | Atf4(bZIP)/MEF-Atf4-ChIP-Seq(GSE35681)/Homer | 1e-6 | -1.482e+01 | 0.0000 | 81.0 | 4.26% | 1015.2 | 2.34% |
| 68 | 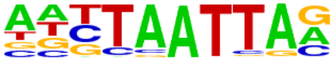 | Lhx3(Homeobox)/Neuron-Lhx3-ChIP-Seq(GSE31456)/Homer | 1e-6 | -1.479e+01 | 0.0000 | 334.0 | 17.58% | 5877.1 | 13.53% |
| 69 | 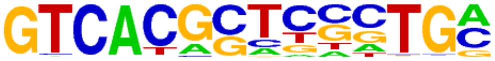 | PAX5(Paired,Homeobox),condensed/GM12878-PAX5-ChIP-Seq(GSE32465)/Homer | 1e-6 | -1.411e+01 | 0.0000 | 47.0 | 2.47% | 484.0 | 1.11% |
| 70 | 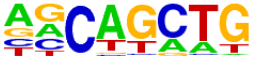 | SCL(bHLH)/HPC7-Scl-ChIP-Seq(GSE13511)/Homer | 1e-5 | -1.330e+01 | 0.0000 | 994.0 | 52.32% | 20402.5 | 46.96% |
| 71 | 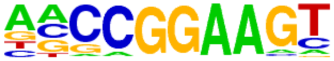 | ELF1(ETS)/Jurkat-ELF1-ChIP-Seq(SRA014231)/Homer | 1e-5 | -1.318e+01 | 0.0000 | 104.0 | 5.47% | 1466.3 | 3.38% |
| 72 | 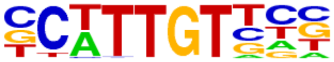 | Sox10(HMG)/SciaticNerve-Sox3-ChIP-Seq(GSE35132)/Homer | 1e-5 | -1.314e+01 | 0.0000 | 335.0 | 17.63% | 6007.9 | 13.83% |
| 73 | 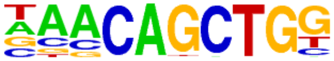 | Tcf21(bHLH)/ArterySmoothMuscle-Tcf21-ChIP-Seq(GSE61369)/Homer | 1e-5 | -1.278e+01 | 0.0000 | 234.0 | 12.32% | 3976.1 | 9.15% |
| 74 | 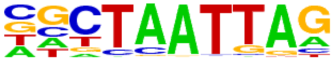 | LXH9(Homeobox)/Hct116-LXH9.V5-ChIP-Seq(GSE116822)/Homer | 1e-5 | -1.269e+01 | 0.0000 | 296.0 | 15.58% | 5237.0 | 12.05% |
| 75 | 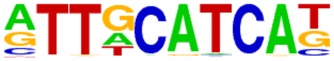 | Chop(bZIP)/MEF-Chop-ChIP-Seq(GSE35681)/Homer | 1e-5 | -1.262e+01 | 0.0000 | 64.0 | 3.37% | 787.6 | 1.81% |
| 76 | 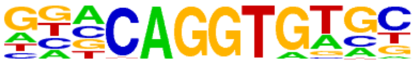 | ZEB2(Zf)/SNU398-ZEB2-ChIP-Seq(GSE103048)/Homer | 1e-5 | -1.245e+01 | 0.0000 | 274.0 | 14.42% | 4803.7 | 11.06% |
| 77 | 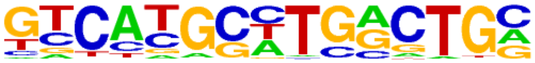 | Pax8(Paired,Homeobox)/Thyroid-Pax8-ChIP-Seq(GSE26938)/Homer | 1e-5 | -1.234e+01 | 0.0000 | 90.0 | 4.74% | 1244.8 | 2.87% |
| 78 | 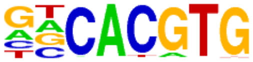 | BMAL1(bHLH)/Liver-Bmal1-ChIP-Seq(GSE39860)/Homer | 1e-5 | -1.203e+01 | 0.0000 | 446.0 | 23.47% | 8421.2 | 19.38% |
| 79 | 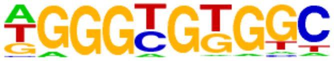 | KLF5(Zf)/LoVo-KLF5-ChIP-Seq(GSE49402)/Homer | 1e-5 | -1.180e+01 | 0.0000 | 240.0 | 12.63% | 4155.1 | 9.56% |
| 80 | 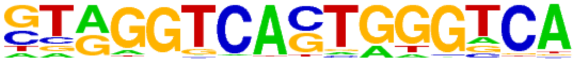 | Reverb(NR),DR2/RAW-Reverba.biotin-ChIP-Seq(GSE45914)/Homer | 1e-5 | -1.158e+01 | 0.0000 | 51.0 | 2.68% | 598.6 | 1.38% |
| 81 | 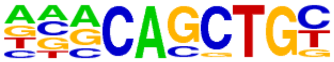 | E2A(bHLH)/proBcell-E2A-ChIP-Seq(GSE21978)/Homer | 1e-5 | -1.158e+01 | 0.0000 | 362.0 | 19.05% | 6683.9 | 15.38% |
| 82 | | Arnt:Ahr(bHLH)/MCF7-Arnt-ChIP-Seq(Lo_et_al.)/Homer | 1e-5 | -1.152e+01 | 0.0001 | 139.0 | 7.32% | 2182.1 | 5.02% |

| | | | | | | | | | |
|---|---|---|---|---|---|---|---|---|---|
| | 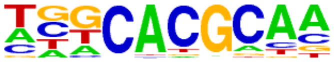 | | | | | | | | |
| 83 | 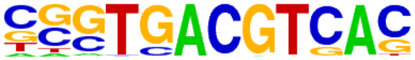 | CRE(bZIP)/Promoter/Homer | 1e-4 | -1.133e+01 | 0.0001 | 66.0 | 3.47% | 855.7 | 1.97% |
| 84 | 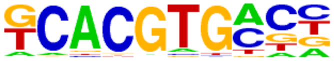 | bHLHE41(bHLH)/proB-Bhlhe41-ChIP-Seq(GSE93764)/Homer | 1e-4 | -1.108e+01 | 0.0001 | 233.0 | 12.26% | 4059.4 | 9.34% |
| 85 | 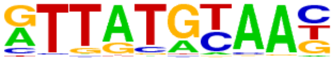 | HLF(bZIP)/HSC-HLF.Flag-ChIP-Seq(GSE69817)/Homer | 1e-4 | -1.093e+01 | 0.0001 | 159.0 | 8.37% | 2596.6 | 5.98% |
| 86 | 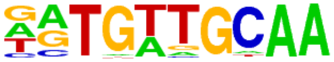 | CEBP:AP1(bZIP)/ThioMac-CEBPb-ChIP-Seq(GSE21512)/Homer | 1e-4 | -1.093e+01 | 0.0001 | 168.0 | 8.84% | 2772.7 | 6.38% |
| 87 | 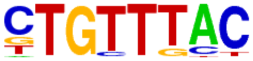 | Foxo1(Forkhead)/RAW-Foxo1-ChIP-Seq(Fan_et_al.)/Homer | 1e-4 | -1.088e+01 | 0.0001 | 371.0 | 19.53% | 6928.8 | 15.95% |
| 88 | 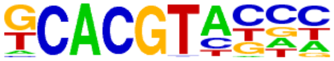 | HIF2a(bHLH)/785_O-HIF2a-ChIP-Seq(GSE34871)/Homer | 1e-4 | -1.054e+01 | 0.0001 | 88.0 | 4.63% | 1270.4 | 2.92% |
| 89 | 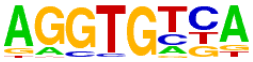 | Tbx5(T-box)/HL1-Tbx5.biotin-ChIP-Seq(GSE21529)/Homer | 1e-4 | -1.024e+01 | 0.0002 | 742.0 | 39.05% | 15055.0 | 34.65% |
| 90 | 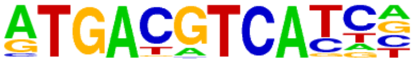 | JunD(bZIP)/K562-JunD-ChIP-Seq/Homer | 1e-4 | -1.021e+01 | 0.0002 | 30.0 | 1.58% | 298.2 | 0.69% |
| 91 | 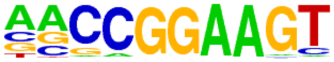 | ETS(ETS)/Promoter/Homer | 1e-4 | -1.005e+01 | 0.0002 | 68.0 | 3.58% | 928.2 | 2.14% |
| 92 | 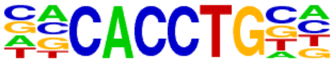 | E2A(bHLH),near_PU.1/Bcell-PU.1-ChIP-Seq(GSE21512)/Homer | 1e-4 | -9.887e+00 | 0.0002 | 373.0 | 19.63% | 7054.8 | 16.24% |
| 93 | 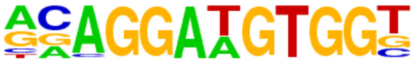 | ETS:RUNX(ETS,Runt)/Jurkat-RUNX1-ChIP-Seq(GSE17954)/Homer | 1e-4 | -9.868e+00 | 0.0002 | 32.0 | 1.68% | 334.2 | 0.77% |
| 94 | 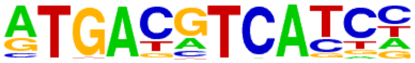 | c-Jun-CRE(bZIP)/K562-cJun-ChIP-Seq(GSE31477)/Homer | 1e-4 | -9.836e+00 | 0.0002 | 90.0 | 4.74% | 1333.6 | 3.07% |
| 95 | 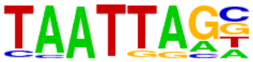 | Lhx2(Homeobox)/HFSC-Lhx2-ChIP-Seq(GSE48068)/Homer | 1e-4 | -9.653e+00 | 0.0003 | 204.0 | 10.74% | 3564.9 | 8.21% |
| 96 | 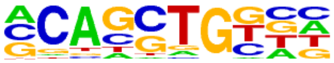 | HEB(bHLH)/mES-Heb-ChIP-Seq(GSE53233)/Homer | 1e-4 | -9.609e+00 | 0.0003 | 458.0 | 24.11% | 8896.4 | 20.48% |
| 97 | 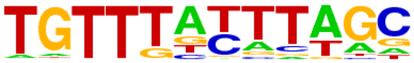 | FoxD3(forkhead)/ZebrafishEmbryo-Foxd3.biotin-ChIP-seq(GSE106676)/Homer | 1e-4 | -9.538e+00 | 0.0003 | 140.0 | 7.37% | 2298.2 | 5.29% |
| 98 | 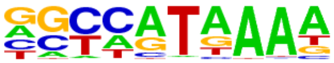 | Hoxd11(Homeobox)/ChickenMSG-Hoxd11.Flag-ChIP-Seq(GSE86088)/Homer | 1e-4 | -9.380e+00 | 0.0004 | 448.0 | 23.58% | 8703.3 | 20.03% |
| 99 | | Stat3+il21(Stat)/CD4-Stat3-ChIP-Seq(GSE19198)/Homer | 1e-4 | -9.243e+00 | 0.0004 | 139.0 | 7.32% | 2294.5 | 5.28% |

| | | | | | | | | | |
|---|---|---|---|---|---|---|---|---|---|
| | 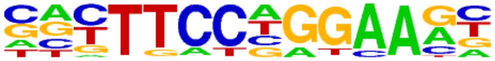 | | | | | | | | |
| 100 | 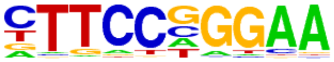 | Stat3(Stat)/mES-Stat3-ChIP-Seq(GSE11431)/Homer | 1e-4 | -9.241e+00 | 0.0004 | 100.0 | 5.26% | 1544.5 | 3.56% |
| 101 | 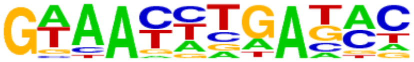 | Six2(Homeobox)/NephronProgenitor-Six2-ChIP-Seq(GSE39837)/Homer | 1e-3 | -9.099e+00 | 0.0005 | 195.0 | 10.26% | 3418.4 | 7.87% |
| 102 | 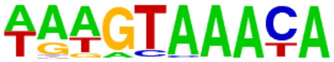 | FOXA1(Forkhead)/MCF7-FOXA1-ChIP-Seq(GSE26831)/Homer | 1e-3 | -9.056e+00 | 0.0005 | 169.0 | 8.89% | 2898.6 | 6.67% |
| 103 | 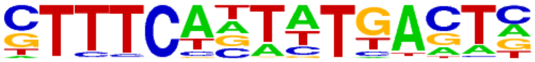 | IRF:BATF(IRF:bZIP)/pDC-Irf8-ChIP-Seq(GSE66899)/Homer | 1e-3 | -8.905e+00 | 0.0006 | 34.0 | 1.79% | 384.8 | 0.89% |
| 104 | 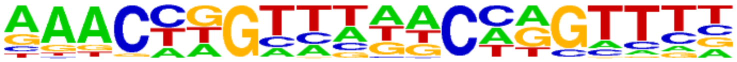 | GRHL2(CP2)/HBE-GRHL2-ChIP-Seq(GSE46194)/Homer | 1e-3 | -8.797e+00 | 0.0006 | 82.0 | 4.32% | 1226.8 | 2.82% |
| 105 | 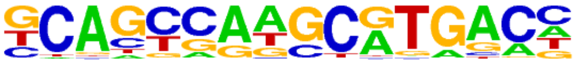 | PAX5(Paired,Homeobox)/GM12878-PAX5-ChIP-Seq(GSE32465)/Homer | 1e-3 | -8.676e+00 | 0.0007 | 91.0 | 4.79% | 1399.5 | 3.22% |
| 106 | 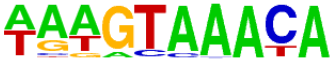 | FOXA1(Forkhead)/LNCAP-FOXA1-ChIP-Seq(GSE27824)/Homer | 1e-3 | -8.572e+00 | 0.0008 | 212.0 | 11.16% | 3800.5 | 8.75% |
| 107 | 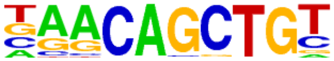 | Myf5(bHLH)/GM-Myf5-ChIP-Seq(GSE24852)/Homer | 1e-3 | -8.527e+00 | 0.0008 | 172.0 | 9.05% | 2991.6 | 6.89% |
| 108 | 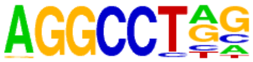 | ZFX(Zf)/mES-Zfx-ChIP-Seq(GSE11431)/Homer | 1e-3 | -8.500e+00 | 0.0008 | 277.0 | 14.58% | 5151.6 | 11.86% |
| 109 | 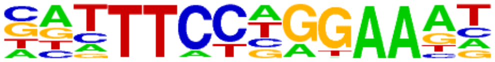 | STAT1(Stat)/HelaS3-STAT1-ChIP-Seq(GSE12782)/Homer | 1e-3 | -8.371e+00 | 0.0009 | 59.0 | 3.11% | 823.3 | 1.90% |
| 110 | 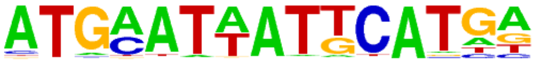 | OCT:OCT(POU,Homeobox,IR1)/NPC-Brn2-ChIP-Seq(GSE35496)/Homer | 1e-3 | -8.315e+00 | 0.0010 | 9.0 | 0.47% | 46.5 | 0.11% |
| 111 | 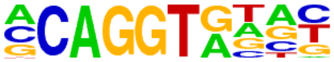 | ZEB1(Zf)/PDAC-ZEB1-ChIP-Seq(GSE64557)/Homer | 1e-3 | -8.307e+00 | 0.0010 | 421.0 | 22.16% | 8229.6 | 18.94% |
| 112 | 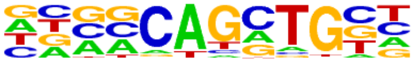 | Ascl1(bHLH)/NeuralTubes-Ascl1-ChIP-Seq(GSE55840)/Homer | 1e-3 | -8.090e+00 | 0.0012 | 362.0 | 19.05% | 6985.7 | 16.08% |
| 113 | 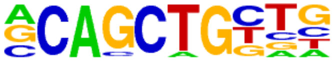 | Tcf12(bHLH)/GM12878-Tcf12-ChIP-Seq(GSE32465)/Homer | 1e-3 | -7.994e+00 | 0.0013 | 229.0 | 12.05% | 4193.3 | 9.65% |
| 114 | 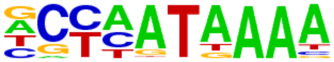 | Hoxd13(Homeobox)/ChickenMSG-Hoxd13.Flag-ChIP-Seq(GSE86088)/Homer | 1e-3 | -7.886e+00 | 0.0014 | 284.0 | 14.95% | 5350.7 | 12.32% |
| 115 | 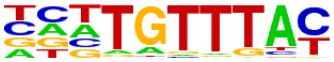 | FOXK1(Forkhead)/HEK293-FOXK1-ChIP-Seq(GSE51673)/Homer | 1e-3 | -7.698e+00 | 0.0017 | 201.0 | 10.58% | 3638.8 | 8.38% |
| 116 | | STAT4(Stat)/CD4-Stat4-ChIP-Seq(GSE22104)/Homer | 1e-3 | -7.648e+00 | 0.0018 | 175.0 | 9.21% | 3110.6 | 7.16% |

| | | | | | | | | | |
|---|---|---|---|---|---|---|---|---|---|
| | 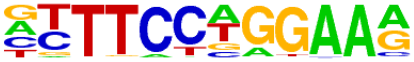 | | | | | | | | |
| 117 | 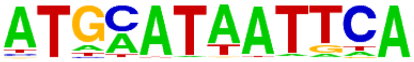 | Pit1+1bp(Homeobox)/GCrat-Pit1-ChIP-Seq(GSE58009)/Homer | 1e-3 | -7.521e+00 | 0.0020 | 67.0 | 3.53% | 999.6 | 2.30% |
| 118 | 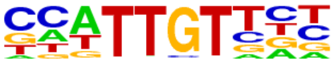 | Sox6(HMG)/Myotubes-Sox6-ChIP-Seq(GSE32627)/Homer | 1e-3 | -7.504e+00 | 0.0020 | 292.0 | 15.37% | 5553.7 | 12.78% |
| 119 | 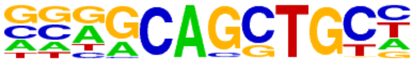 | Ascl2(bHLH)/ESC-Ascl2-ChIP-Seq(GSE97712)/Homer | 1e-3 | -7.437e+00 | 0.0021 | 267.0 | 14.05% | 5032.5 | 11.58% |
| 120 | 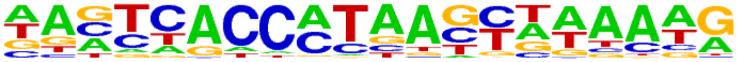 | PSE(SNAPc)/K562-mStart-Seq/Homer | 1e-3 | -7.402e+00 | 0.0022 | 101.0 | 5.32% | 1650.2 | 3.80% |
| 121 | 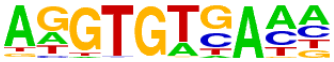 | Tbet(T-box)/CD8-Tbet-ChIP-Seq(GSE33802)/Homer | 1e-3 | -7.271e+00 | 0.0025 | 270.0 | 14.21% | 5110.7 | 11.76% |
| 122 | 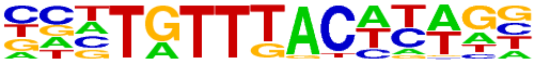 | Foxa3(Forkhead)/Liver-Foxa3-ChIP-Seq(GSE77670)/Homer | 1e-3 | -7.235e+00 | 0.0025 | 68.0 | 3.58% | 1029.3 | 2.37% |
| 123 | 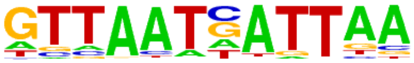 | HNF1b(Homeobox)/PDAC-HNF1B-ChIP-Seq(GSE64557)/Homer | 1e-3 | -7.101e+00 | 0.0029 | 31.0 | 1.63% | 376.3 | 0.87% |
| 124 | 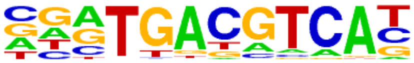 | Atf7(bZIP)/3T3L1-Atf7-ChIP-Seq(GSE56872)/Homer | 1e-3 | -7.092e+00 | 0.0029 | 120.0 | 6.32% | 2040.4 | 4.70% |
| 125 | 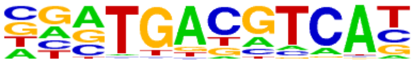 | Atf2(bZIP)/3T3L1-Atf2-ChIP-Seq(GSE56872)/Homer | 1e-3 | -7.068e+00 | 0.0029 | 90.0 | 4.74% | 1454.8 | 3.35% |
| 126 | 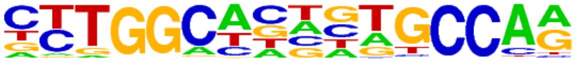 | NF1(CTF)/LNCAP-NF1-ChIP-Seq(Unpublished)/Homer | 1e-3 | -6.988e+00 | 0.0031 | 90.0 | 4.74% | 1458.6 | 3.36% |
| 127 | 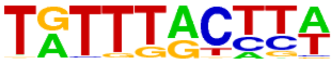 | FOXM1(Forkhead)/MCF7-FOXM1-ChIP-Seq(GSE72977)/Homer | 1e-3 | -6.962e+00 | 0.0032 | 175.0 | 9.21% | 3159.1 | 7.27% |
| 128 | 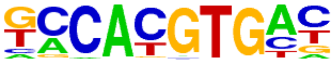 | NPAS2(bHLH)/Liver-NPAS2-ChIP-Seq(GSE39860)/Homer | 1e-3 | -6.951e+00 | 0.0032 | 255.0 | 13.42% | 4823.1 | 11.10% |
| 129 | 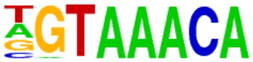 | Foxo3(Forkhead)/U2OS-Foxo3-ChIP-Seq(E-MTAB-2701)/Homer | 1e-2 | -6.866e+00 | 0.0035 | 140.0 | 7.37% | 2454.4 | 5.65% |
| 130 | 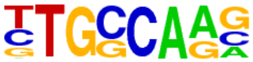 | NF1-halfsite(CTF)/LNCaP-NF1-ChIP-Seq(Unpublished)/Homer | 1e-2 | -6.827e+00 | 0.0036 | 378.0 | 19.89% | 7459.1 | 17.17% |
| 131 | 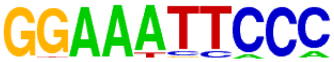 | NFkB-p65-Rel(RHD)/ThioMac-LPS-Expression(GSE23622)/Homer | 1e-2 | -6.761e+00 | 0.0038 | 18.0 | 0.95% | 178.2 | 0.41% |
| 132 | 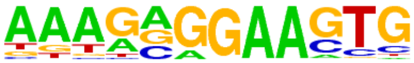 | SpiB(ETS)/OCILY3-SPIB-ChIP-Seq(GSE56857)/Homer | 1e-2 | -6.747e+00 | 0.0038 | 51.0 | 2.68% | 735.1 | 1.69% |
| 133 | | HOXB13(Homeobox)/ProstateTumor-HOXB13-ChIP-Seq(GSE56288)/Homer | 1e-2 | -6.358e+00 | 0.0056 | 172.0 | 9.05% | 3143.1 | 7.23% |

| | | | | | | | | | |
|---|---|---|---|---|---|---|---|---|---|
| | 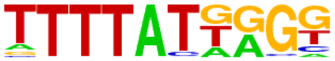 | | | | | | | | |
| 134 | 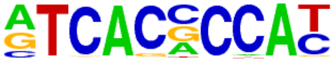 | Srebp1a(bHLH)/HepG2-Srebp1a-ChIP-Seq(GSE31477)/Homer | 1e-2 | -6.219e+00 | 0.0064 | 64.0 | 3.37% | 997.4 | 2.30% |
| 135 | 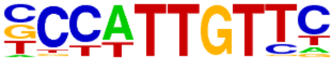 | Sox2(HMG)/mES-Sox2-ChIP-Seq(GSE11431)/Homer | 1e-2 | -6.181e+00 | 0.0066 | 157.0 | 8.26% | 2848.9 | 6.56% |
| 136 | 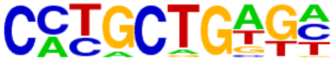 | Zic(Zf)/Cerebellum-ZIC1.2-ChIP-Seq(GSE60731)/Homer | 1e-2 | -6.177e+00 | 0.0066 | 191.0 | 10.05% | 3551.4 | 8.17% |
| 137 | 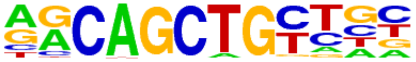 | MyoD(bHLH)/Myotube-MyoD-ChIP-Seq(GSE21614)/Homer | 1e-2 | -5.930e+00 | 0.0083 | 172.0 | 9.05% | 3177.1 | 7.31% |
| 138 | 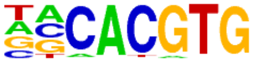 | NPAS(bHLH)/Liver-NPAS-ChIP-Seq(GSE39860)/Homer | 1e-2 | -5.923e+00 | 0.0083 | 360.0 | 18.95% | 7173.0 | 16.51% |
| 139 | 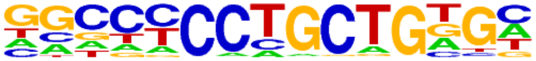 | Zic3(Zf)/mES-Zic3-ChIP-Seq(GSE37889)/Homer | 1e-2 | -5.896e+00 | 0.0085 | 132.0 | 6.95% | 2358.0 | 5.43% |
| 140 | 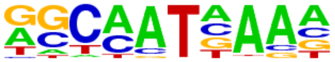 | Hoxa9(Homeobox)/ChickenMSG-Hoxa9.Flag-ChIP-Seq(GSE86088)/Homer | 1e-2 | -5.892e+00 | 0.0085 | 527.0 | 27.74% | 10830.4 | 24.93% |
| 141 | 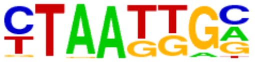 | Isl1(Homeobox)/Neuron-Isl1-ChIP-Seq(GSE31456)/Homer | 1e-2 | -5.841e+00 | 0.0088 | 362.0 | 19.05% | 7226.8 | 16.63% |
| 142 | 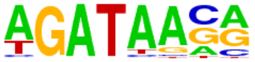 | GATA3(Zf)/iTreg-Gata3-ChIP-Seq(GSE20898)/Homer | 1e-2 | -5.789e+00 | 0.0092 | 259.0 | 13.63% | 5019.3 | 11.55% |
| 143 | 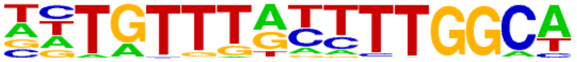 | NF1:FOXA1(CTF,Forkhead)/LNCAP-FOXA1-ChIP-Seq(GSE27824)/Homer | 1e-2 | -5.662e+00 | 0.0104 | 12.0 | 0.63% | 108.7 | 0.25% |
| 144 | 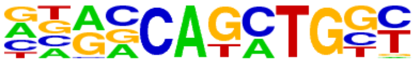 | Atoh1(bHLH)/Cerebellum-Atoh1-ChIP-Seq(GSE22111)/Homer | 1e-2 | -5.660e+00 | 0.0104 | 243.0 | 12.79% | 4692.3 | 10.80% |
| 145 | 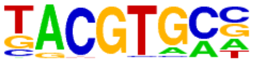 | HIF-1a(bHLH)/MCF7-HIF1a-ChIP-Seq(GSE28352)/Homer | 1e-2 | -5.626e+00 | 0.0106 | 55.0 | 2.89% | 853.2 | 1.96% |
| 146 | 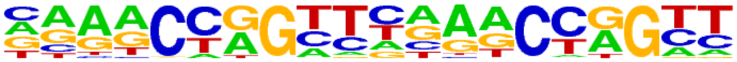 | Tcfcp2l1(CP2)/mES-Tcfcp2l1-ChIP-Seq(GSE11431)/Homer | 1e-2 | -5.615e+00 | 0.0107 | 32.0 | 1.68% | 433.3 | 1.00% |
| 147 | 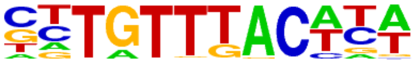 | Foxa2(Forkhead)/Liver-Foxa2-ChIP-Seq(GSE25694)/Homer | 1e-2 | -5.404e+00 | 0.0131 | 154.0 | 8.11% | 2847.7 | 6.55% |
| 148 | 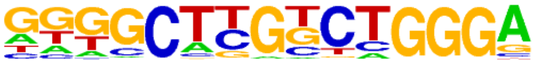 | Zfp809(Zf)/ES-Zfp809-ChIP-Seq(GSE70799)/Homer | 1e-2 | -5.345e+00 | 0.0138 | 44.0 | 2.32% | 659.9 | 1.52% |
| 149 | 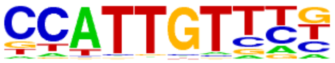 | Sox17(HMG)/Endoderm-Sox17-ChIP-Seq(GSE61475)/Homer | 1e-2 | -5.220e+00 | 0.0155 | 136.0 | 7.16% | 2490.6 | 5.73% |
| 150 | | Six1(Homeobox)/Myoblast-Six1-ChIP-Chip(GSE20150)/Homer | 1e-2 | -5.089e+00 | 0.0176 | 54.0 | 2.84% | 858.5 | 1.98% |

| | | | | | | | | | |
|---|---|---|---|---|---|---|---|---|---|
| | 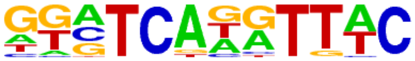 | | | | | | | | |
| 151 | 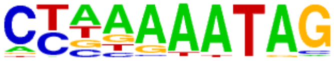 | Mef2a(MADS)/HL1-Mef2a.biotin-ChIP-Seq(GSE21529)/Homer | 1e-2 | -5.032e+00 | 0.0185 | 65.0 | 3.42% | 1073.3 | 2.47% |
| 152 | 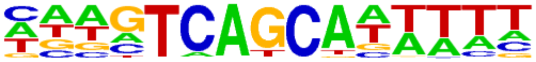 | MafF(bZIP)/HepG2-MafF-ChIP-Seq(GSE31477)/Homer | 1e-2 | -5.005e+00 | 0.0189 | 59.0 | 3.11% | 959.0 | 2.21% |
| 153 | 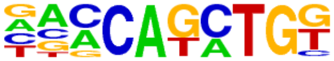 | BHLHA15(bHLH)/NIH3T3-BHLHB8.HA-ChIP-Seq(GSE119782)/Homer | 1e-2 | -4.974e+00 | 0.0194 | 305.0 | 16.05% | 6097.7 | 14.04% |
| 154 | 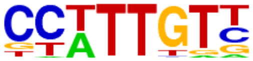 | Sox3(HMG)/NPC-Sox3-ChIP-Seq(GSE33059)/Homer | 1e-2 | -4.924e+00 | 0.0202 | 308.0 | 16.21% | 6168.2 | 14.20% |
| 155 | 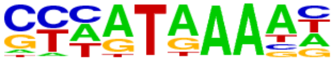 | Hoxa13(Homeobox)/ChickenMSG-Hoxa13.Flag-ChIP-Seq(GSE86088)/Homer | 1e-2 | -4.832e+00 | 0.0220 | 432.0 | 22.74% | 8887.1 | 20.46% |
| 156 | 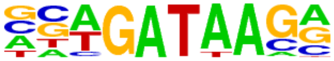 | Gata4(Zf)/Heart-Gata4-ChIP-Seq(GSE35151)/Homer | 1e-2 | -4.773e+00 | 0.0232 | 174.0 | 9.16% | 3320.7 | 7.64% |
| 157 | 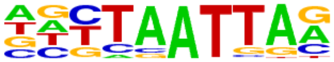 | Lhx1(Homeobox)/EmbryoCarcinoma-Lhx1-ChIP-Seq(GSE70957)/Homer | 1e-2 | -4.772e+00 | 0.0232 | 185.0 | 9.74% | 3552.6 | 8.18% |
| 158 | 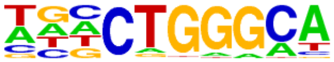 | ZNF416(Zf)/HEK293-ZNF416.GFP-ChIP-Seq(GSE58341)/Homer | 1e-2 | -4.678e+00 | 0.0252 | 275.0 | 14.47% | 5485.8 | 12.63% |
| 159 | 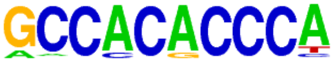 | Klf4(Zf)/mES-Klf4-ChIP-Seq(GSE11431)/Homer | 1e-2 | -4.643e+00 | 0.0259 | 62.0 | 3.26% | 1035.1 | 2.38% |
